# Tissue-Specific Dependence of Th1 Cells on the Amino Acid Transporter SLC38A1 in Inflammation

**DOI:** 10.1101/2023.09.13.557496

**Authors:** Ayaka Sugiura, Katherine L. Beier, Channing Chi, Darren R. Heintzman, Xiang Ye, Melissa M. Wolf, Andrew R. Patterson, Jacqueline-Yvonne Cephus, Hanna S. Hong, Costas A. Lyssiotis, Dawn C. Newcomb, Jeffrey C. Rathmell

**Affiliations:** Department of Pathology, Microbiology, and Immunology, Vanderbilt University Medical Center, Nashville, TN 37232, USA; Department of Medicine, Division of Hematology and Oncology, Vanderbilt University Medical Center, Nashville, TN 37232, USA; Department of Medicine, Division of Pulmonary and Critical Care, Vanderbilt University Medical Center, Nashville, TN 37232, USA; Rogel Cancer Center, University of Michigan, Ann Arbor, MI 48109 USA; Department of Molecular and Integrative Physiology, University of Michigan, Ann Arbor, MI 48109 USA; Department of Internal Medicine, Division of Gastroenterology and Hepatology, University of Michigan, Ann Arbor, MI 48109 USA; Vanderbilt Center for Immunobiology, Vanderbilt University Medical Center, Nashville, TN 37232, USA

**Keywords:** T cell, amino acid transport, Slc38a1, glutaminolysis

## Abstract

Amino acid (AA) uptake is essential for T cell metabolism and function, but how tissue sites and inflammation affect CD4^+^ T cell subset requirements for specific AA remains uncertain. Here we tested CD4^+^ T cell AA demands with *in vitro* and multiple *in vivo* CRISPR screens and identify subset- and tissue-specific dependencies on the AA transporter SLC38A1 (SNAT1). While dispensable for T cell persistence and expansion over time *in vitro* and *in vivo* lung inflammation, SLC38A1 was critical for Th1 but not Th17 cell-driven Experimental Autoimmune Encephalomyelitis (EAE) and contributed to Th1 cell-driven inflammatory bowel disease. SLC38A1 deficiency reduced mTORC1 signaling and glycolytic activity in Th1 cells, in part by reducing intracellular glutamine and disrupting hexosamine biosynthesis and redox regulation. Similarly, pharmacological inhibition of SLC38 transporters delayed EAE but did not affect lung inflammation. Subset- and tissue-specific dependencies of CD4^+^ T cells on AA transporters may guide selective immunotherapies.

**HIGHLIGHTS:**

- T cells dynamically regulate glutamine amino acid transporters when activated
- SLC38A1 supports Th1 cell mTORC1 and proliferation by redox and hexosamine pathways
- Targeting SLC38A1 does not affect lung inflammation but delays IBD and EAE
- Nutrient transporter needs of T cell subsets vary based on disease and tissue site

## INTRODUCTION

An effective immune response requires appropriate T cell activation, proliferation, differentiation, and function. Each of these processes utilizes specific cellular metabolic programs with distinct nutrient requirements. In addition to glucose and lipids, coordination of appropriate amino acid (AA) uptake and metabolism is critical to CD4^+^ T cell activity ^1,2^. While essential AAs must be imported from the extracellular environment, non-essential AAs can be synthesized intracellularly. Cells may also become dependent on uptake of conditionally essential AAs from extracellular pools when synthesis fails to meet increased demands, such as may occur in rapid cell growth and proliferation following T-cell receptor (TCR) stimulation. Previous studies have shown that CD4^+^ T cell fate and function may be altered or impaired in the setting of specific AA insufficiencies, including glutamine ^3–9^, leucine ^10,11^, arginine ^12,13^, serine ^14^, alanine ^15^, and methionine ^16–18^. Of these, the conditionally-essential AA glutamine has been most clearly shown to differentially affect both effector and regulatory CD4^+^ T helper (Teff and Treg) cell subsets ^8,19,20^.

AA transport is mediated by the solute carrier (SLC) family of nutrient transporters. The SLC family is comprised of over 400 facilitative and secondary active membrane transporters that carry a variety of organic and inorganic substrates and includes approximately 60 family members that selectively transport AAs. With more than 100 monogenic diseases linked to SLCs to date, these transporters have attracted interest as potential therapeutic targets ^21,22^. The breadth of transporters with overlapping substrates provides transport redundancy and opportunities to fine-tune nutrient use and metabolism by expression of different transporter isoforms. Previous studies have shown that SLC7A5 (LAT1) and SLC1A5 (ASCT2) play critical roles affecting inflammatory functions in CD4^+^ Teff cells ^4,11^. SLC7A5 forms a heterodimer with the heavy chain SLC3A2 (CD98) and imports Leu and other large neutral AAs in exchange for glutamine. The intracellular glutamine pool is in turn primarily maintained by SLC1A5-mediated uptake and is essential to drive transport of essential AAs and support a wide array of metabolic pathways and regulatory mechanisms. Loss of SLC7A5 is associated with failure in mTORC1 and MYC signaling-driven metabolic reprogramming leading to impaired clonal expansion and Teff cell differentiation following TCR engagement ^11^. Similarly, loss of SLC1A5 is associated with mTORC1-mediated impairment in Th1 and Th17 cell differentiation and function ^4^. Despite these clear roles for SLC1A5, T cells express multiple glutamine transporters that may also contribute to T cell metabolism in specific settings.

How AA transport mechanisms and requirements of distinct CD4^+^ T cell subsets are shaped by specific tissue microenvironments remain poorly understood. Differential gene expression and transport redundancy of SLC family members together with variable nutrient access in distinct tissues suggests that individual nutrient transporters may have context-specific phenotypes. To test this, we assessed AA requirements and focused on the conditionally essential AA glutamine *in vitro* and across multiple disease models of autoimmunity and inflammation. Glutamine was required by Teff, but not by Treg *in vitro*. CRISPR screening of glutamine transporters and metabolizing enzymes showed that while dispensable *in vitro*, the sodium-coupled neutral AA transporter SLC38A1 (SNAT1) was specifically required *in vivo* in a model of Experimental Autoimmune Encephalomyelitis (EAE) in pathogenic Th1 but not Th17 cells. SLC38A1-inhibition or deficiency delayed EAE symptom onset and lessened weight loss associated with Th1-cell driven inflammatory bowel disease (IBD). SLC38A1 was dispensable, however, in a T-cell driven model of allergic airway disease. These data show cell subset- and tissue-specific AA transport dependencies that illustrate the role of the tissue microenvironment on T cell metabolism and provide a new mechanism for selectivity in potential immunotherapy targets.

## RESULTS

### CD4^+^ T cells are dependent on uptake of select AAs to support activation, proliferation, and differentiation

Previous studies have shown that CD4^+^ T cells are dependent on the uptake of select essential and conditionally essential AAs ^23^. To extend these studies, we first sought to evaluate relative AA dependencies during T cell activation, proliferation, and differentiation by culturing cells in complete media replete of all AAs equivalent to the commercial formulation of RPMI-1640, and 19 additional media each lacking one AA and replete of all others. Since the commercial RPMI-1640 formulation does not contain alanine, alanine-subtracted medium was not included. Although the final culture media were supplemented with 10% fetal bovine serum (FBS) containing a rich variety of nutrients including AAs, nuclear magnetic resonance (NMR) analysis of the media confirmed that significant reduction was achieved in each of the AA-subtracted media relative to complete media (representative NMR spectra for glutamine shown in **Figure S1A,** relative abundancies of other AAs shown in **Figure S1B**).

Primary murine CD4^+^ T cell were activated with anti-CD3 and anti-CD28 antibodies in the indicated media and cultured for 72 hours (Th0 cells). Of the 19 AAs tested, histidine, isoleucine, leucine, lysine, methionine, phenylalanine, threonine, tryptophan, and valine are essential and cannot be *de novo* synthesized. CD4^+^ T cells activated in media deficient in any one of these essential AAs generally exhibited impaired activation, proliferation, and survival as expected (**Figure 1A-E**). Notably, subtraction of several conditionally essential and non-essential AAs also impaired Th0 cell activation and proliferation, including arginine, asparagine, cystine, glutamine, and tyrosine. In contrast, subtraction of aspartate, glutamate, glycine, proline, or serine did not significantly affect any of the measured parameters (**Figure 1A-E**). These data showing T cell dependence on uptake of extracellular arginine, glutamine, leucine, and methionine are consistent with published literature ^3–6,10,12,17,7^. Next, the experiment was repeated in the presence of either IL-6, TGFß, IL-1ß, and IL23 or TGFß and IL-2 to promote Th17 cell or Treg cell differentiation. Th17 cells generally showed a similar pattern of AA dependencies as Th0 cells (**Figure 1F**). Lysine, methionine, threonine, glutamine, and asparagine-deficient media each resulted in statistically significant impairment in upregulation of the transcription factor RORγt that is characteristic of the Th17 cell lineage (**Figure 1F**). Treg cells displayed similar requirements for the same set of AAs for FoxP3 upregulation, with the notable exception of not relying on glutamine (**Figure 1G**).

**Figure 1:**
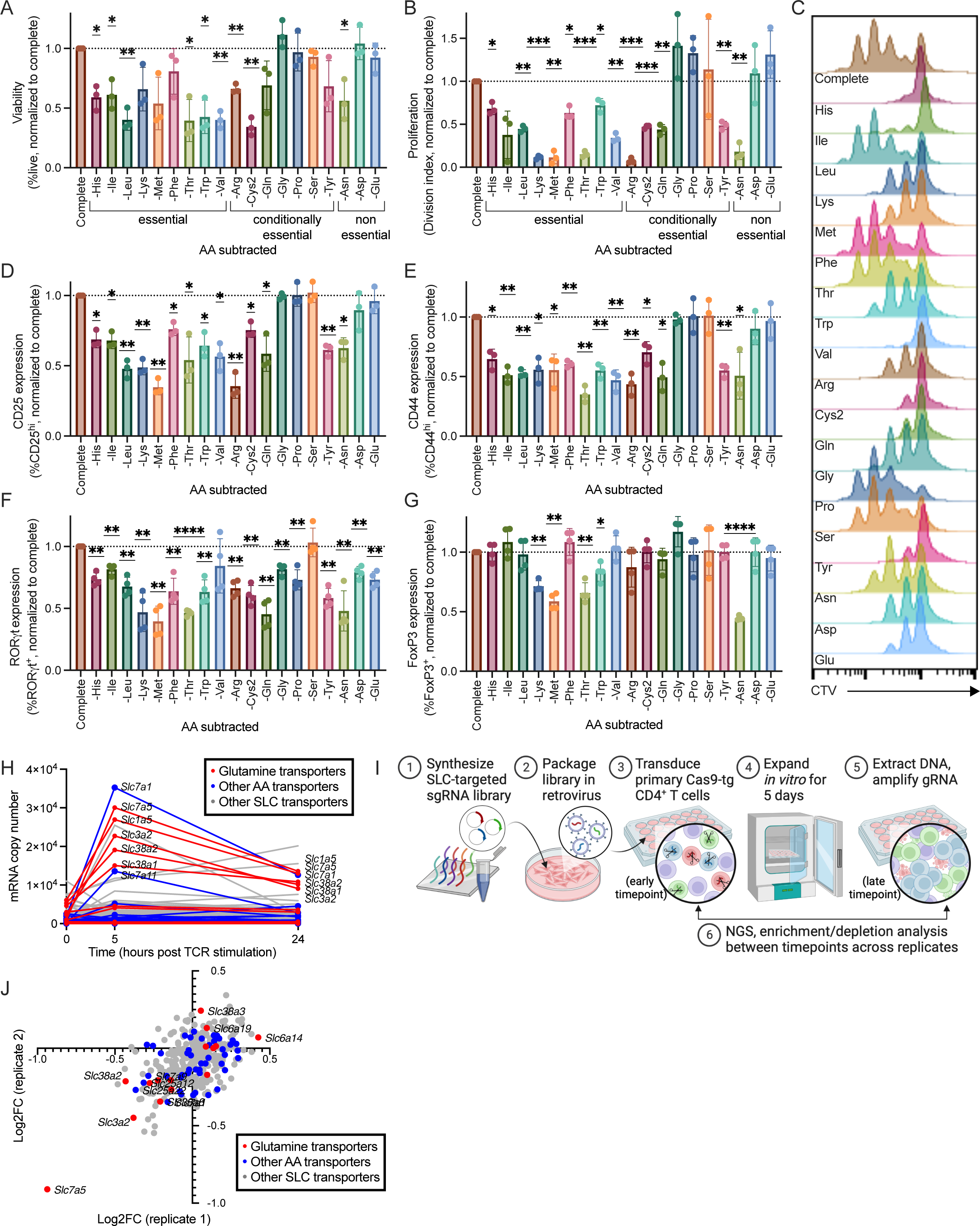
CD4^+^ T cells are dependent on uptake of select essential and non-essential AAs through dynamic regulation of SLC transporters. (A-E) Characterization of primary CD4^+^ T cells activated with anti-CD3 and anti-CD28 antibodies and cultured in vitro for 72 hours (Th0 cells) in either complete media or test media deficient in the indicated AA. (A) Cell viability normalized to complete media, (B-C) proliferation quantified as division index based on CTV dilution, and (D-E) expression of activation markers CD25 and CD44 as measured by flow cytometry (mean±SD, one-sample t-test, α=0.01, n=3 biological replicates representing at least three independent experiments compared to complete media). (F-G) Expression of lineage-characterizing transcription factors RORγt and FoxP3 in CD4^+^ T cells activated and cultured with Th17 or Treg cell polarizing cytokines in complete or test media for 72 hours (mean±SD, one-sample t-test, α=0.01, n=4 biological replicates across four independent experiments compared to complete media). (H) mRNA expression of SLC transporters in primary CD4*^+^* T cells at 0, 5, and 24 hours post activation with anti-CD3 and anti-CD28 antibodies, referenced from published RNAseq dataset ^24^. Red = transports glutamine, blue = transports other AAs, gray = all other SLCs detected (mean, n=3 biological replicates). (I) Experimental design for SLC gRNA library delivery to Cas9-transgenic primary CD4^+^ T cells and *in vitro* CRISPR screening. (J) Volcano plot showing change in gRNA abundancies from *in vitro* CRISPR screens performed using the SLC gRNA library in primary CD4^+^ T cells from Cas9-transgenic mice activated with anti-CD3/CD28 antibodies (replicate 1) or from OTII/Cas9 double-transgenic mice activated with OVA_323-339_ peptide (replicate 2). Red = transports glutamine, blue = transports other AAs, gray = all other SLCs detected (n=3 biological replicates with 2 technical replicates each, statistical analysis performed using MAGeCK ^46^). Data are available a https://figs.app.vumc.org/figs/. Ns denotes p>0.05, * p≤0.05, ** p≤0.01, *** p≤0.001, **** p≤0.0001. See also Figure S1.

### Teff cells are dependent on extracellular glutamine and TCR stimulation regulates multiple glutamine transporters

Within the SLC family, over 60 transporters are predicted to have AA-carrying capacity. To better understand mechanisms by which T cells acquire necessary AAs, we analyzed a previously collected RNAseq dataset ^24^ to survey changes in expression of SLCs in primary CD4^+^ T cells that are induced by TCR stimulation. At 5- and 24-hours post activation, Th0 cells markedly upregulated transcription of multiple AA-carrying SLCs, with highest mRNA copies detected at 5 hours for *Slc7a1, Slc7a5, Slc1a5, Slc3a2, Slc38a2, and Slc38a1,* and *Slc7a11* (**Figure 1H**). Notably, glutamine transporters were selectively overrepresented among the upregulated SLCs, including the sodium-dependent small neutral AA transporters *Slc1a5*, *Slc38a1*, and *Slc38a2* (**Figure 1H**). Further, import of essential Aas by the SLC7A5/SLC3A2 dimer and cystine for glutathione synthesis by SLC7A11/SLC3A2 dimer are coupled to glutamine and glutamate export, respectively, and therefore rely on maintenance of the intracellular glutamine pool ^4,25^. Moreover, expression of glutamine transporters was highly dynamic across T cell activation states and subsets as well as across multiple autoimmune diseases affecting different tissues (**Figure S1C-E**). These data suggest that, while multiple SLCs can have overlapping substrate profiles, differential expression and regulation may provide selective and non-redundant functions.

To test T cell AA transport and metabolism requirements, several different CRISPR screens were performed to cross examine relative dependencies. First, a custom guide RNA (gRNA) library targeting 362 SLCs with four gRNAs per gene and 20 non-targeting controls (NTCs) was curated from the whole-genome Mouse CRISPR Knockout Pooled Libraries, Brie ^26^ and GeCKO v2 ^27^ (**Table S1, Figure 1I**). The pooled SLC gRNA library was transduced into activated Cas9-transgenic CD4^+^ T cells, as previously described ^28^. T cells were sampled prior to and following *in vitro* expansion with anti-CD3/CD28 stimulation (replicate 1) to determine changes in gRNA abundancies. The same library was also tested in OTII/Cas9 double-transgenic CD4^+^ T cells activated with OVA_323-339_ peptide presented by irradiated splenocytes to confirm reproducibility (replicate 2). Of the SLCs included in this library, cells carrying the SLC7A5-targeted gRNAs were the most depleted from the final population, consistent with previously published literature showing that SLC7A5 is required for clonal expansion in CD4^+^ T cells ^10,11^ (**Figure 1J**). To a lesser extent, gRNAs targeting genes encoding the neutral AA transporters *Slc1a5, Slc38a1, Slc38a2, Slc38a8 (Snat8), Slc38a10 (Snat10), and Slc7a8 (Lat2),* the heavy chain subunit *Slc3a2,* the cationic AA transporters *Slc7a1* and *Slc7a9*, the phosphate transporter *Slc17a7*, the mitochondrial aspartate/glutamate-antiporter *Slc25a12*, and the mitochondrial ADP/ATP-antiporter *Slc25a12* were also depleted. Thus, the genes identified by the RNAseq and CRISPR screens were highly overlapping, with a notable overrepresentation of SLCs that transport glutamine and its derivatives (**Figure 1J**).

We next sought to better characterize the effects of extracellular glutamine deficiency on CD4^+^ T cell subsets. Primary CD4^+^ T cells were activated and cultured in either glutamine-replete (+glutamine) or glutamine-deficient (-glutamine) media. Here, glutamine deficiency was associated with significantly reduced proliferation in Th1, Th17, and Treg cells (**Figure S1F**), and viability was reduced in Th1 and Th17 cells but not Treg cells (**Figure S1G**). Under these conditions, Th1 and Th17 cell differentiation as measured by T-bet and RORγt expression, respectively, was also impaired, while FoxP3 induction and maintenance were spared in Treg cells (**Figure S1H**). Additionally, cells cultured in glutamine-deficient media exhibited decreased effector functions with fewer IFNγ^+^ Th1 cells and IL-17^+^ Th17 cells, while the fraction of IL-2^+^ Th1 cells was increased (**Figure S1I**). Lastly, the activity of mTORC1, a central nutrient sensor, was significantly reduced as measured by the expression of its downstream target phosphorylated ribosomal protein S6 (phosphor-S6) in both Th1 and Th17 cells as expected (**Figure S1J**). These data are in line with previously published literature ^3–7^, and supported further investigation of relative dependencies of CD4^+^ T cell subsets on glutamine transport and metabolism.

### Glutamine transport and metabolism requirements vary depending on cell type, experimental setting, and tissue microenvironment

To interrogate relative CD4^+^ T cell dependencies on specific glutamine transport and metabolism pathways, we next synthesized a second gRNA library targeting 32 glutamine uptake and metabolizing genes including *Slc1a5*, *Slc38a1,* and *Slc38a2* (**Table S2**). This smaller gRNA library was used to perform an *in vitro* CRISPR screen in Th0 cells as before. When cells were sampled after a brief interval in culture, cells with *Slc1a5, Slc38a1, and Slc38a2* gRNAs were significantly depleted at a similar magnitude as in the initial SLC screens (**Figure 2A**). Notably, loss of any one of these transporters at this timepoint had a greater effect size than loss of any of the other genes encoding glutaminolysis-associated enzymes. Upon re-stimulation and further expansion with IL-2 supplementation for an additional 48 hours of culture *in vitro*, however, only transporter *Slc1a5* and *Gfpt1, Cad, and Ppat* gRNAs targeting hexosamine and nucleotide synthesis were depleted (**Figure 2B**).

**Figure 2:**
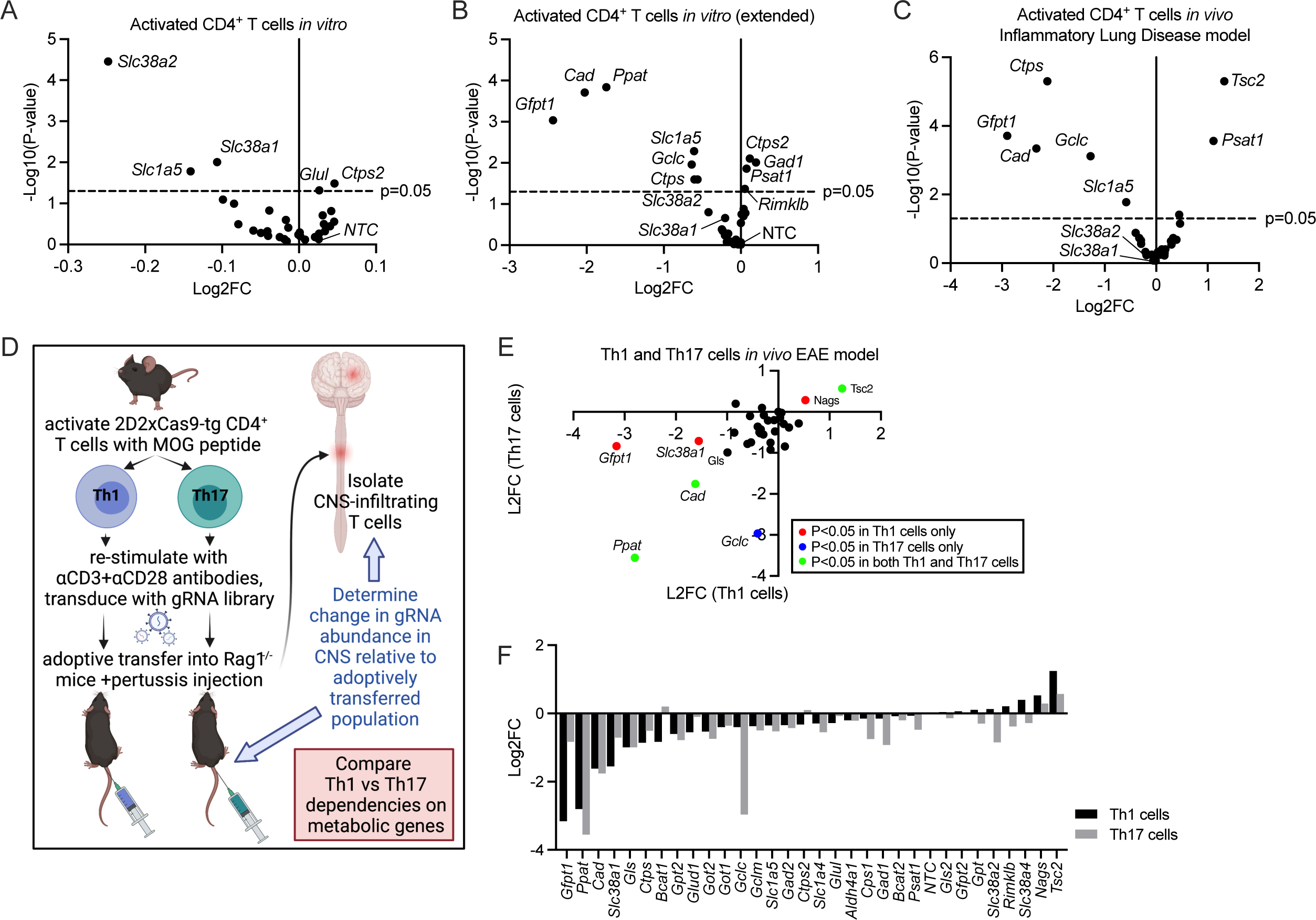
AA transporter requirements are subset-specific and dependent on culture conditions *in vitro* and tissue microenvironment *in vivo*. (A-B) Volcano plots showing change in gRNA abundancies from *in vitro* CRISPR screens performed using the glutamine gRNA library, with cells cultured for either (A) 7 days or (B) 9 days post gRNA transduction (n=3 biological replicates with 2 technical replicates each, statistical analysis performed using MAGeCK ^46^). (C) Volcano plot showing change in gRNA abundances from *in vivo* CRISPR screening performed in a model of inflammatory lung disease using the same glutamine gRNA library following previously published methods ^21^ (n=3 biological replicates with 2 technical replicates each, statistical analysis performed using MAGeCK). (D) Workflow for *in vivo* CRISPR screens performed in Th1- and Th17-cell driven EAE model. Disease induced by adoptively transferring 2D2/Cas9-double transgenic primary CD4^+^ T cells activated with MOG_35-55_ peptide transduced with the gRNA library into Rag1^-/-^ mice. (E) Results obtained from *in vivo* CRISPR screens performed in Th1- and Th17-driven EAE models as outlined in (D). Change in gRNA abundance for Th1 cells is shown on the x-axis and for Th17 cells on the y-axis. Green denotes statistical significance only in Th1 cells, blue denotes significance only in Th17 cells, red indicates significance in both subsets (statistical analysis performed by MAGeCK, representative plot from n=3 biological replicates with 2 technical replicates each across 2 independent experiments). Data are available a https://figs.app.vumc.org/figs/. (F) Same data as (E) represented as bar graphs to directly compare gene dependencies in Th1 and Th17 cells in the setting of EAE. Ns denotes p>0.05, * p≤0.05, ** p≤0.01, *** p≤0.001, **** p≤0.0001. See also Figure S2.

The smaller glutamine gRNA library was next used to perform *in vivo* CRISPR screens in several different disease models. There is growing evidence that the nutrient composition of the extracellular environment can significantly impact T cell metabolism and function. Previous studies have demonstrated how different synthetic medias compared to human plasma-like media can cause large shifts in the cellular metabolome *in vitro* ^29–31^, as well as in the setting of the tumor microenvironment *in vivo* ^1^. Here, it was hypothesized that nutrient composition may also vary across inflammatory disease processes involving different cell types and tissue microenvironments that would translate to distinct metabolic dependencies in the associated pathologic T cell populations.

The glutamine gRNA library was first used to perform an *in vivo* CRISPR screen in a model of inflammatory lung disease ^28^. OTII/Cas9 double-transgenic primary CD4^+^ T cells were activated with OVA_323-339_ peptide, transduced with the gRNA library, and adoptively transferred into *Rag1^-/-^* mice by tail vein injection on day 0. The recipient mice were administered intranasal ovalbumin on days 1, 3, 5, and 7 to promote recruitment of the transferred cells to the lungs and induce inflammation. 24 hours after the last treatment, T cells were recovered from the lungs and associated lymph nodes for gRNA sequencing. Relative to the baseline gRNA frequencies established from the adoptively transferred population, the lung population was significantly depleted of *Gfpt1*, *Cad*, *Ctps*, *Gclc*, and *Slc1a5* gRNAs while *Tsc2* and *Psat1* gRNAs were enriched (**Figure 2C**). Notably, *Slc38a1* and *Slc38a2* gRNA frequencies were not significantly changed in this setting, indicating that these transporters were dispensable for pathogenic T cells infiltrating the lung while loss of *Slc1a5* conferred only a modest disadvantage (**Figure 2C**).

The same approach was next applied to a model of Experimental Autoimmune Encephalomyelitis (EAE), representing a CNS demyelinating disease driven by autoreactive T cells akin to multiple sclerosis. While EAE classically involves the combined activity of pathogenic Th1 and Th17 cells, the model was modified according to previously published protocols ^32^ to test the two lineages independently in comparable *in vivo* tissue microenvironments. 2D2/Cas9 double-transgenic primary CD4^+^ T cells expressing myelin oligodendrocyte glycoprotein (MOG)-specific TCR were activated with MOG_35-55_ peptide. These cells were then polarized to either the Th1 or Th17 cell subsets, re-stimulated with anti-CD3 and anti-CD28 antibodies to induce pathogenicity, transduced with the glutamine gRNA library, and adoptively transferred into separate *Rag1^-/-^* mice. After several weeks, the recipient mice developed expected clinical manifestations of CNS demyelination characterized by ascending paralysis in the Th1 cell-driven model and ataxia in the Th17 cell-driven model. At the study endpoint, Th1 and Th17 cells were recovered from the brain and spinal cord of the recipient mice for gRNA sequencing. The relative change in gRNA abundancies were then compared between the Th1 and Th17 cell-driven models to identify differential requirements for glutamine transport and metabolism (**Figure 2D**). In this model, *Ppat and Cad* gRNAs were similarly depleted in Th1 and Th17 cells, indicating shared requirement for *de novo* nucleotide biosynthesis pathways (**Figures 2E-F, S2A-B**). On the other hand, *Tsc2* gRNAs that were included as positive controls were again enriched in both subsets as expected with loss of negative regulation of mTORC1 activity (**Figures 2E-F, S2A-B**).

Notably, gRNAs targeting *Gfpt1* and *Slc38a1* were only depleted in the Th1 cells, while those targeting *Gclc* were only depleted in Th17 cells (**Figures 2E-F, S2A-B**). These data suggest that Th1 and Th17 cells may be distinguished by their differential dependencies on SLC38A1-mediated AA import, hexosamine biosynthesis catalyzed by glutamine-fructose-6-phosphate transaminase 1 (GFPT1), and glutathione synthesis mediated by glutamate/cysteine ligase catalytic unit (GCLC). Incongruencies between the results of the *in vivo* CRISPR screens between the inflammatory lung disease model and the EAE model highlight how T cell metabolic requirements may vary between subsets as well as across different tissues.

### Direct genetic testing validates a role for SLC38A1 in Th1 cells

We next directly validated the effects of targeted SLC38A1 loss using CRISPR/Cas9. Primary Cas9-transgenic CD4^+^ T cells were activated, polarized to Th1, Th17, or Treg cell lineages, and transduced with one of three unique NTCs (WT) or one of two unique *Slc38a1* gRNAs (Δ*Slc38a1)* marked with a Thy1.1 reporter. The frequencies of the Thy1.1^+^ gRNA-transduced populations were monitored over time to determine relative fitness. This showed that SLC38A1 loss significantly hindered Th1 cell expansion, mildly reduced Treg cell proliferation, but had no measurable effect on Th17 cell proliferation (**Figures 3A, S3A)**. SLC38A1 protein expression was quantified for WT and *ΔSlc38a1* Th1 and Th17 cells by immunoblot, confirming reduced expression of SLC38A1 protein in the Δ*Slc38a1* condition compared to WT in both Th1 and Th17 cells, as well as overall lower expression of the transporter in Th17 cells compared to Th1 cells (**Figure 3B**) consistent with the proteome atlas data (**Figure S1D)**. Residual expression of SLC38A1 was noted in the *ΔSlc38a1* conditions (**Figure 3B**), which may be attributed to the remaining Thy1.1^-^ cells following Thy1.1^+^ enrichment by positive in addition to the small fraction of Thy1.1^+^ cells that failed to successfully disrupt *Slc38a1*.

**Figure 3:**
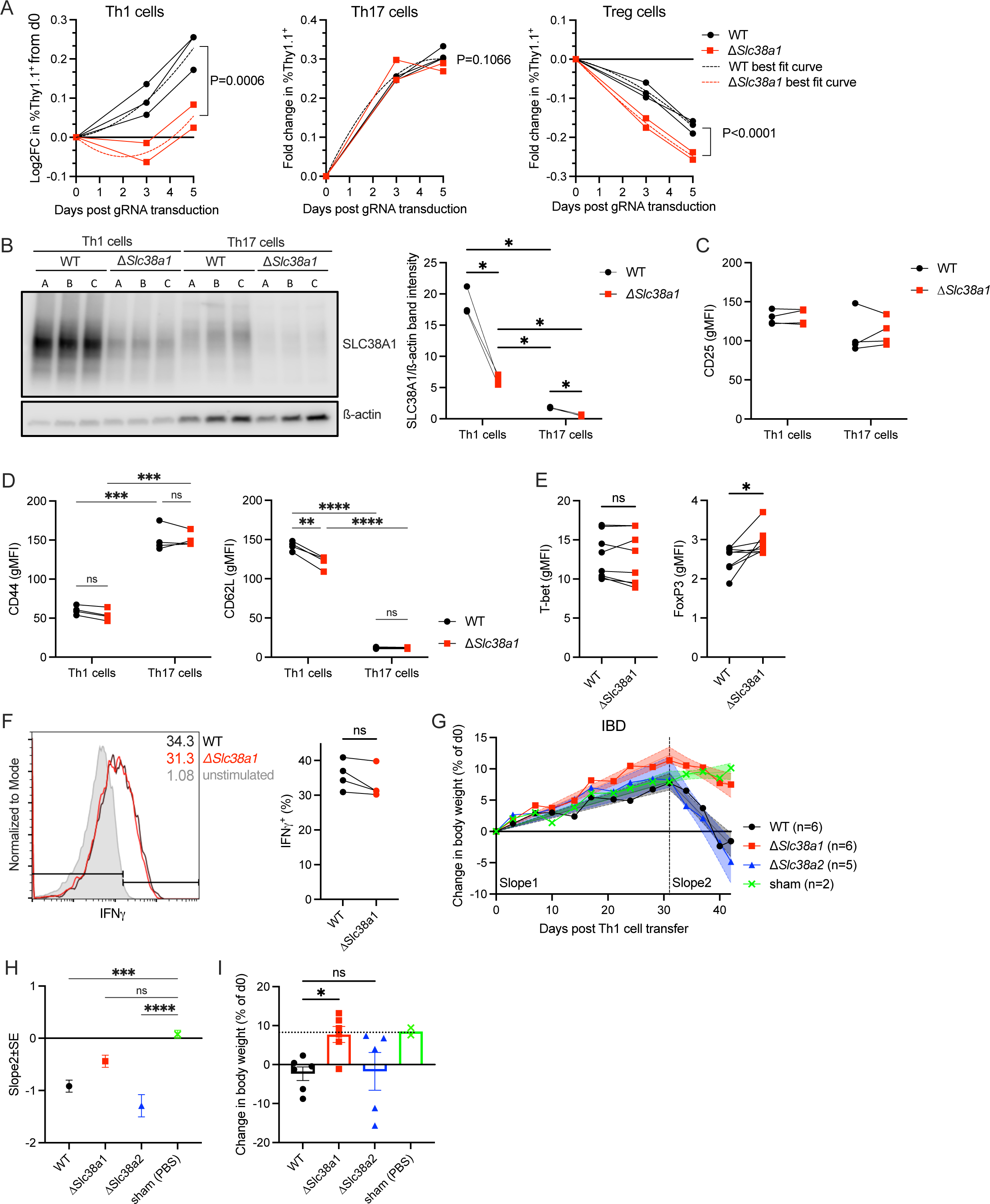
Genetic ablation of *Slc38a1* selectively impairs Th1 cell proliferative capacity, resulting in lesser colitis-associated weight loss in a model of Th1 cell-driven IBD. (A) Change in Thy1.1^+^ frequency over five days from day 2 post transduction with NTC (WT, black) or *Slc38a1*-targeted gRNA (*ΔSlc38a1*, red) in activated Cas9-transgenic Th1, Th17, and Treg cells (mean±SD, comparison of quadratic least squares fits, dotted lines show corresponding best fit curves, n=3 unique NTC gRNA sequences and n=2 unique *Slc38a1*-targeted gRNA sequences). (B) Immunoblot of SLC38A1 and ß-actin in WT and Δ*Slc38a1* Th1 and Th17 cells after isolation by Thy1.1-positive selection using magnetic cell-separation column (representative blot from n=3 independent experiments). Graph shows quantification of SLC38A1/ß-actin band intensity ratios from immunoblot (paired t-test, n=3 biological replicates). (C-D) Mean expression of activation markers (C) CD25, (D) CD44 and CD62L in WT and *ΔSlc38a1* Th1 and Th17 cells (repeated measures two-way ANOVA with Tukey’s multiple comparisons test, n=4 biological replicates across 2 independent experiments). (E-F) Expression of (E) transcription factors T-bet and FoxP3 and (F) effector cytokine IFN in WT and Δ*Slc38a1* Th1 cells (paired t-test, n=8 biological replicates for transcription factors and n=4 biological replicates across 2 independent experiments, flow cytometry histogram shows representative replicate). (G-I) Effect of SLC38A1 and SLC38A2 deficiency on IBD disease course as measured by (G) percent change in body weight from day of adoptive transfer trended over 42 days. IBD was induced in *Rag1*^-/-^ mice by transferring WT, Δ*Slc38a1*, or Δ*Slc38a2* Th1 cells generated using CRISPR/Cas9 or PBS control for sham by intraperitoneal injection. After period of relatively table weight gain in all groups (Slope 1), the (H) WT, Δ*Slc38a1*, and Δ*Slc38a2* groups lost weight while the sham group continued to linearly gain weight at from 31 to 42 days post T cell transfer (Slope 2). (I) Percent change in body weight from day of adoptive transfer and at the end of study (mean±SEM, one-way ANOVA with Dunnett’s multiple comparisons test, n=6 biological replicates for WT, n=6 for Δ*Slc38a1,* n=5 for Δ*Slc38a2*, and n=2 for sham). Ns denotes p>0.05, * p≤0.05, ** p≤0.01, *** p≤0.001, **** p≤0.0001. See also Figure S3.

CRISPR/Cas9-mediated SLC38A1 deletion did not significantly impact the expression of the activation markers CD25 or CD44 but did result in a small reduction in CD62L expression in Th1 but not Th17 cells relative to WT (**Figure 3C-D**). Further, no changes were detected in expression of lineage-characteristic transcription factors T-bet and RORγt in Th1 and Th17 cells, respectively (**Figures 3E, S3B**). *ΔSlc38a1* Th1 cells were again found to have a minor increase in aberrant FoxP3 expression relative to WT (**Figure 3E**). Lastly, Th1 effector cytokine production was unaffected as measured by fraction of IFNγ^+^ cells among Thy1.1^+^ cells (**Figure 3F**). These data show that specifically ablating SLC38A1 results in a proliferative defect in Th1 but not Th17 cells, with sparing lineage identity and effector functions.

Given the differential effects of SLC38A1 and SLC38A2 deficiency yet similar substrate preference, we directly compared their roles of these transporters in a model of colitis. Primary CD4^+^ T cells were isolated from Cas9-transgenic mice, activated, polarized to Th1 cells, and transduced with non-targeting (WT), *Slc38a1* (*ΔSlc38a1)*, or *Slc38a2 (ΔSlc38a2)* gRNAs marked by a Thy1.1 reporter. Thy1.1^+^ cells were enriched by positive selection and i.p. injected into recipient Thy1.2^+^ *Rag1^-/-^* mice. Sham controls were injected with PBS. Recipient mice were monitored for symptoms of disease progression over the next six weeks ^33^. All groups gained weight until 4 weeks post T cell transfer, after which WT and Δ*Slc38a2* groups began to lose weight rapidly while the sham group maintained weight (**Figure 3G-I**). The Δ*Slc38a1* group, in comparison, exhibited slower progression of weight loss (**Figure 3G-I**). SLC38A1 but not SLC38A2, therefore, contributes to pathogenic T cell activity in colitis.

Despite the delay in weight loss, SLC38A1 deficiency did not provide ultimate protection from disease. All three experimental groups appeared to have inflamed colons with marked edema on gross inspection and abundant CD3^+^ cell infiltration microscopically at the experimental endpoint (**Figure S3C**), consistent with *in vitro* studies showing spared differentiation and effector functions. At the end of study, there were no statistically significant differences across the experimental groups in the total number of CD4^+^ T cells recovered from the mesenteric lymph nodes (MLNs) (**Figure S3D)** or spleens (**Figure S3E)**. Moreover, the Thy1.1^+^ fractions within these MLN and splenic CD4^+^ T cell compartments contracted relative to the population transferred on day 0 in both the *ΔSlc38a1* and *ΔSlc38a2* groups when normalized to WT (**Figure S3F-G**). Among the MLN Thy1.1^+^ population, there were no statistically significant differences in T-bet expression (**Figure S3H**), IFNγ or IL-17 production (**Figure S3I**), or mTORC1 activity (**Figure S3J).** Only the FoxP3^+^ fraction was marginally increased (**Figure S3H)**. Together with findings from the *in vitro* studies, these data suggest that SLC38A1 supports Th1 cell proliferation but is dispensable to ultimately develop and maintain effector functions, such that loss of the transporter delays clinical progression of colitis without conferring complete protection.

### SLC38A1 deficiency in Th1 cells is selectively associated with diminished availability of intracellular glutamine with concurrent changes in the transcriptome and metabolome

To identify potential mechanisms underlying the proliferative defect in Δ*Slc38a1* Th1 cells, transcriptomic and metabolomic changes were surveyed by RNAseq and liquid chromatography-tandem mass spectrometry (LC-MS/MS)-based metabolomics. For these experiments, WT and *ΔSlc38a1* Th1 and Th17 cells were prepared by gRNA transduction followed by five days of expansion in culture to allow for sufficient protein turnover before enrichment by Thy1.1 positive selection. These cells were used for all further studies including RNAseq, metabolomics, NMR, and extracellular flux analysis. As anticipated, WT Th1 and Th17 cells had numerous gene expression and metabolite differences (**Figure S4A-B)**, with overrepresentation of phosphatidylinosityol signaling pathways (**Figure S4C**). Analysis of the RNAseq dataset verified that *Slc38a1* mRNA expression was indeed reduced in both the *ΔSlc38a1* Th1 and Th17 cells relative to their WT counterparts (**Figure 4A-C**). Most strikingly, while this was accompanied by altered expression of numerous other genes in *ΔSlc38a1* Th1 cells, no other significant changes were detected in *ΔSlc38a1* Th17 cells (**Figure 4A-B**). Among genes with altered expression in *ΔSlc38a1* Th1 cells relative to WT were multiple SLCs, including the glucose transporters *Slc2a1* and *Slc2a3,* and the SNAT *Slc38a3* (**Figure 4C**). Given downregulation of glucose and AA, these data suggest that the rate of anabolic metabolism was likely reduced in *ΔSlc38a1* Th1 cells relative to WT.

**Figure 4:**
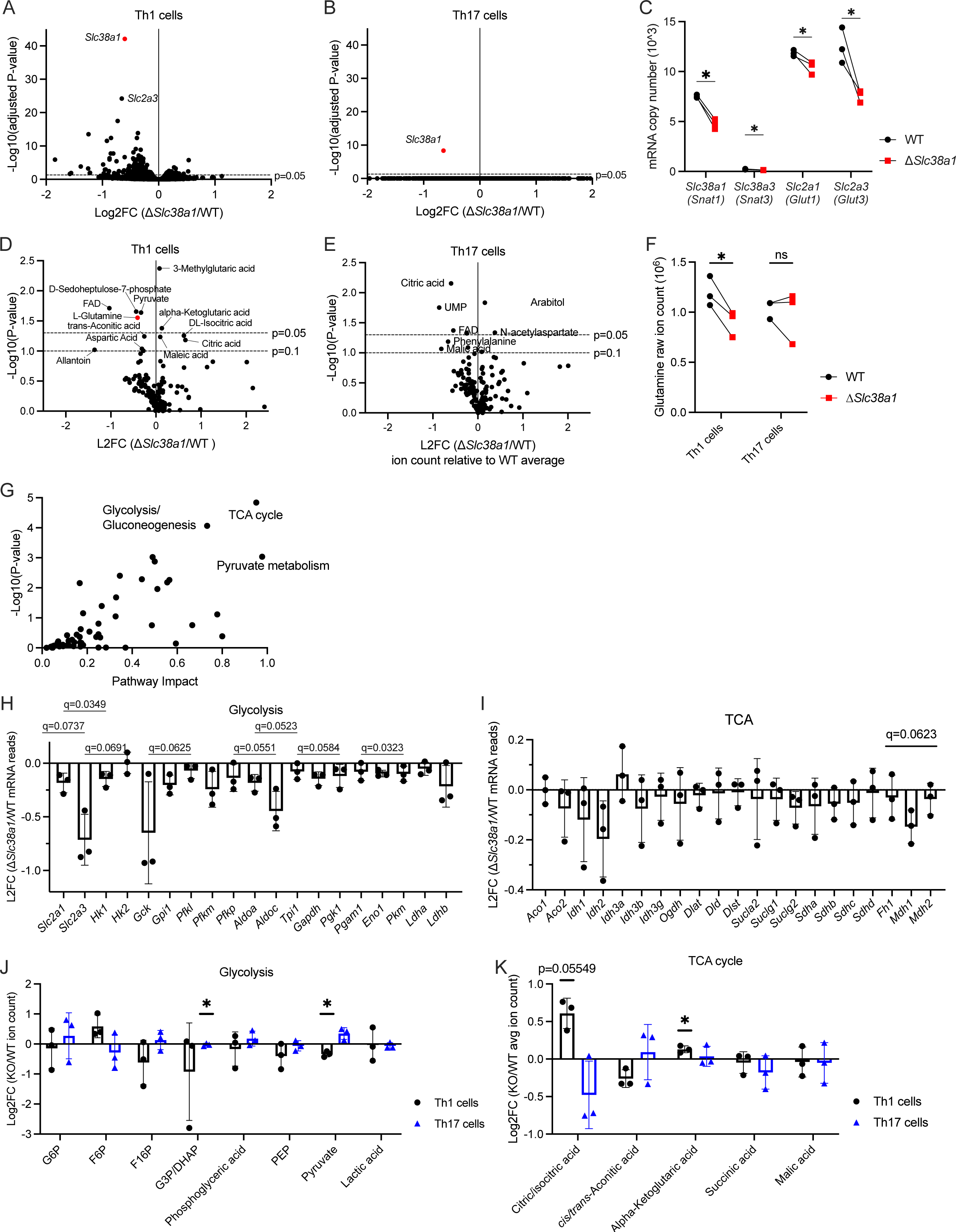
SLC38A1 loss is associated with altered metabolic activity mediated in part by reduced intracellular glutamine pool in Th1 cells but not Th17 cells. (A-B) Volcano plot of RNAseq data showing fold change in mRNA expression between WT and Δ*Slc38a1* (A) Th1 and (B) Th17 cells measured by RNAseq (data analyzed with DESeq2 ^47^, n=3 biological replicates). (C) mRNA copy numbers of SLCs with the most significant expression changes between WT and Δ*Slc38a1* Th1 cells detected by RNAseq from (A) (paired t-tests with multiple comparisons corrections by FDR where Q=5%, n=3 biological replicates). (D-E) Volcano plot of mass spectrometry data generated from the same samples as (A-B) showing change in metabolite ion counts in WT and Δ*Slc38a1* (D) Th1 and (E) Th17 cells (multiple paired t-tests, n=3 biological replicates) (F) Ion counts of glutamine in WT and Δ*Slc38a1* Th1 and Th17 cells as detected by mass spectrometry from (D-E) (repeated measures two-way ANOVA with Tukey’s multiple comparisons test, n=3 biological replicates). (G) Joint pathway analysis combining significantly changed genes from RNAseq and compounds from mass spectrometry performed on the same samples, identifying differential pathway dependencies between WT and Δ*Slc38a1* Th1 cells (data analyzed with MetaboAnalyst v5.0 ^48^, n=3 biological replicates). (H-I) Changes in mRNA read counts of (H) glycolysis pathway and (I) TCA cycle associated metabolites in Δ*Slc38a1* Th1 cells relative to WT, referenced from RNAseq dataset from (A-B) (mean±SD, one-sample t-tests with multiple comparisons corrections by FDR where Q=5%, n=3 biological replicates). (J-K) Changes in ion counts of (J) glycolysis pathway and (K) TCA cycle associated metabolites in Δ*Slc38a1* relative to WT Th1 and Th17 cells, referenced from mass spectrometry dataset in (D-E) (mean±SD, one-sample t-tests with multiple comparisons corrections by FDR where Q=5%, n=3 biological replicates and 2 technical replicates for negative and positive ionization modes). Ns denotes p>0.05, * p≤0.05, ** p≤0.01, *** p≤0.001, **** p≤0.0001. For multiple comparisons corrections using FDR, nd denotes no discovery and * q≥Q where Q=5% unless otherwise specified. See also Figure S4.

More metabolites were found to be altered in Th1 cells compared to in Th17 cells when assessed by metabolomics (**Figure 4D-F**). Specifically, *ΔSlc38a1* Th1 cells had increased amounts of citric acid and isocitric acid relative to WT Th1 cells, while having significantly reduced glutamine, glutathione (rGSH), flavin adenine dinucleotide (FAD), deoxyguanosine monophosphate (dGMP), adenosine monophosphate (AMP), cytosine, and cytidine (**Figure 4D**). In comparison, *ΔSlc38a1* Th17 cells had increased intracellular pools of the AAs aspartate, threonine, and asparagine relative to WT, while riboflavin availability was reduced (**Figure 4E**). Notably, glutamine concentrations were unchanged in SLC38A1-deficient Th17 cells (**Figure 4E-F**). These data show that while Th1 cells are dependent on SLC38A1 to maintain intracellular glutamine, SLC38A1-mediated glutamine import does not significantly contribute in Th17 cells. Given previously established dependence of Th17 cells on glutaminolysis ^8^, Th17 cells thus maintain the necessary intracellular glutamine pool by other means, whether that be import via a separate nutrient transporter, interconversion of other AAs, or scavenged from breakdown of macromolecules such from lysosomes or through autophagy.

To identify specific metabolic pathways impacted by loss of SLC38A1 and consequent reduction in intracellular glutamine availability in Th1 cells, genes with significantly changed mRNA transcript levels identified by RNAseq (Padj <0.05) and metabolites identified by mass spectrometry (P <0.1) were combined to perform joint pathway analysis. This revealed that glycolysis/gluconeogenesis, pyruvate metabolism, and the tricarboxylic acid (TCA) cycle were most significantly impacted by SLC38A1 loss in Th1 cells (**Figure 4G**). To better assess how these pathways were affected, change in mRNA expression of associated genes (**Figures 4H-I**) and concentrations of intermediate metabolites (**Figures 4J-K**) between *ΔSlc38a1* and WT Th1 cells were calculated. The RNAseq data showed that glycolysis was predominantly affected by downregulation in mRNA transcription of key glucose transporters and enzymes including SLC2A1 and SLC2A3 (**Figure 4C, H)**, while transcription of enzymes within the TCA cycle were largely unaffected (**Figure 4I**). The metabolomic data showed that the relative abundancies of glycolytic intermediates were largely unchanged in both Th1 and Th17 cells (**Figure 4J**). While intermediates most proximal to pyruvate entry to the TCA cycle, citrate and isocitrate, were increased in *ΔSlc38a1* Th1, the remainder were not significantly changed to suggest decreased glutamine anaplerotic flux (**Figure 4K**).

### SLC38A1 loss is associated with altered glutamine and alanine-dependent glycolytic and mitochondrial flux

To test for alterations in metabolic flux associated with impairment in SLC38A1-mediated transport of glutamine and alanine, a series of modified substrate oxidation tests designed to quantifying the relative contributions of SLC38A1-dependent AA import to fuel Th1 cell glycolysis and oxidative phosphorylation (**Figure 5A-B**). WT and *ΔSlc38a1* Th1 cells were plated in base assay media lacking both glutamine and alanine and equilibrated to establish baseline extracellular acidification rate (ECAR) and oxygen consumption rate (OCR) measures. Once the cells were equilibrated in the base media, vehicle, glutamine, or alanine was added at a final concentration of 2mM to determine whether substrate availability elicited any acute changes in metabolic activity. In WT Th1 cells, glutamine addition resulted in increased OCR (**Figure 5C**) and decreased ECAR (**Figure 5D**) relative to vehicle, consistent with previously published findings ^34^ and the established role of glutamine in anaplerosis to support mitochondrial metabolism. In comparison, *ΔSlc38a1* Th1 cells responded to glutamine addition with changes of approximately half the magnitude as that observed in WT cells (**Figure 5C-E**). The OCR and ECAR changes measured in the knockout condition may be attributed in part to the activity of the non-transduced Thy1.1^-^ cells, comprising approximately 50% of the total assayed population (**Figure S4D**) and the small minority of Thy1.1^+^ cells that retained native SLC38A1 activity.

**Figure 5:**
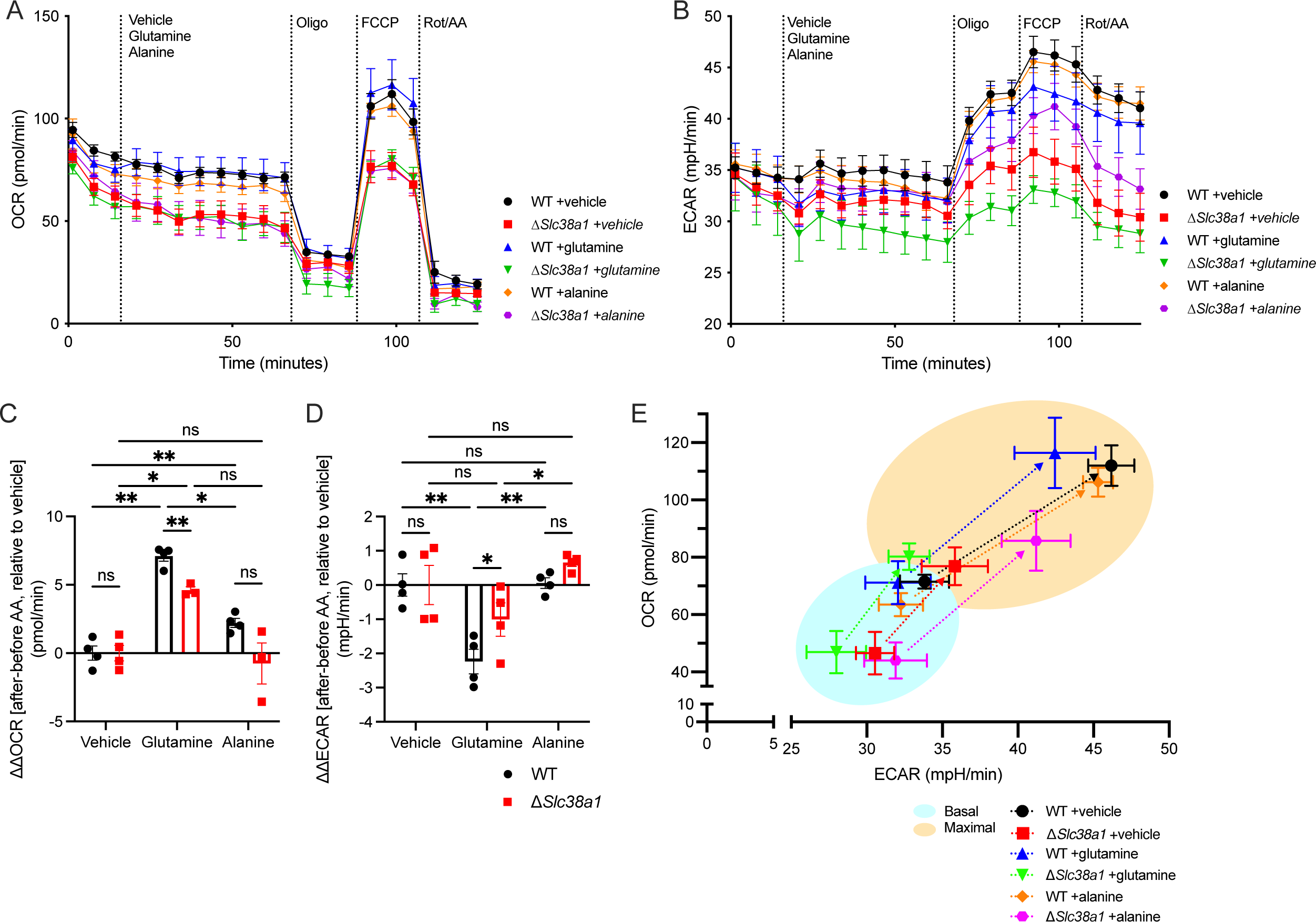
SLC38A1-mediated glutamine import is critical to support mitochondrial metabolism in Th1 cells. (A-B) Modified substrate oxidation test measuring changes in metabolic activity in WT and Δ*Slc38a1* Th1 cells with the addition of vehicle, glutamine, or alanine. (A) shows OCR reflecting rate of mitochondrial respiration and (B) shows ECAR reflecting concomitant changes in extracellular acidification as a function of glycolysis derived lactate and TCA cycle derived CO_2_ (mean±SEM, n=4 biological replicates with 2-4 technical replicates per condition). (C-D) Quantification of interval change in (C) OCR and (D) ECAR immediately before and after supplementing WT and Δ*Slc38a1* Th1 cells with vehicle, glutamine, or alanine (mean±SEM, ordinary two-way ANOVA with Tukey’s multiple comparisons test, n=4 biological replicates with 2-4 technical replicates per condition). (E) Energy phenotypes of WT and Δ*Slc38a1* Th1 cells after equilibration in vehicle, glutamine, or alanine-supplemented media (basal) and under stress upon exposure to electron transport chain modulators (maximal) as represented by the OCR:ECAR (mean±SEM, n=4 biological replicates with 2-4 technical replicates per condition). Ns denotes p>0.05, * p≤0.05, ** p≤0.01, *** p≤0.001, **** p≤0.0001.

### SLC38A1 provides AA uptake to support reactive oxygen species (ROS) management, protein synthesis, and mTORC1 signaling

In addition to serving directly as building blocks for protein synthesis, imported glutamine and alanine may participate in many different regulatory and metabolic pathways. The role of SLC38A1-mediated AA transport in total protein synthesis was measured using short-term puromycin exposure. Despite expressing multiple other potentially redundant transporters, *ΔSlc38a1* Th1 cells indeed incorporated less puromycin per cell relative to WT over the same assay period to indicate reduced protein synthesis (**Figure 6A**). Accompanying the deficit in mitochondrial respiration, *ΔSlc38a1* Th1 cells also had elevated mitochondrial superoxide compared to WT Th1 cells (**Figure 6B**). No difference was observed between WT and Δ*Slc38a1* Th17 cells as expected (**Figure 6B**). Moreover, SLC38A1 loss was also associated with reduced phosphor-S6 expression in Th1 cells indicating lower mTORC1 activity (**Figure 6C**), consistent with the function of mTORC1 as one of the central nutrient sensors.

**Figure 6:**
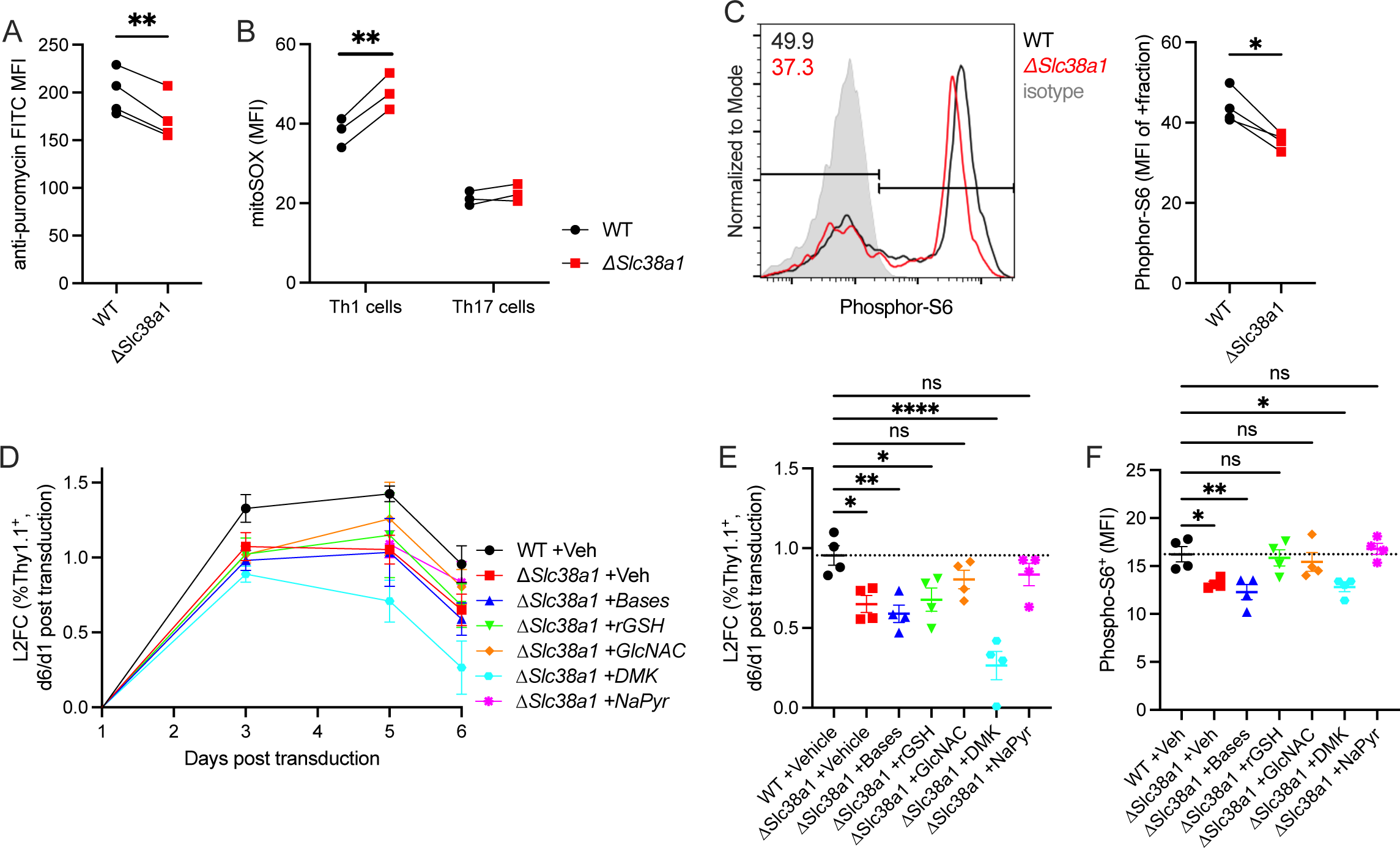
SLC38A1 function supports protein synthesis, and also modulates mitochondrial redox balance and hexosamine biosynthesis for glycosylation. (A) Rate of protein synthesis as measured by puromycin incorporation over one hour incubation period in WT and Δ*Slc38a1* Th1 and Th17 cells (paired t-test, n=3 biological replicates). (B) Mitochondrial superoxide levels measured by mitoSOX staining in WT and Δ*Slc38a1* Th1 and Th17 cells (ordinary two-way ANOVA with Tukey’s multiple comparisons test, n=3 biological replicates). (C) mTORC1 activity as measured by phosphor-S6 expression in WT and Δ*Slc38a1* Th1 and Th17 cells (paired t-test, n=4 biological replicates representative of 2 independent experiments). (D-F) Rescue experiments by exogenous supplementation of potential downstream metabolites including nucleotide bases, reduced glutathione (rGSH), GlcNAc, dimethyl-ketoglutarate (DMK), and sodium pyruvate (NaPyr) in ΔSlc38a1 Th1 cells compared to WT on (A) proliferative capacity as measured by relative change in Thy1.1^+^ fraction over time and on (B) day 6 post gRNA transduction. (C) shows phosphor-S6 expression on day 6 (mean±SD, one-way ANOVA with Dunnett’s multiple comparisons test, n=4 biological replicates across two independent experiments). Ns denotes p>0.05, * p≤0.05, ** p≤0.01, *** p≤0.001, **** p≤0.0001.

Next, a series of rescue experiments were performed to test if the proliferation defect in *ΔSlc38a1* Th1 cells could be restored by provision metabolites representing other potential fates of glutamine and alanine metabolism. Δ*Slc38a1* Th1 cell cultures were supplemented with vehicle, a solution of nucleotide bases, rGSH, GlcNAc, dimethyl-ketoglutarate (DMK) as the membrane permeable substitute for alpha-ketoglutarate, or sodium pyruvate (NaPyr) (**Figure 6D**). Provision of exogenous GlcNAc or NaPyr partially restored *ΔSlc38a1* Th1 cell culture proliferation rate to that of WT as measured by expansion of Thy1.1^+^ fraction (**Figure 6E**). Endogenous GlcNAc is produced through the hexosamine biosynthesis pathway using glycolytic intermediates and glutamine as substrate. In contrast, supplementation with bases or rGSH had no measurable effects, while DMK addition resulted in further contraction of the Thy1.1^+^ population (**Figure 6E**). The same approach was employed to test effect on mTORC1 activity by measuring phosphor-S6. Mirroring results of the previous experiment, GlcNAc and NaPyr supplementation again partially compensated for loss of SLC38A1 in supporting mTORC1 activity (**Figure 6F**). Of note, while provision of rGSH was not sufficient to rescue the proliferation defect associated with SLC38A1 loss, it did partially restore mTORC1 activity to that of WT cells (**Figure 6F**).

### Pharmacologic SLC38 inhibition selectively impairs Th1 cell expansion and function in the EAE model but has no measurable effect in the allergic-airway disease model

To functionally establish the effects of SLC38 inhibition, primary CD4^+^ T cells were activated with anti-CD3 and anti-CD28 antibodies and polarized to Th1, Th17, or Treg cells in the presence of 0, 1, or 5mM alpha-methylaminoisobutyric acid (MeAIB), a nonselective SLC38A1 and SLC38A2 competitive inhibitor. These concentrations were selected for MeAIB treatment to capture cell count and activation as measured by CD44 without significantly impairing cell viability (**Figures S5A-C).** MeAIB treatment resulted in decreased proliferation of all subsets in a dose-dependent manner (**Figure S5D)** and Ki-67 expression (**Figure S5E)**. However, viability was only reduced in Th1 cells (**Figure S5F**). Mirroring the proliferation defect, MeAIB treatment reduced activation as measured by CD25 expression (**Figure S5G**). Expression of lineage-characterizing transcription factors T-bet and RORγt were generally reduced with MeAIB treatment, though the expression of FoxP3 was mildly increased in Th17 cells and unchanged in Treg cells (**Figure S5H**). Further, MeAIB-treated Th1 and Th17 cells had reduced expression of effector cytokines IFNγ and IL-17, respectively (**Figure S5I**). CD8^+^ T cells also showed a dose-dependent reduction in total cell numbers with MeAIB treatment (**Figure S5J**), but with no accompanying change in cell viability (**Figure S5K).**

Metabolic extracellular flux assays were performed on primary CD4^+^ T cells activated and cultured in the presence of Th1 or Treg polarizing cytokines and either vehicle or 5mM MeAIB. MeAIB treatment had no effect on basal or maximal ECAR in Th1 cells, while both the basal and maximal ECAR were significantly reduced in MeAIB-treated Treg cells relative to vehicle (**Figure S6A-C)**. Th1 cells treated with MeAIB had lower rates of oxidative phosphorylation (OXPHOS) compared to vehicle though this difference was extinguished under stressed conditions. Both basal and maximal OCR in Treg cell were found to be unchanged with SLC38 inhibition (**Figure S6D-F)**, however, mitochondrial ROS increased selectively in Th1 cells as measured by MitoSox (**Figure S6G**).

Next, we sought to evaluate the therapeutic potential efficacy of pharmacologically inhibiting SLC38A1/2 in inflammatory disease. To test a Th17-based disease model, neutrophilic allergic-airway disease was induced by intranasal administration of house-dustmite (HDM) and lipopolysaccharide (LPS) solution on days 7 and 14 to promote sensitization with Th17 cell recruitment ^35^. Healthy control mice were given intranasal phosphate-buffered saline (PBS). The mice were given a final HDM/LPS challenge on day 21 and sacrificed the following day for bronchioalveolar lavage fluid (BALF) and lung tissue collection. The total number of live cells recovered from the long homogenate was unchanged between conditions (**Figure S7A**) and MeAIB treatment had no effect on CD4^+^ and CD8^+^ cell frequencies (**Figure S7B)** or the prevalence of CD4^+^IL-17^+^ cells in the lung tissue (**Figure S7C**). Moreover, there were no statistically significant differences in the immune cell composition of the BALF used to characterize disease burden, including lymphocytes, neutrophils, eosinophils, and macrophages (**Figure S7D).** A Th1-dependent model was next tested using EAE. Mice pre-treated with i.p. injection of vehicle or MeAIB daily for seven days prior to disease induction and continued daily until the end of study. EAE was induced by immunization with MOG/CFA emulsion on day 0 and augmentation with pertussis on day 0 and 1. Mice were scored daily according to clinical symptomology to monitor disease progression until humane endpoints were reached at approximately 5 weeks post induction (**Figure 7A**). While there was no difference in the maximal disease severity (**Figure 7B**), disease onset was delayed in mice treated with MeAIB compared to vehicle (**Figure 7C**). At the end of the study, spinal cords and brains were collected and dissociated for analysis by flow cytometry. MeAIB treatment group had significantly lower frequencies and counts of pathogenic CD4^+^ and CD8^+^ T cell infiltrating the spinal cords and brains (**Figure 7D-E, Figure S7E-F**). CD4^+^ T cells in the spinal cords had similar expression of lineage-characterizing transcription factors and cytokines except for a mild decrease in RORγt expression with MeAIB treatment compared to vehicle (**Figure S7G-H**). Together, these data show T cell metabolic requirements are subset- and tissue microenvironment-dependent and SNAT1 plays a selective role in Th1 cell proliferation or recruitment to diseased sites but not differentiation *in vivo*.

**Figure 7:**
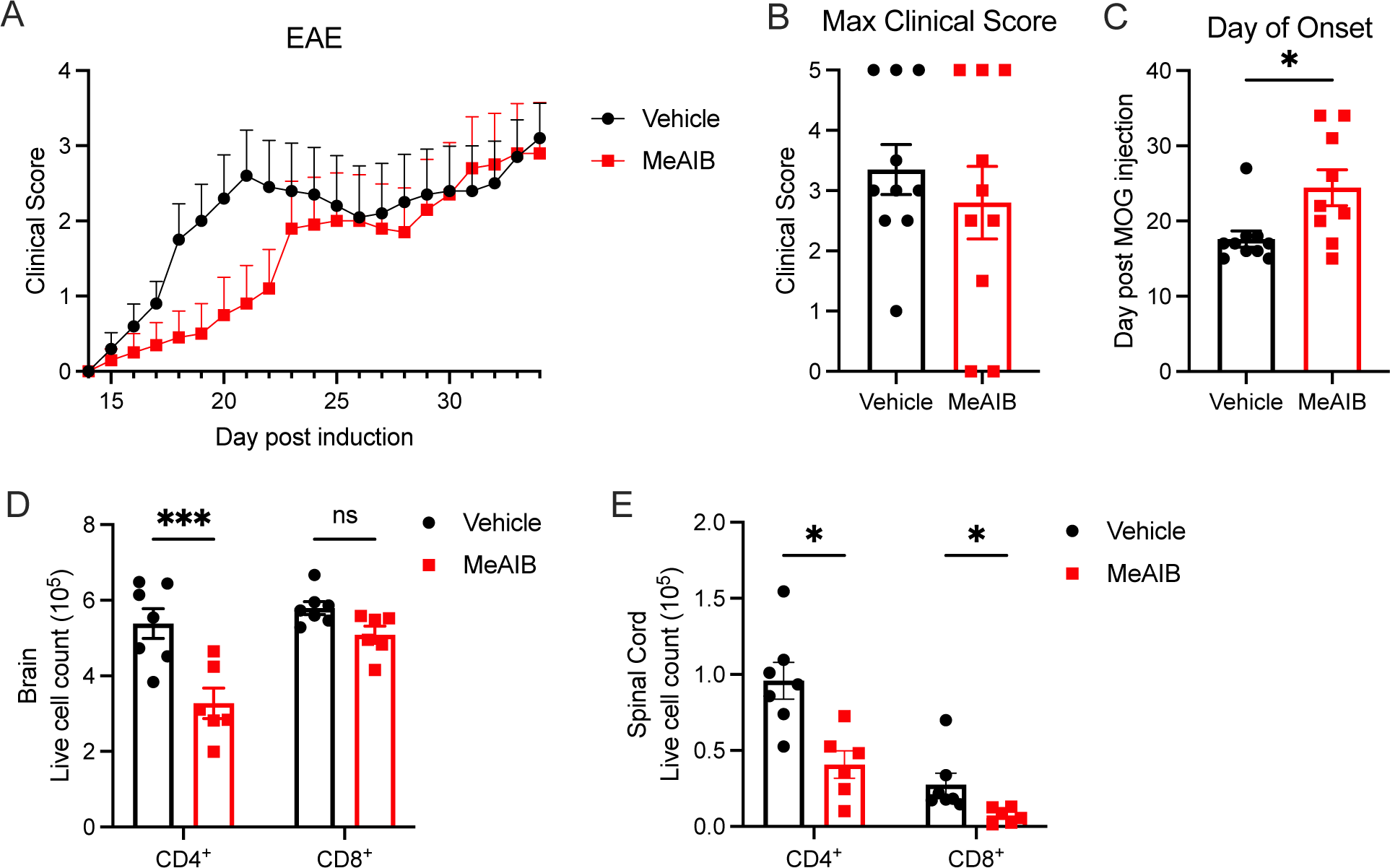
SLC38A1/2 inhibition *in vivo* delays disease onset of EAE. (A) Average clinical score overtime in mice immunized with MOG/CFA and PTX to induce EAE and treated daily with either i.p. vehicle or MeAIB from 7 days before immunization until end of study (mean±SEM, n=10 biological replicates for each condition). (B) Maximum clinical score attained in EAE mice from (A) (mean±SEM, unpaired one-tailed Mann-Whitney test, n=10 biological replicates for each condition). (C) Day of onset of clinical symptoms in EAE mice from (A) (mean±SEM, unpaired one-tailed Mann-Whitney test, n=10 biological replicates for each condition). (D-E) CD4^+^ and CD8^+^ T cell count and frequency in the (D) spinal cords and (E) brains collected from EAE mice at the end of study (mean±SEM, repeated measures two-way ANOVA with Šídák’s multiple comparisons test, n=7 biological replicates or vehicle and 8 for MeAIB). Ns denotes p>0.05, * p≤0.05, ** p≤0.01, *** p≤0.001, **** p≤0.0001. See also Figure S5-7.

## DISCUSSION

AA uptake and metabolism are critical to fuel and regulate T cell activities in health and disease. In this study, we investigated AA and nutrient transporter requirements in primary CD4^+^ T cell subsets to identify specific dependencies that may be exploited as an approach to immunotherapy. We designed and performed targeted *in vitro* and *in vivo* CRISPR screens using multiple models of autoimmune and inflammatory diseases to demonstrate cell type- and tissue microenvironment-dependent requirements for specific AA transporters. These studies revealed that pathogenic Th1 cells but not Th17 cells require the small neutral AA transporter SLC38A1 *in vivo* in the setting of EAE-associated CNS infiltration and inflammation. Notably, Th1 cells required SLC38A1 despite concomitant expression of multiple other transporters with overlapping substrate profiles. In contrast, SLC38A1 was dispensable for sustained expansion and persistence of Th0 cells both *in vitro* as well as *in vivo* in a model of inflammatory lung disease. Mechanistically, SLC38A1 loss in Th1 cells was associated with decreased intracellular glutamine, dampened mTORC1 signaling, alter glycolytic and mitochondrial metabolism, impaired ROS management and hexosamine biosynthesis, and slowed protein synthesis. Together, these results demonstrate that, while Teff cells generally require uptake of extracellular glutamine and alanine to meet biosynthetic and bioenergetic demands, specific transporter dependencies may vary between T cell subsets as well as across tissue microenvironments.

The family of SLC38A1/2 transporters have been described to preferentially carry glutamine ^25,36,37^ and or alanine ^15,38^ depending on experimental conditions. While both alanine and glutamine may be *de novo* synthesized intracellularly, additional supply may be needed from the extracellular environment when metabolic demands exceed synthetic capacity. Functionally, extracellular glutamine is required for Teff cell activation, expansion, fate, and pro-inflammatory functions ^3^, whereas glutamine deprivation promotes generation of Treg cells with suppressive capacity ^5^. This involves metabolic reprogramming mediated by TCR-induced upregulation of mTORC1 activity that is accompanied by rapid upregulation of multiple glutamine transporters including SLC7A5, SLC1A5, SLC38A1, and SLC38A2 ^3^. Upon import, glutamine is converted to glutamate by glutaminase (GLS), GLS deficiency attenuates Th17 cell differentiation while enhancing Th1 cell differentiation and effector functions through altered epigenetic and ROS regulation ^8^. Glutamine metabolism and CD4^+^ T cell subset activities are thus tightly intertwined. While SLC38A1/2 transporters have long been recognized as relevant glutamine transporters in CD4^+^ T cells, how these transporters fit into this intricate network of metabolic regulation in relation to T cell activity has not been directly investigated thus far.

As secondary active transporters, SLC38A1/2 transporters can drive steep concentration gradients and are responsive to microenvironmental factors including pH, hypoxia, tonicity, and nutritional deficiencies ^39,40^. One feature that distinguishes SLC38A1 and SLC38A2 from SLC1A5 is that the former are Na^+^-dependent symporters while the latter is a Na+-dependent antiporter ^41^. It has been proposed that these transporters contribute to cellular AA homeostasis in distinct ways based on their transport kinetics, with SLC1A5 acting in conjunction with SLC7A5/SLC3A2 to maintain a balanced mixture of AAs within the intracellular pool and SLC38A1 and SLC38A2 functioning to specifically accumulate their preferred substrates glutamine and alanine ^41^. In the present study, ΔSlc38a1 Th1 cells were found to have impaired proliferation and survival with associated depression in mTORC1 activity despite intact SLC1A5 expression. SLC38A1 status also had no measurable impact on Th1 cell lineage stability or effector functions. Notably, this phenotypic pattern is opposite of that observed for SLC1A5 ^4^. The same study also showed that SLC38A1 is similarly unable to compensate for SLC1A5 deficiency to support Th1 and Th17 cell differentiation and effector functions ^4^. Further, SLC1A5 was found to be required for mTORC1 signaling for naïve T cells but not activated Teff cells. These data together support the hypothesis that these transporters are non-redundant even though they have shared substrates, instead serving complementary functions depending on cellular demands and tissue location.

In the absence of SLC38A1, provision of extracellular GlcNAc partially rescued both proliferation and mTORC1 activity to suggest that glutamine uptake is crucial to support hexosamine biosynthesis. This pathway, like mTORC1, functions as a global nutrient sensor and integrator to calibrate cellular activities to reflect cellular nutrient supplies and demands. In the *de novo* synthesis pathway, glucose, glutamine, uridine triphosphate (UTP), and acetyl-CoA are combined to produce UDP-GlcNAc, whereas the salvage pathway may instead use imported or recycled glucosamine or GlcNAc. UDP-GlcNAc in turn serves as the substrate for N- and O-glycosylation reactions that can modify and regulate the activity of a wide array of proteins, lipids, and nucleic acids to induce global changes in cellular activities. This includes the expression and activity of essential T cell regulators including TCR, CD4, CD25, MYC ^42^, NFAT and NFkB ^43^, and PD-1 ^44^. Previous studies have shown that adequate expansion of the intracellular UDP-GlcNAc pool upon TCR stimulation is dependent on the uptake of extracellular glucose and glutamine ^45^, which is consistent with our findings that SLC38A1 deficiency, associated with decreased intracellular glutamine and downregulation of key glucose transporters, can be partially compensated by GlcNAc supplementation. Further investigation is warranted to better understand how immune cell dependencies on *de novo* hexosamine synthesis through targeting nutrient transport and salvage through dietary intake of glucosamine may be exploited in the context of anti-inflammatory and anti-tumor immunotherapies.

The findings from this study also underscore the utility of performing small-scale CRISPR screens *in vivo* using the most relevant disease model to identify physiologically relevant drug target candidates. This is highlighted by the distinct transporter requirements identified in this study across *in vitro* and *in vivo* platforms as well as between disease models involving CNS, pulmonary, and enteric inflammation. In sum, these data suggest that, while having overlapping substrate profiles, nutrient transporters may serve distinct roles in regulating T cell activities. Therefore, pathogenic T cell populations may be distinguished and selectively targeted based on their cell-type and tissue site-dependent requirements for specific nutrient transporter as an approach to immunotherapy.

### Limitations of the Study

In this study, the role of SLC38A1 was investigated by a combination of genetic and pharmacological approaches using CRISPR/Cas9-mediated gene editing and MeAIB treatment, respectively. The method employed for genetic ablation of SLC38A1 involved gRNA delivery into activated Cas9-transgenic primary T cells by retroviral transduction, which excluded the possibility of testing transporter requirements in naïve T cells and for activation events and requirements immediately following TCR-stimulation. In contrast, the pharmacologic approach provided the advantage of initiating treatment prior to T cell activation. However, this was limited by the lack of specificity in that MeAIB is a substrate for all system A transporters that include SLC38A1, SLC38A2, and SLC38A4. Thus, future studies are needed to develop models for inducible and conditional ablation of SLC38A1 to facilitate further characterization. Further, while the scope of the present study was limited to characterization of Th1 and Th17 cell subsets, the role of SLC38A1 in Th2 and Treg cell subsets as well as in CD8^+^ T cell populations in association to SLC1A5 and SLC38A2 is warranted given related findings in previous studies and the distinct transporter expression patterns observed among the cell types in health and in disease.

## AUTHOR CONTRIBUTIONS

A.S. and J.C.R. designed research; A.S., K.L.B., D.R.H., M.M.W., C.C., J.Y.C., and H.S.H. performed research; A.S., C.A.L., D.C.N., and J.C.R. analyzed data; and A.S. and J.C.R. wrote the paper with contributions from other authors.

## INCLUSION AND DIVERSITY STATEMENT

We worked to ensure sex balance in the selection of non-human subjects. One or more of the authors of this paper self-identifies as an underrepresented ethnic minority in science. While citing references scientifically relevant for this work, we also actively worked to promote gender balance in our reference list. The author list of this paper includes contributors from the location where the research was conducted who participated in the data collection, design, analysis, and/or interpretation of the work.

## ACKNOWLEDGEMENTS

We thank members of the Rathmell lab for contributing to this project. We acknowledge the expert technical support of the VANGARD core facilities, supported in part by the Vanderbilt-Ingram Cancer Center (P30 CA068485) and Vanderbilt Vision Center (P30 EY08126). We acknowledge the Translational Pathology Shared Resource supported by NCI/NIH Cancer Center Support Grant 5P30 CA68485-19 and The Shared Instrumentation Grant S10 OD023475-01A1 for the Leica Bond RX. We thank J. Cools (VIB) for providing the pMx-U6-gRNA-GFP construct. Diagrams were created with Biorender.com. This work was supported by the William E. Paul Distinguished Innovator Award for the Lupus Research Alliance (J.C.R.), R01s DK105550 (J.C.R.), HL136664 (J.C.R., D.C.N.), CA217987 (J.C.R.), and AI153167 (J.C.R.), and T32 GM007347 (A.S.).

