## Supplemental Table 2 for "Tissue-Specific Dependence of Th1 Cells on the Amino Acid Transporter SLC38A1 in Inflammation"

**Supplemental Table 2: Glutamine Metabolism gRNA Library**

| Target Gene | gRNA sequence |
|---|---|
| *Aldh4a1* | GTTCAACGCAAAGTTCGCCG |
| | CGCTCGGCATTCGAGTACGG |
| | CAACTGGTACTGTATATCGG |
| | ACCTTTATGACAGGGCAACG |
| *Bcat1* | TAGAAAAATAAGGTCCCACG |
| | CCCCACATTTATCGGAACTG |
| | CACAGCGTAGTGCAAAACAG |
| | CAAGATCCGATTGTTCCGGC |
| *Bcat2* | ACGGAACGAGCCTCTACGTG |
| | CAGGAACTATGGACCCACTG |
| | GTGGAGTGGAATAACAAGGC |
| | GCACAGAATGACGTACAGGA |
| *Cad* | CGCAGGGGTACCCGACCGTG |
| | AGGATTAGAACCTTTCGTGG |
| | ATGGTGAGTGCCCACCACAA |
| | CTCAGAAACTCTGTTACGGG |
| *Cps1* | TGAGCCTCACAATTTCGTCG |
| | ATGCAGACCGAATCATCACA |
| | TACAGTATTCCATGGAAGTG |
| | GTTGGTGGCATCTCGTGTCG |
| *Ctps* | ATTGGCCATTAACCACAAGC |
| | ATACCAGTACGTCATTAACA |
| | GCCCACAAGAGCGATCGAGC |
| | TTAATACCCGTAGACGAAGA |
| *Ctps2* | GGATAGCATCAGTAATATGG |
| | TTTGGTGTATTTACCAACCA |
| | AAAAACCCACCTGTGGCACA |
| | CTTCACAGCCATCTCAATGG |
| *Gad1* | CTATTCCATAAAGAAAGCCG |
| | GACATTTGATCGCTCCACCA |
| | GCGGTTGCATTGACATAAAG |
| | AGATGAGAGAGATCGTTGGA |
| *Gad2* | TTCTGATTAAATGTGACGAG |
| | GTATAAGATCTGGATGCACG |
| | GGCATCGGAAACAAGCTGTG |
| | GACAGCTGATTAAAATATCG |
| *Gclc* | TGTGCCGGTCCTTGACTGCG |
| | CAATATGAGGAAACGCCGGA |
| | AGAAACATCCGGCATCGGAG |

| | |
|---|---|
| | TGTAGATGATAGAACACGGG |
| *Gclm* | TACTTACCCTGACTAAATCG |
| | GTGCCCGTCCACGCACAGCG |
| | TTAACTCCATCTTCAATCGG |
| | GCATTTACAGCCTTACTGGG |
| *Gfpt1* | TGTGGCACAAGTTACCACGC |
| | AAGCTGCGGTCTTTCCCGTG |
| | GGAGAGAGGAGCCTTAACTG |
| | TCTGTTGTGAACACAATGAG |
| *Gfpt2* | CTGCGATACTGTAAGGATCG |
| | TACTGCACTTACAGCCACAG |
| | CACATGTCGGATATAAGACG |
| | AATTAGTGATGATCCCGTTG |
| *Gls* | CGACGCGTTCGGCAACAGCG |
| | TGTACATCGCTATGTTGGGA |
| | GATTGCGAACATCTGATCCC |
| | ATATAACTCATCGATGTGTG |
| *Gls2* | CGTCCGGTACTACCTCGGTG |
| | GGGGATCGGAATTACGCCAT |
| | AAAAGCAGGTCACCAAGTCG |
| | TGAGTCAGGCAGTGTCATGG |
| *Glud1* | CAGTAGCGGAGATGCGCCCG |
| | CCGCGGCGCCAGCATCGTAG |
| | ACATACAAGTGCGCTGTGGT |
| | GGAAGGAGAGGCTCAACACA |
| *Glul* | GATTACGGGGACAAATGCGG |
| | TGGAAGGCCAACCAAATGGG |
| | CTTGCCCAGAGTTACCTGAG |
| | TATTTCTAGAGACCAACTTG |
| *Got1* | GATCCCCGCAAGGTTAACCT |
| | GTTGGTGATGATACGTAGAT |
| | AGACCTAGAGAAAGATGCGT |
| | CATTCGGCCCTATTGCTACT |
| *Got2* | TGGAGGTCCCATTTCAACAT |
| | TTTCTGCCCAAACCATCCTG |
| | CATCCTCCTCACCTTCACCA |
| | AGCTCACCTTCCGGACACTG |
| *Gpt* | GTACTATGCGTCATCAACCC |
| | TTGATGACGCATAGTACTCG |
| | CCCTACCACGATGGCATCGC |
| | GTCCGGACTGCTCAGAAGAT |
| | GCGGTGGAGTACGCTGTGCG |

| | |
|---|---|
| *Gpt2* | ACGCTAAGAAACGAGCGCGG |
| | GTTCTCTGCATTATCAACCC |
| | GGGGATGGGAATCATCACGC |
| *Nags* | GGATGAACTAAGGCACAACG |
| | CAGCTACGGTGGCATCGTCG |
| | GACAGCCAGAAGGTGCCGTG |
| | GCTAGCGGCTGTAATGACTG |
| *Ppat* | ACCTTGGAATCGGACATACG |
| | ATAAGACGCCCGATGCAGAG |
| | TGATCACTCTGGGACTCGTG |
| | AGGGGTGTATGCGAGTAACT |
| *Psat1* | TGGAAGGAGTGCTGACTACG |
| | TGCAAACGAGACTGTGCACG |
| | CAATACAGAGAATCTTGTGA |
| | CTTTGTAGTCAAGGACTGAT |
| *Rimklb* | ACTATCACTCTGCACCCATG |
| | ATGGTGCTGACGGTCGAGCA |
| | GTACGAGTCATTGTTGTGGG |
| | AAGAATACAAGGGGTCATAG |
| *Slc1a4* | GTCTGCAACCGATTACACAG |
| | TAGAGCCACTCCTAACACCA |
| | GATGCCACCCAGACGCCCGA |
| | ACCCACCAACACTCCCGACA |
| *Slc1a5* | AATCCCTATCGATTCCTGTG |
| | TACAACAGAGTCGTTGATGG |
| | GCGGGAGATCAATTCAACCA |
| | GTGGTGTGCAGCCTGATCGG |
| *Slc38a1* | ATACTTTGGTGTGCACGCGT |
| | TGCATGGTGTATGAGAAGCT |
| | TCACCATCACCACCAACACT |
| | AGATTGGCAGGACGGACGGG |
| *Slc38a2* | CCACCAAAGCAGCTTCCACG |
| | CTCAAGACTGCCAACGAAGG |
| | GCAGTGACAATGGAAGAATG |
| | GAGTTGAAGATGAAATAGCG |
| *Slc38a4* | CCACGGACACAAATAAGACG |
| | GAAGAAGCTAGCCGATTACG |
| | GATCTCGCTGCCTAATGACT |
| | GATGAAGAGGTAGCTTGACA |
| *Rheb* | AACAAACTGAATTGTCAATG |
| | CCATATCCAACAACTTGCCA |
| | TTCAGCTTGTAGACACAGCG |

| | |
|---|---|
| | TCATAGGATACCTATTATGT |
| *Tsc2* | TGAACCACATGGCTATGACG |
| | CACAGGGTGATAATGAACAG |
| | CAGCTCCAAAGACCCTTGAG |
| | CTGATCCTAGCACACATGTG |
| NTC | AAACTCCCGTGTCAACCGAT |
| | AAACCTAGCGTAGATTCGGC |
| | AAAAAGTCCGCGATTACGTC |
| | AAACTCATACGTAGCGAATC |
| | AAAACGTAATTATACCGAGC |
| | AAAACGGCTCGATCGGTGAT |
| | AAACCCCCGCGCGGAGCGTC |
| | AAACGAGGCTGTTCGTACAC |
| | AAAATTGCACCTTCCCGGCC |
| | AAAGACGTGCATTCAGCGAG |
