## Supplemental Table 1 for "Tissue-Specific Dependence of Th1 Cells on the Amino Acid Transporter SLC38A1 in Inflammation"

**Supplemental Table 1: SLC gRNA Library**

| Target Gene | gRNA sequence |
|---|---|
| *Slc10a1* | TACAGCAAAGGAATCTACGA |
| | TTAACCCTCGGTCCTACCTG |
| | AGGACGTAGGGTACATAGTG |
| | GAGGGGCATGATACCGTACT |
| *Slc10a2* | CTATTGGATAGATGGCGACA |
| | GTTGCTCTCAGGTACTACGC |
| | TAGGACATATAAAGAGACCA |
| | GCTCACCATCCTCTTAGCCA |
| *Slc10a3* | CTGTTCATCAGCCTACCATG |
| | GTGATCTCTAGCCAATACAC |
| | TGGTGTTGAGAGCAGAAGGG |
| | TGAGAGTGGTAGGAAACCAG |
| *Slc10a4* | AAGCATCGGCTTTAGCCCCG |
| | CATGTCGCCGTCTACCAGCA |
| | CTCACCCTGTCTCCCAACGC |
| | AGGCACGGCTGTAGATCCAG |
| *Slc10a5* | TAGACACAATCTTTAACACA |
| | ACCCCAAGAAGTATTGGCAA |
| | GGATCATCGATAAGACAAGC |
| | GTCAACCTGAAGACTTTCCC |
| *Slc10a6* | TGCTCTGATACGGAATGACG |
| | CTGTGACATGGCCGACCAAG |
| | CAGTAACTGCCACCACCAGG |
| | GAAGGTGAGAACATTAGAGA |
| *Slc10a7* | CCGTCGGTCGGAGTGAACGG |
| | GTTGAAGAATATCGTTGCGA |
| | GAAGAAGCCACCATTTGGTG |
| | AACTGCCTTGGTTAAAATCA |
| *Slc11a1* | GGGTGTGCTACCACATACTG |
| | GAGAAGTAGACAGAACCCGC |
| | CCTAGCATGATACCGTCCAG |
| | GAATGGGGATCTTCTCACTC |
| *Slc11a2* | ATGTCACCGTCAGTATCCCA |
| | AAACACAAAAGTGTCTGCGA |
| | TGAGAAAATCCCCATTCCTG |
| | CCTTGACTAAGGCAGAATGC |
| *Slc12a1* | CATCATCGGTTCCATCACCG |
| | ACAAACGGAGTGGTGCGAGG |
| | GTGGGTTATATCCAAGAGAG |

| | |
|---|---|
| | CCATGGCTATCACAAAAGTG |
| *Slc12a2* | CGGTTTCCGAGAACGCCGGG |
| | AAACGTCCCTATGACGAAGT |
| | GTTAAGATGTAACCACGAAG |
| | TCCGACAACATACATAGCAA |
| *Slc12a3* | AACCTGGTACCCGACTGGAG |
| | CGGTTACAACACCATAGACG |
| | TGTGGTCTTCCACCTCGTTG |
| | CACAGGCTAGCCCTTCGCAG |
| *Slc12a4* | AGCAAACGGTGAACCGACGT |
| | GGTGGCGCTTGACATGTCAT |
| | GTTACTCACGGAAACACGGG |
| | CCCAGAAGTCTATCCCAGTG |
| *Slc12a5* | CCAAAGGGGTGATTGTCGAG |
| | AGCCATGGCGAGACAGCGTG |
| | TGGATTACAGAGCCTCACAG |
| | GCAGGATCTCGATCGTGCCA |
| *Slc12a6* | AATCCCCAGGATGTTACGGA |
| | AACTATACAAATCTGACACA |
| | AATGGGGGGATGGTATCCGT |
| | CAAGATCGACACAATTACAC |
| *Slc12a7* | AATGAAAGATGTAGTCACGA |
| | AACAACGTTACTGAGATACA |
| | TGGTCATGGAAAGGCCAACG |
| | AGCAGATGAACATCAGCGCG |
| *Slc12a8* | CATCTGCCCACCAAGCACCG |
| | CAGTGAAGAATGACTCCCCG |
| | GAAAGGGCCAAATAAAACAC |
| | CCAGTTCCTCTATTGGCTGG |
| *Slc12a9* | TAGATGCCGATCAACACGAG |
| | GCCATGCCGGTGTGGCACGG |
| | GCTCTTCTCACCGCACGAGG |
| | AAACACGACTATGCTAAACA |
| *Slc13a1* | AACATGGGGAACATTAACCC |
| | TTGAATCTGTAACATAACCC |
| | AAGAATTAGAGCAACCGGGA |
| | GGTGGGCTGACAACAATCAC |
| *Slc13a2* | TACACCACAGCAGCGCCATG |
| | ACCCCCGACGAACAATATGT |
| | GGGTGTGGTACCCGTCAGGG |
| | GCTAGCACGGACTCACAGGG |
| | AGTTTCTTGCCAGTTCGGAA |

| | |
|---|---|
| *Slc13a3* | TCCAGAGCCAGCCCACCAGT |
| | GGCTAAAGCGGTGATCCAGG |
| | CCACCGTACCTTGGGCGGCA |
| *Slc13a4* | AAACAACAGACAGACGTCCA |
| | GACGGTGCTGTGGTTTACCC |
| | GGTGGTCAGGCCACCAATGG |
| | CTCAGACACCCAGTACACAG |
| *Slc13a5* | GATAGCCGCCAATACAGCTG |
| | GGGTCCCGTCCCGGTCAAGG |
| | AGTCCAGAACCTTCAAAAGT |
| | AAGAAGGTGTGTTTACCGTG |
| *Slc14a1* | AAACGTATTGTAGTGTCCTG |
| | TCATAGACATAGCAGATACA |
| | AGGAGAGCAGGATAGCACAT |
| | GTTGCTGACAAACACCACCT |
| *Slc14a2* | GTCCCAGCAATCGTCCACCA |
| | GAGTACCATCTTCGCCAAGT |
| | CAGCAAAGTCACCTACCCGG |
| | ACTCACAAAGAGCTCCACTG |
| *Slc15a1* | TTGTCCAATCGTGTAGACGA |
| | GAAGTAAGGCATATCCCAAG |
| | CCACCAAACGCAGACACACA |
| | CTGGGACGACAATCTCTCCA |
| *Slc15a2* | AGGAGGTATCAAACCCTGTG |
| | ACATTCCAAAGCGACAACAT |
| | TATCGGCTGATCTCCAAGTG |
| | CTGATGGACTCCACCAAGAG |
| *Slc15a3* | GGTGGTGAGCAACAAGCCAG |
| | CAGCAACACCAAGTAGAGGG |
| | CAGAACATCAGCTTCCTATG |
| | CAACAAAGAGATGTTAGGGT |
| *Slc15a4* | GCGTGGAGGGCCGTTCACAG |
| | GACACGACGCTGACCCGTTG |
| | TATCACCACCACCCATCACA |
| | CCAATTGAAAAATCTCCGAG |
| *Slc15a5* | CAAACGCCCTGGCGATCGTC |
| | ACTTACTCATGCACGCTGAC |
| | ATGGCGTTCATTAGCGCAAT |
| | GGTTCTCTGGCCGATAGATC |
| *Slc16a1* | ACTACTAAGAAAGACCAAAG |
| | CACCAGCGATCATTACTGGA |
| | GACTTGCAGCCAACACCAAG |

| | |
|---|---|
| | AGGCCCTATTGGTCTCATCA |
| *Slc16a10* | TGGCATCCAGAACGCCTACG |
| | GCCACGTCGGAGACATCCGA |
| | GAGGTGCTCTTCATGTGCAT |
| | TGAAGCTCTTTAACACGCTG |
| *Slc16a11* | TTCGTTCAAAGTGCTCCGCG |
| | CTACCGCCAAGACCCGACGG |
| | GCGGACCCAAATGAACGTAG |
| | GCCAGCCGAAAGTATCAAGG |
| *Slc16a12* | CAACAATATCGATTACCCCG |
| | GCCAGGAAAACTGTTCGATG |
| | GCTTACCTGTAAGAACTCCC |
| | ACCAGTTATCCTGTCAAGCG |
| *Slc16a13* | CAGATGCCACGCGCCCCACG |
| | GATCGCTTCCATAGGGATCG |
| | CTGACAGCAGCCCAATACTC |
| | CATTCCATACGTCCACCTGG |
| *Slc16a14* | GGCCCTAGGAGTCCTCAACG |
| | AAAGGCTACAAACATGCGGT |
| | AATAAAGAGAGACTGCACAT |
| | AAGCCCCACCCAGATATCGA |
| *Slc16a2* | GTAGGGGACGAAGTAACCAA |
| | CCTCTACTCCATGCTACTAG |
| | CCACACCATTGGCTAGACCT |
| | GCCTGCGCTACTTCACCTAT |
| *Slc16a3* | TATGGGTGTACCCGACACAA |
| | CTGAGTGTCTTCCGAGACCG |
| | AGTATCGATTGAGCATGATG |
| | AAAAGACGCTGACCGCCTTG |
| *Slc16a4* | AAGGAACGTCCCAAGAATGG |
| | CAATTGCCCGTTCTGGAATG |
| | CTGGCTACTAGGTGGAAAGT |
| | TTGCCATTGCTCATAAGTAA |
| *Slc16a5* | GCCGTACGCTATGCATCATG |
| | AGGCTCCACATACACAACAG |
| | GGCTTCTGCATATACGTCAC |
| | CTGTGATCACTCCTGCGGTG |
| *Slc16a6* | GCTGTCGACGATGGCCATTG |
| | ACAGAATGGTCACAGTTGGG |
| | CAATTATTATCCAAGGGCCG |
| | TGAGTCGATGGAGTCTATTG |
| | ATTACCTCCAATGAAGCCAA |

| | |
|---|---|
| *Slc16a7* | AGAGGTACTGGATTCGTGGA |
| | GCTCAGTACGCTAAACACAT |
| | TTCACCAACACACTACTGAT |
| *Slc16a8* | AGAGACCCCTCGCCCCACGG |
| | GACAGCCAAAGCGCGTCACG |
| | CCTGCTGGTGAACTACGCCA |
| | GAAAGCAACGAGGGTCCCGT |
| *Slc16a9* | CTAACGGGGATCCGTAAGAG |
| | GGACAGTCGGAGGTTTGCAA |
| | AAACAGAAAGTAGATATTGG |
| | TACCTGTTGAAATCAGGCCG |
| *Slc17a1* | ACAGATTCGTTAGATAAATG |
| | ATTGTGTGTCGAGTACTCCA |
| | TGGGCACCTCCCTTAGAACG |
| | ATGTCGGCGTGTATGTAACC |
| *Slc17a2* | GAATCAAGGACTTTAGTACC |
| | ATCCCCGCTAAATATCCACT |
| | TGTGGGAGGATTAATCTCAC |
| | ATCTGACCATCGCCTTTATG |
| *Slc17a3* | TATGGCATGATACTGATGCA |
| | TCTGCACTATGACCCATCAG |
| | AGCGCATGGAACATAAACTC |
| | ATGTACCTGTTGGTTCAGAG |
| *Slc17a5* | TCGTCACCCAGATTCCCGGT |
| | TAGAACGTCTAAGGAGTGTG |
| | GATCGTTATCTTACCCGCAT |
| | GCTGGCCGCAGACTTAGGCG |
| *Slc17a6* | CAGGAGGATATATCGCATCG |
| | AGAAATTTAAGACCCCATGG |
| | AAGCACGTGCAGTCGCATAG |
| | AACAACAGCACTATCCACCG |
| *Slc17a7* | TGGCGATGATGTAGCGACGA |
| | GGAGGAGCGCAAATACATTG |
| | GACACAGCCATAGTGAACGC |
| | TCCATCCTGAATACTGCACA |
| *Slc17a8* | CAGCAATGATGTACCGTTTG |
| | AATGATGGCATAGACAGGCA |
| | ACTCGCTGGGCATCTTACAA |
| | TCAGCCAGTTGTCCTCCGAT |
| *Slc17a9* | GATTCGAGAGAATGTCAGGA |
| | GGCAACAGTACAGACGGGCA |
| | TTGGTTACCGATCCCCAAGG |

| | |
|---|---|
| | TCTGGGCGTACTATGTGTAC |
| *Slc18a1* | GGACAATATGCTGCTCACTG |
| | GTAGTTATCCGTATAGACAC |
| | TCATAAAGCCAACAAACATG |
| | CTAAAAACAACTGCTTGCAA |
| *Slc18a2* | CGAGCCATACGTACCTACGA |
| | CCATCTGCTTTGCAAACATG |
| | GTACATACCTAAGACCCCCA |
| | ATGCAGAATCCAGCAAACAT |
| *Slc18a3* | AGATAGACGCCTAACACGTG |
| | GGTCGGCTCGGTCAATCCTG |
| | GCTTGCCCGCGAACTCGTAG |
| | CTATATCGCTCACATGCGCG |
| *Slc18b1* | AGTCTGTCAACTGGATACGT |
| | TAACCCAAGAGTCCGATCCA |
| | GTGAGAATAGTAGACGCCAT |
| | GTATCAATCCTTTGGCTACG |
| *Slc19a1* | TCTTTCTAAAGCGCCCTAAG |
| | CCTGGAACGTAAATTCACCA |
| | CATGGCCAGCTACTCACGGG |
| | CCGGTGCATACTCAGCAGTG |
| *Slc19a2* | GGAGTAGTAGGCGATTTCGG |
| | GACTCAGCCTGATTGTGACG |
| | CAAGTGGTGAACTACGCGCA |
| | GCAGGAACCAGCACTCGCGA |
| *Slc19a3* | CTGTCGGAGTATCACACTGG |
| | TGCCAATTCAAAGAATCTAG |
| | ACCAGAGGGACCAGTAAACA |
| | ACAACGTGTAGCATGATGAC |
| *Slc1a1* | CGACTCACCTAGTACCACGG |
| | TAGGATTACAGCAATGACGG |
| | ATCATGCTGGATACGATCAG |
| | TCACCTGATCAGGTCCAACA |
| *Slc1a2* | CATGTTGATAGCCTTCCCGG |
| | CCATAGCTCTCGTGCCTAGG |
| | TAATTGCCCATAGGTCTGAT |
| | GTTCATGGTTTCATTCAACA |
| *Slc1a3* | GTATAAAATGAGCTACCGGG |
| | GACTCTGACCCGGATCCGGG |
| | GAGGCCGACAATGACTGTCA |
| | AGGCTTCTACCAGATTGGGA |
| | GTCTGCAACCGATTACACAG |

| | |
|---|---|
| *Slc1a4* | TAGAGCCACTCCTAACACCA |
| | GATGCCACCCAGACGCCCGA |
| | ACCCACCAACACTCCCGACA |
| *Slc1a5* | AATCCCTATCGATTCCTGTG |
| | TACAACAGAGTCGTTGATGG |
| | GCGGGAGATCAATTCAACCA |
| | GTGGTGTGCAGCCTGATCGG |
| *Slc1a6* | ACAGCACACGAGTGGTGACA |
| | ATTGGTGGCATGAAGCACAA |
| | CCATGACCCGAGAGCACGTG |
| | GAAGGGAAAGGGGTTCCGAT |
| *Slc1a7* | GTGACAACACATACCCATCG |
| | AGTCCACAGGTAGTATGCCA |
| | CCGGCGGATCGTCATCTATG |
| | GTGATGAGGAAGTACAGTAG |
| *Slc20a1* | GTAGAAAGGTTACCTTACGG |
| | TCAGTATCACACCGTGCACA |
| | CCGGAACGGCTTGATAGATG |
| | GCCACATATTGCCATAGTGT |
| *Slc20a2* | ATACTGTACACAAAGACTCG |
| | GCCTTACCATAGGAAGCCCG |
| | AAGTGGCGATATAAACCAGG |
| | ATGGTAGCAGCATAAAACAG |
| *Slc22a1* | GAAGAAGCCCAAGTTCACAC |
| | TGTTAACACGCCAGACAGGG |
| | CGGAACAGGTCTGCAAACGA |
| | TCTTTAGCTCCCATCTACGT |
| *Slc22a12* | GGGCCTGGGAGTTACATACC |
| | ACGGTAGGCAAGCTGGACCA |
| | GAGGTGCTATTGTCCAGGAG |
| | CATCACCAAAGGGCTACCCT |
| *Slc22a13* | CCTGAAGAAGACCTCCCAGT |
| | AGTAGCCAAGACAAAGCGTA |
| | GAAGAGCAGTAAGACGGGTG |
| | ATCTTGGAGAACACAAGCCG |
| *Slc22a15* | TAGCTCCAATCAGTGCAATG |
| | CCGATCTTACAAAGTCAGCG |
| | CCTCGTTGGTTATACTCCCA |
| | GTCTTGCTAAATGAGTGCGT |
| *Slc22a16* | GCTATTTGTTGAAAACGTGG |
| | TCTTATCACTAGATGTGACG |
| | ATTTACGCTTACCCACTAGG |

| | |
|---|---|
| | TAGACGTAGCCATCGAAGCA |
| *Slc22a17* | TTCTCTCAATGACTCTCACG |
| | GAGGCCCAGGAAGCTTTGCA |
| | GCAGACTCCAGAAACAGACC |
| | ATCGCCAGTCCTTAGAGACA |
| *Slc22a18* | CTAGGTCACTTACCCGTATG |
| | CAGGTTCGCAGACCAGTGCG |
| | TACCTGGAAGTGTGTGCATG |
| | GGCCAGAGATTACCTTGACC |
| *Slc22a19* | ACTGTGGTTGAATATAGCGA |
| | TGTACGGAACAAGTCCAGTG |
| | GTTGGACAGACACTGTTGGG |
| | GTTAATCTGCCATGCCCTGT |
| *Slc22a2* | TTGGTGTTTCGATTTCTACA |
| | AAACGATGCCCACATAGATG |
| | TTCAGTCATTAGTGAACGTG |
| | GGAGACTCCGGTATGCACCT |
| *Slc22a20* | CTGGCGTGGCATACCTGATC |
| | GCTGGACGTACCAAGAATAG |
| | GGTGTGGTCGTACCTGCAGC |
| | GAGCCAGGTCACGAAGGGTG |
| *Slc22a21* | GGGACATATATCCAACTACG |
| | ACAACCCAGTAAAGCCATTG |
| | TTCGGACCAGATCATAAATG |
| | GAAGCTGAATCCGGTGTGCA |
| *Slc22a22* | ATTGGGTGTACCTAAAGCGT |
| | CTAGTCCCCCAAGTACACCC |
| | ATCATGAGGGGATAACACCA |
| | AGGGAACAATACATCAACAA |
| *Slc22a23* | GCCCGATTTCTGGTGCCGCG |
| | TCGAGAAAGAGCTTTCACGG |
| | AAATTGCGACTGCCACGCGT |
| | AAAACGGTTCATAATTACCA |
| *Slc22a26* | ATGAGAACTCATTGACAGTG |
| | TGGAACAAGTCACGTAGAGA |
| | CCAACTCTAATAATAACAGT |
| | ATGGCTGGGTGTATGACCAG |
| *Slc22a27* | TCCCCCTGGATTCCAACCTG |
| | TGTGGTACCTCATATTGCAA |
| | ATACACCCAACCATCCACAC |
| | GTTCCACAGACAATCCTGCC |
| | TGGACAAACCCTAAATTCCT |

| | |
|---|---|
| *Slc22a28* | GGTGGATGTCAGAGTCAGCT |
| | CAAGTTACTATTTACTGATA |
| | TCTTCTTCTTTATACTCACA |
| *Slc22a29* | ATACATCCATCCTACGTAAG |
| | TCCAATATGTGGTACTACAA |
| | GAATTCTGGAGACCTAACCA |
| | AAAGCACTGCCATCATGGCT |
| *Slc22a3* | AGGACACCAGAGGCGCGGAG |
| | AGGAAGAATAATGATCCCGA |
| | CCTCTCATCAAATTACTCAG |
| | CCCATTACCAGTAATAGAGG |
| *Slc22a30* | TGGACAAGCCCTAAATTCCT |
| | GGACCTGGTGTGTGAATCTC |
| | ACACTCACCTGTCTGACAAA |
| | CTACTGCTTACTTCGATTCC |
| *Slc22a4* | CAGAGCAAAGTAACCCACTG |
| | GAGAACGCCTACGAAGAACA |
| | CGCAAAGATGAACAGCATCG |
| | AACCAGGCAACGGTGCTCGG |
| *Slc22a5* | TTTATGATCTGATCCGAACA |
| | GGGTCAGATCTCCAACTACG |
| | CACAAGGCAACGGTGCTCCG |
| | CACACCCACGAAAAACAAGG |
| *Slc22a6* | AAGGAACTGACTCTAAACAA |
| | ATTCACACCCGTGCCTATGT |
| | TCTTCATTGAGTCAGCCCGC |
| | CCCGTAGTAGGCAAAGCTAG |
| *Slc22a7* | CGGAACAGGTCTAAGTACGA |
| | CCGTATCAGGTACCCAACCA |
| | GAGCCTGGGGATAGGCAAAG |
| | TGCATCGGGAGCAGGAATCG |
| *Slc22a8* | TCTGAAGACACTCCAACGTG |
| | GATCTGTAGCAAGTTGTGGT |
| | AATGGGTACCCACCTCCACG |
| | CGTTTGGCAGATGCACGAAG |
| *Slc23a1* | CTGGCATCCTCGTATCCGGG |
| | ACAGGCATAGTAATCACCGA |
| | CTCATCCAGTCCCAACATTG |
| | AGGCTCGAACTGATGCCCGA |
| *Slc23a2* | CTGGCATCCCCGAATCCAAG |
| | GTGTTTCAGTGGCACGATCG |
| | ACAGGCGTAGTAGTCACCGA |

| | |
|---|---|
| | GCGTGCATAGTAGCCATAGT |
| *Slc23a3* | GGGCACTATAGGACTTCTAG |
| | GGAAGGCCCACACACCCGAG |
| | ACCTACCTACCGAAAGGAGT |
| | GACTCACTGTGCCTACGTTG |
| *Slc24a1* | CACACAAGTCCACCGATGTG |
| | AGTGTGGAGGATCGACGACA |
| | GGTGAACACATAGAGAGCGT |
| | AAAGCTATATCCCAAAACCC |
| *Slc24a2* | TCGAAGCCACGCCTCCAACG |
| | TGAGAATGAGAGGCAGAATG |
| | GGATAACGTTATCATGTGGT |
| | AAAGAACTCATCACAGACGA |
| *Slc24a3* | AATCCCAATTAAGCACACAG |
| | GGCCCCAGAGCTGTTCACGT |
| | CAGCAACTGCGATGCCACTG |
| | TCCAATGAAGACCGACGACA |
| *Slc24a4* | TCTCATCCACCATAACCACG |
| | ACCCACGGAGATGTCGGTGT |
| | TCTGTGCAGTTCTTAGCCAG |
| | AAACGGGAGACATGAGAACA |
| *Slc24a5* | CCTTCGGGAAACTCCGATGC |
| | TGAACTAGTTACCGCCTTCT |
| | CAACATCCTGCGACAGTCCA |
| | GAAGTACTTGTCGCAGACGA |
| *Slc25a1* | ATGAACGAGCGAACCCACCG |
| | GAATAATCTCTCTAACCCCG |
| | ACTGCGACTGTACTGAAGCA |
| | CTTCACGTATTCGGTCGGGA |
| *Slc25a10* | CATGCGGGACTACATGACCA |
| | TACACGGTACAGACCATCCA |
| | GTCAGAGAGTAGGTCATCTG |
| | GACATTGACCAAATCTGCTG |
| *Slc25a11* | ACAGACTTAGGGGAGGTACG |
| | ATTTGTGGGAACGCCAGCTG |
| | AAGAACCGGATGCAGTTGAG |
| | GCCTGAAGGGCATTTACACT |
| *Slc25a12* | ACCCGGATGCAAAACCAGCG |
| | ATAGCACTTTAGCAGGCACG |
| | CTCTTGATAGGAGATCAACC |
| | TCACTGGAAGTCTTACCCTG |
| | GGGGCGACTCCCAGTAACTG |

| | |
|---|---|
| *Slc25a13* | CAGATTTATATGAGCCGAGG |
| *Slc25a13* | ACAAGGCATCCGGAGCACAC |
| *Slc25a13* | TACAAGATCGATAGGATACA |
| *Slc25a14* | GATGCCTGTCTTAGTAACGC |
| *Slc25a14* | TACCAGCAAGAAGGTACCAG |
| *Slc25a14* | TGAAACGAACATCGATACTC |
| *Slc25a14* | TTATTTGTAGAACGTTTGGA |
| *Slc25a15* | ACCTTTCCAGACCTCTACCG |
| *Slc25a15* | GCCCATGGTAGAAGCCCAAG |
| *Slc25a15* | GCACAGCATGCGTACTGACT |
| *Slc25a15* | GGACAGCACTTACTTCTGAC |
| *Slc25a16* | CTGCACTTACCCTCTCGATG |
| *Slc25a16* | CCAGTAGAAGTCTCGGCGTG |
| *Slc25a16* | ATGAACTGGATTGCACCGTA |
| *Slc25a16* | ATATAGTAGGCATCAGACCT |
| *Slc25a17* | GAAGTCTAGCAGTATCCAAG |
| *Slc25a17* | GCCTGGCACCATACCGAGGA |
| *Slc25a17* | ACTACAAAGGCATTATCGGT |
| *Slc25a17* | TTAATAGCCTCAAAGCAGTG |
| *Slc25a18* | TGGGACAGGTAATCACCACC |
| *Slc25a18* | TGGGATGTGCCTCACCTAAG |
| *Slc25a18* | AAGGTCAATGGGAAACACGC |
| *Slc25a18* | CCTGATTCCACAGGACGCAG |
| *Slc25a19* | ACGGACCATGTATAAGACCG |
| *Slc25a19* | TCAGCGCACTTTGTGTGCGG |
| *Slc25a19* | GACCCCAATGCCAAATACCA |
| *Slc25a19* | AGAACTGCAGGCCCGCGTAG |
| *Slc25a2* | GGCATCCGTGGCCTTTACAG |
| *Slc25a2* | GACACTTCACAAGCTCAGTG |
| *Slc25a2* | GGTCCCTTAGGCTTCTATCG |
| *Slc25a2* | CCACTTTCCTGACAAACTGT |
| *Slc25a20* | ATAGGGGTGACTCCAATGAT |
| *Slc25a20* | TCCAAGGTCCCAGAGTACAT |
| *Slc25a20* | TCAGGGGAGAACAAGTACAG |
| *Slc25a20* | TCCCAGCTGTAAACAGCTGT |
| *Slc25a21* | CCTGATGCATCCTCTCGATG |
| *Slc25a21* | CAATCGGAACTTGTTCAAAG |
| *Slc25a21* | CCAGATGATCTTCCGAACAG |
| *Slc25a21* | GATATATGTCACTATCACCA |
| *Slc25a22* | GTGTCTTAGCCAGGTCGATG |
| *Slc25a22* | ATACATGCCGAAGTAGCCCT |
| *Slc25a22* | TGATTCAGGTTGGCAAACAG |

| | |
|---|---|
| | GACACCAGCTCTCTAAGGAT |
| *Slc25a23* | GCAACTCGTGCACGTCCACG |
| | CAACCGGCTTAACATTCTAG |
| | GTTATACCCTCAGATCAAGC |
| | TCAGAGACATCTATGTGACC |
| *Slc25a24* | ATGAGAAAAAATCAGGACAG |
| | AGAGACTGGACAATTTCAGA |
| | AAGAAGTTGCTTACCGAGGA |
| | ATCTTTGTTGACATCGCCAG |
| *Slc25a25* | AACAACATGTGCATCGTAGG |
| | GAGGATCCACGAAAGGCTTG |
| | CTTGACACCCAGGTCCCGCA |
| | ACTGGAAGCACTCGACGGTG |
| *Slc25a26* | GGCATTCAAGGACTGTACCG |
| | AACACCCTTACCGTTAGGAA |
| | CCAGCCTTGTTAAATCCCTG |
| | ACTTTCAAGGATTCCCACAA |
| *Slc25a27* | GCAGCGCTTGTGAACATGGG |
| | AGTAGGAACTTGCTCGTCCG |
| | CCTACGTGTCTGTAAATGGC |
| | GCGCACCATGCCCCTATAAG |
| *Slc25a28* | CAACACGTTCCGATAGCGGG |
| | TTGAGTGACGTAATCCACCC |
| | CCCGAACACAGTCTGTCACG |
| | ACGTCACAGCAACAGGCGCG |
| *Slc25a29* | TCGTCTTGGCCAGTTCCATG |
| | TACCGGCATGAGGGCCTGCG |
| | GATGTAATGACCCGCGCCAT |
| | GGTCCTGCCCGTACCTACAA |
| *Slc25a3* | GCCCCGAAGTGAATGTACAA |
| | GTCAGCGAAGAATTCAGCAC |
| | AATACAGTGATGTGCGCCAC |
| | TGCGGCACTTTACTAAGTCC |
| *Slc25a30* | GAGCCTGAAGCGGTTAGCTG |
| | AGTCGGATTAGCAATAGCTG |
| | CAATTGATTTAACTAAGACA |
| | CAGAACAGCGCTGTTCAAGG |
| *Slc25a31* | GGGTTCTGGCAAAATCTAGT |
| | CACAGCTGTCTTCGACACCG |
| | GACTGATTGGTCTATACCAA |
| | AGCCCTGAGGCGCGCTACAA |
| | CGGATCTTCACGAGGTCGAG |

| | |
|---|---|
| *Slc25a32* | GGTTCTCGTACCGGACGTGG |
| | GGAGTAACCCCGAATGTGTG |
| | AGGTGTGCGTGGATTATACA |
| *Slc25a33* | AGTCCAGGCGTCACAGACGT |
| | CTATTTGGATGGTTAAAACG |
| | TGTCTCCGAGATCCCAGCGT |
| | CTCCAACCAAATTTGGACCC |
| *Slc25a34* | GCCGCGTGCTGACGACGTCC |
| | CCCTGGACGTCGTCAGCACG |
| | AGCTTCGGAAACTGGTTGCG |
| | GTGGGGACCCAAGCGCAGAT |
| *Slc25a35* | GCCAGAAACATGGTCTAGTG |
| | TCTGGGAAGCCCAATCTACA |
| | AAGACCAGAATGCAATTGCA |
| | AGAACTGGTACAAGAGCGCA |
| *Slc25a36* | TTGTAGATGTGGTGGCACAG |
| | AGGGCTATCCTGGAAAAAGA |
| | GAGTCTTTATAAGCCAAATG |
| | TATCAGACAGACGGACTGCG |
| *Slc25a37* | CTGGATTCATTACTGCATCG |
| | CACTGTCCGGATACAACTGA |
| | GATGAAGTGAATTGACTGGA |
| | GCATCTATGGCGCCCTCAAG |
| *Slc25a38* | TCCAGGGACACATCTCACAA |
| | CTGAGAAGGGGGCATCACGG |
| | AGAGGAAAGCCTTAATCACT |
| | AGTGATCAAGACACGCTATG |
| *Slc25a39* | GTACTCACCGCTGGTTGCCG |
| | CAAGGTGCCAGTGAACCGTG |
| | AAGTGCCTCCTATACTGCAA |
| | GGCGGACCTTCACCACATCC |
| *Slc25a4* | CAGTTTGACCCTCTCGATCG |
| | GGAAGATCCCTTGCCCACGT |
| | TTGTGTCGTGAGAATCCCCA |
| | GGGGAAGTACCGGATCACGT |
| *Slc25a40* | TATATCCATAGTGACTCCCC |
| | TTTCAACTTACAGAGTAGGA |
| | ATATACATACCTTTGGGGAA |
| | ATGTCTTTACCAATGTCCCA |
| *Slc25a41* | GACGCTCATAAACCCCATGG |
| | CTAGGTCGGTGCAAGCATAG |
| | GAACTTGATAGCATACTCTG |

| | |
|---|---|
| | ACAAACCTGCATGTACACCC |
| *Slc25a42* | GCTGGGTGTCATTCCCTATG |
| | AGGAATCACTCGCACCATGG |
| | GTACGTGAGGGAAGCCGCAG |
| | CTTCATCCGAATCTCGAGAG |
| *Slc25a43* | GTGCTGGAACCATCATACAG |
| | TGTAGGGTATGTGACAATCG |
| | GTTCCTGGCCCTTTATCGAG |
| | TCCACTGGGAAATACGGCCC |
| *Slc25a44* | GTCAAATCACTCGTAGCCGG |
| | GATGTCCTTAGTCTGGCCAA |
| | TGGGATGTACGTTAGCAGCG |
| | TGAGTGTGAAGGTGTTGACC |
| *Slc25a45* | ACAAAGTACATTCCTAACGT |
| | GGCTACCTACTGACTCATGG |
| | TTTGTAGACGGACTTTGATG |
| | TGCACAGGCCCCCGGTACCG |
| *Slc25a46* | GTGAATTTACACCTTTACCG |
| | TGAGATGGTAATGCCGAGCA |
| | GCGGAACCCTCGAGCGTCGG |
| | AATTGGACGAGTGATAGGCT |
| *Slc25a47* | GGGACACGTATCGTCAAGAG |
| | CAAAGGAGTGACCCTCACGA |
| | GTTCCTGACGTCACCCACTG |
| | ACTGCACTGTTTAGTCACAG |
| *Slc25a48* | AGATCAGACAGACTTCGGCC |
| | GAACACCACCGAGTTATAGA |
| | CACATTCAACTGCATCCGCA |
| | CTGGTATGCTGCCACCGCCC |
| *Slc25a5* | CAAGGGCATCATAGACTGCG |
| | CGCAGCCATCTCCAAGACAG |
| | TTAAGATCTACAAATCTGAT |
| | GCACAAGGATGTAGCCCCAG |
| *Slc25a51* | GGGATTGCAGAGTATTACCG |
| | ACCAAACATAAGTGCCAGCG |
| | TGACAAGTCCTGGTTCGAAG |
| | TTACTTGTGTGGCTACTGCG |
| *Slc25a53* | GTCACAGGCCTACACCCTTG |
| | GGCCCTCAGTACTTCTACCG |
| | CTGTCACTGGGCTATTATCG |
| | AAGGGAGTGTGGCCCAACCG |
| | AGAGACAGCAATCAAGATCG |

| | |
|---|---|
| *Slc25a54* | GAATAGCGAGTGCGATCACA |
| | CTCCGTGTCTTCGCTAAACG |
| | TGGCCTGAGAGAGAGAAATG |
| *Slc26a1* | ACAGGCCAACATATGTGATG |
| | CCAGCCGGAGGATACCCATG |
| | TACCAGACCCTAAGATAATG |
| | GCCCACATTAACATGGCGGG |
| *Slc26a10* | CGTGTAGAGTCCGAACACCG |
| | GCTCCCTTTCGATTCCACTG |
| | GGACACAAGATACAATGTCC |
| | AAAAGCAGCCGATCTGGCGT |
| *Slc26a11* | CTTGCCGTGAAGTTCAGCCG |
| | TCCGCTGCCAGCATCACAAT |
| | GGTACCTGGTCTCTCCGATG |
| | AGGGTAGGCGACGCTGTCCT |
| *Slc26a2* | GAGCCGACACCATGACTCCG |
| | ATGGCCGGAGAGCTTTCCGT |
| | ACTGTGCCTTATGATTGGTG |
| | TCCAAAATGAGAAGCCAATG |
| *Slc26a3* | CTATCCGAGTCCCTAATCAG |
| | TCCACGCTGGGTGTAATAGG |
| | CTGGCAACCAAGATGCTATG |
| | TAATCCGAGTCCAAGGACAA |
| *Slc26a4* | TTGTGAATCCGCCAACCAAG |
| | GCTCCCAAAATACCGAGTCA |
| | TCTGGTTCCAGTGTCTAACG |
| | GAAGACGTTGCTTATCCCAA |
| *Slc26a5* | GGTGCGAGGCTTTACCACTG |
| | CCTACATGAGTCCTACAGTG |
| | GATGAGTACAGGCCAAACAC |
| | TGACGATCACAATGGCGGAA |
| *Slc26a6* | ACCGGAACCAAGTTCGCCAC |
| | AGCTGATGATGCACGCGTGC |
| | CCCGGTATCCTGTGCGTGAA |
| | GTCCTAGATGCTCCCGTGCT |
| *Slc26a7* | TAGACCATACTTATTACCTG |
| | TTGTGATGTCACGACATGGG |
| | GCAGTTCAACAGGTGGCACA |
| | TCAGCTCTTTAACGAGAACA |
| *Slc26a8* | TGGGATCATCTTAAGCAATG |
| | AAATGTTAATGACTACCGGG |
| | GCGTAAGTGACGTTGAGGGG |

| | |
|---|---|
| | ACTTACAAGCAAGATAACCA |
| *Slc26a9* | TGTCACCAATGAGACCTACG |
| | CTGACCATTCCCTCCTATAC |
| | CTTTGCCGTTATCAGCATCC |
| | ACTCAGTGGAGGGTGTATCC |
| *Slc27a1* | GGATGGGGTACACATGCGTG |
| | GGCTAGAGCTGCGACGACAC |
| | GAGGCAGCCGGTTCGCGACG |
| | ATACCTGCAGAGTGGTAGAG |
| *Slc27a2* | TGAGCTGATCAAGTATGACG |
| | ATGCGTAGAACTCATACACG |
| | GCTTACGTAAAAGACGGACA |
| | CTTCACGGATGCATCGTGGT |
| *Slc27a3* | CTACGTTCAATTACACAGGA |
| | GCAGCGCTTTAGCTACGCGG |
| | GTGCCCACCGCTTTACGCCG |
| | TGTGGTACAGTGGGAGTGCG |
| *Slc27a4* | CAAACGGATAGGGTACACAA |
| | TCTACACATCGGGCACCACG |
| | TGACTTCAGGAAACATCGTG |
| | ACAGACCCACAAACTCATTG |
| *Slc27a5* | GTGGGCTTAATGAACTATGT |
| | TACCTCTGTACCATACGATA |
| | GTAACAGTGATCTTGTATGT |
| | CCTTTGTGGATGCTTTAGAG |
| *Slc27a6* | GAACTGTACGGAGCTACCGA |
| | AAGCCTTCATCATTTATGAG |
| | CAACTGCCAAACGTACCCGA |
| | GGTGCGCTACGGAATTCAGA |
| *Slc28a1* | ACACCAGCTGTTCCGGCCGC |
| | ACGAACAGCGCCAGCGCCCT |
| | TGGCCTACAATCTGCTAAAG |
| | CGCCCAGCGGCCGGAACAGC |
| *Slc28a2* | ACCGCTTTGCCCAAAGATTC |
| | TCCTCCTCCAGGGTGTTCGT |
| | AAAACACCCACCTGAATCTT |
| | CTGTTGTTGGCCTCATCCTC |
| *Slc28a3* | AGTTCCGACTTGGTGACGTG |
| | TGCAGGTGCGGACATGACCG |
| | AAGGACACGCCAAACAGGTA |
| | TCTGTGGAAGTTTATCGCAC |
| | GCTGATGCAGAAACGAGTTG |

| | |
|---|---|
| *Slc29a1* | GCATGATTGATCAGTGTCCG |
| | TACACAGCCCCCATCATGAG |
| | GGCCAAAATGACAACTGCAC |
| *Slc29a2* | CAGGCTGAGGTAACATACGA |
| | GCAGCATCCCCGAGTCGGTG |
| | TCTGGGCATCCACGCCACCT |
| | GATGAACCAGACGGACGCCA |
| *Slc29a3* | AATGATGGCCATGCACGCGA |
| | GGGAAACTGCGCAGAACCCG |
| | CAAGGAAGACTGCTGCCATG |
| | ACCAGAAAACACTCGAACTG |
| *Slc29a4* | GCTGTTGCACCGATACGTCG |
| | ACTGCCCAAGAGGTACACGC |
| | GTGGTGAAGATAGTCGACAT |
| | TGTGCTCCTAAACAACGTTG |
| *Slc2a1* | CCTGCTCATCAATCGTAACG |
| | TCAGCATGGAGTTCCGCCTG |
| | GTGTCACCTACAGCTCTACG |
| | CAAACATGGAACCACCGCTA |
| *Slc2a10* | CAGCCCCGTGGCGACCAAGG |
| | AGAGGCGTAATACAGCACAT |
| | TGTGCTGGTGTCCCTCTACG |
| | AGCCAACAGATAAACGGCCC |
| *Slc2a12* | GTCCTACACGCTCCTTATAA |
| | ATTGGAGCCCGTTAGGTGTC |
| | ATACCAGTGTCAGACCTATC |
| | GGCGCGGCCTCTAATTCCAC |
| *Slc2a13* | CATGGCCCCCGACACCACGC |
| | CCTCGGTGGCTGATTCAGAA |
| | CACATCCCACAACTAACGCT |
| | AAGGATAATCAGTGCTACTG |
| *Slc2a2* | AGAGGGCTCCAGTCAATGAG |
| | TTACCGACAGCCCATCCTCG |
| | GGACTGGTTCCAATGTACAT |
| | TGTGATCAATGCACCTCAAG |
| *Slc2a3* | TTAGAAGACCTACCAAGTGA |
| | GACCACGCCTGCTCCAATCG |
| | CTGGAATGATGGTTAAGCCA |
| | TGTGCCTATGTACATTGGAG |
| *Slc2a4* | GCAGCCTCTGATCATCGCAG |
| | AACAGAGCTACAATGCAACG |
| | AACCAGAATGCCAATGACGA |

| | |
|---|---|
| | CCAGGTCTAAAGCGCCTGAC |
| *Slc2a5* | CTGGCCCCGAAAAACCTACG |
| | TGCGCTGAGGTAGATCTGAT |
| | CCTGCCCAGTTTATTCACCA |
| | CGGGGACTCCAGTTAGACCC |
| *Slc2a6* | GAGTAGCCGAGTGTCGTGGG |
| | GAGAGACAGGGATCCAAACA |
| | CATCTTCGACAACACATCCG |
| | ACTACACCTGGACAAAATCC |
| *Slc2a7* | CGGCGATGTTGTAGCCATAC |
| | TGCCCCACTTATTGACCATC |
| | GACACGCACTTTGAGCGACA |
| | TTCCCACCAGCACTCGAGAC |
| *Slc2a8* | CCTGCGCCTCGGAGACAATG |
| | ACAGCAAAGCCAGTCACGAA |
| | GATTATGCCCACAGTGACCG |
| | TTGAGTGAGGAGAAAACGTG |
| *Slc2a9* | CGGAGAGGTTGTACCCGTAG |
| | GCATGCCATCAGCAACGCTG |
| | CCAACTATGTGGACTCAATG |
| | TCTCAACGAGATCTCACCCA |
| *Slc30a1* | TGGGCCAGCGTCACGCATCG |
| | CCAGGAGGAGACCAACACGC |
| | GGCATTCACGACCACGATCA |
| | GGAGTCTGACAATCTGGAAG |
| *Slc30a10* | CCATGACAACAGTCAAACTG |
| | AGCCGTGATGACCACAACCA |
| | GCACAGCAGTGACTCTCCGG |
| | TGCTGCAGATGGTCCCCAAG |
| *Slc30a2* | CTGGTCGGGAAGACACCCAG |
| | TGGAGATTATGAGATCAAAG |
| | AGGCCACATAGAGTTTGCGT |
| | GATGGAAAGCACGGACAACA |
| *Slc30a3* | GATAGTCACTATGAAGCAGG |
| | GTCTCCCAGCACATGCACAA |
| | GCCCACTTGCTAGCAGACAT |
| | TGCAGAGTATGCACCTCTAG |
| *Slc30a4* | CGGACGAGGTGAGCGACGAG |
| | TCCACTAGATACCATGCTTG |
| | ACCTTTGGATTTCATCGCCT |
| | ACTTTGTACCAAGTCTCCCA |
| | TGTTCTTTCTGTGACGACTG |

| | |
|---|---|
| *Slc30a5* | GGCGTGCTAACCAACAGTCT |
| | ACCACTCTATCACTTCATGG |
| | ACAGAGCACGTCCTGTCTGG |
| *Slc30a6* | AACTGACCGATAACATGAGG |
| | TCAGAACAAACGGATTCATG |
| | GGGCAACATACCGATGCTGT |
| | CTTACCCAAATGAATAGACA |
| *Slc30a7* | AGGAGTCGGAGATCAAGCCT |
| | CATTAAAGAGGGAATGACTG |
| | AAACAAGCAGCAGTCTCTCG |
| | CATGACCATGACCACGCTCA |
| *Slc30a8* | GCTCTGAAATACATCCCCCA |
| | TGGTATGACTTACACAATGT |
| | TCTGGCTATCCTCACTGATG |
| | CTTGCTCGACCTGTTCCCTG |
| *Slc30a9* | AAGGACTCGGTGTCATCGTG |
| | ACTGAGACCGCTCTGGAACG |
| | ATTATCTGATACTTGTAACC |
| | CTACGAATGTCCAGAAAGGA |
| *Slc31a1* | TGTGGTTCATACCCATATGG |
| | TTGGTAATCAATACACCTGG |
| | GGACTCAAGATAGCCCGAGA |
| | TCAGCCTCACACTCCCACGG |
| *Slc31a2* | CTGTATGAGGGCATCAAGGT |
| | CAGACAATAGGACCCGCCTC |
| | GGACCAGACCAGGATTCTAC |
| | CAAGATGAACTGCTGGCTGT |
| *Slc32a1* | GACGTGTATCTTGTACGTCG |
| | ACAAACCCAAGATCACGGCG |
| | GGAGACATTCATTATCAGCG |
| | CAGCAGACTGAACTTGGACA |
| *Slc33a1* | GTGACTTACCTAAAGCCCCG |
| | ATGTTATCCCGGGAAAACGT |
| | GAAAGGGTAACGATTCCCCT |
| | AAATATTGATGGCAGAACAC |
| *Slc34a1* | AGTTGAGCATCTTCACAAGG |
| | TGACCCACTACCTACCAAGC |
| | GTGTCACCCAGACACAACAG |
| | TGCCATCCTATCCAACCCAG |
| *Slc34a2* | TCATAGAGGAGCATCCCGAG |
| | CTCCATCACCAACACGATCG |
| | AGAGGTGCAGTTATCAGTCG |

| | |
|---|---|
| | CTCACCGATGAGTGGAGTCA |
| *Slc34a3* | AGGTTAGGCCTGCGCCAACG |
| | AGTTGAGCAGTTTAACGATG |
| | GATAGCAGTGTGATAACCAG |
| | CAATGACCAAGCCCGCCACA |
| *Slc35a1* | GAACACTCAGCAAATTACAG |
| | ATAGCACCAAAGCCTAACAA |
| | TGCACAGCATACACTAGTGA |
| | TCTTAAAGCTACGGTGTAAG |
| *Slc35a2* | TCACCCGCTGTAGTGGACCC |
| | CTGCTCTTCGCACAAAAGAG |
| | CTGCAAGGTATAGATGAGAG |
| | GGCCACTGGATCAGAACCCG |
| *Slc35a3* | GCTGTAGAAGACAGATAACG |
| | TGAAGCTCGCTATCCCGTCA |
| | TCCTGCCATCAGAATTACTA |
| | CAAACTGTGAGCCTGTTGAA |
| *Slc35a4* | TGTTTGCCAGCCTACCAGGA |
| | AGTGGACAGGAAGAGCAACA |
| | GTTCTTAGCACTGTGCCATG |
| | TCCCAGTGGAGTGATATGCA |
| *Slc35a5* | ATTGAACTATAATCAGAACG |
| | AAGAGACAATTGTACATCGA |
| | GATTCTGTTCTTGTCTATCG |
| | CTCTGAGCACACGTTCACAG |
| *Slc35b1* | CCTGTGTGTATTGCTAATTG |
| | GTTTGTCAACTATCCAACTC |
| | TGTTTCGTCTTAGCACAAGA |
| | GGAGACCATGGCACCCACAT |
| *Slc35b2* | AAGCAATATAGCCTGCCAGT |
| | CTGCAGGAAAGAGTGATGAC |
| | GGCTCCGCGGACAGAGACAG |
| | TACATGGGTGCACCATGACG |
| *Slc35b3* | GTACACAAACCCGATTGAGT |
| | TCCTCAACTGACTTGATGTG |
| | AAGGTAAGGTACCAGCCGTA |
| | TCTTGTCTCACATAGGAAAG |
| *Slc35b4* | GATCTGAATATCATATGCAG |
| | CACGCTGACAGTGAAGAACA |
| | GCAGTTCTTATTTATTGCTG |
| | GAGGCCAAGAAGATGAACCC |
| | AGGGCACCCCTACGTACTTG |

| | |
|---|---|
| *Slc35c1* | AGGTTAGGCGCCAGATACTG |
| | CTCACCAATGATGACGCCGC |
| | GCAGTGAGGTCACCAGGCAT |
| *Slc35c2* | CGAAGGCCAGAATCCCACCG |
| | GTATTTGTGTAAGGGTCCAG |
| | CCTTCGAATATGGCAAAGAG |
| | AGGTGGAACATGGTGTCAAT |
| *Slc35d1* | GGCGCCGGCCGAAACGCTAA |
| | TGTGTGTGGAGGATTTCGCG |
| | ACTGTATTTGCAATGATCAT |
| | TGGTTCCCAAAATATAGTAG |
| *Slc35d2* | AATTCCAAGTACAATTGGTG |
| | TTCCTGAGCACGGTGAACAT |
| | CTTGAATGATATCTTCACCG |
| | TCCTGGAGGCTATCATACTT |
| *Slc35d3* | CGCCCGTTACGTACCCAATG |
| | CTCACTCTCTGGTCGCTGCG |
| | GACGGCGATCACATACTGTG |
| | CGAATATGGAAACCATGGCC |
| *Slc35e1* | CAGAGCTGTCCTTCGACGTG |
| | TGTTCTTCATGATCCCCACG |
| | GACCAGCCCCCACACGTCGA |
| | CTGGGTTGCGCAGCATAATG |
| *Slc35e2* | TGCGTTATCCACCAACATTA |
| | ATGCTGTCAACAACGTTAAT |
| | ACTAGCGTTGTGCACTGCCA |
| | ATGATTCTGGGGGAGTACAC |
| *Slc35e3* | CGTCAGAATACAGCTCACGC |
| | AAATGGATCTATGTACACCA |
| | AGTCACGTCCCTTTATCAAG |
| | CTGGTAGTAAAGTAGCTGCA |
| *Slc35e4* | TGAGCAGCAGCACTCGACGG |
| | CTTGAAGCCTCGCAGACAGG |
| | GCTGCCACAACCACTCTCGC |
| | CAAAGAAGTGGCATACAGCA |
| *Slc35f1* | TTCGTCGTAAAATTGCCAGG |
| | GACATTTGAGATCCCGTACA |
| | ACCGGCCCCGCCGAACCATG |
| | AGGAGAGCAAAATCACCACC |
| *Slc35f2* | GGCCGTGCCGCAAATACACA |
| | TTGGTGCAGACATATTAGCT |
| | TCTTGCCCGGAGAATAAACC |

| | |
|---|---|
| | CCAGCATCAACGTGTAAACC |
| *Slc35f3* | GTCATGATGACCTATGCCGA |
| | GGTCCTGACCCTCACCAAAG |
| | GTTGGTGGCAAACCACGTCA |
| | ACTGGACATGTTCTCAGGAG |
| *Slc35f4* | GATGGCCGGAATAATAGACT |
| | TCAGTTACTCGATGCAAACC |
| | CATTGTCATGATGGCGTATG |
| | ATGGTTCTGAAGACCATCTG |
| *Slc35f5* | GCTGTATTTCCGAGTAACAG |
| | CAGGAGAATCACTATCCCCA |
| | TCAGAGAAGCCCTTGCAACA |
| | CTCCCAAGAAGTCCCGTGTA |
| *Slc35f6* | GGGACCAGCATCATGTACGT |
| | GGATGTAGAACATAGGCACA |
| | AATGCTGGTACCTGGACGAA |
| | GATCACTATTGCGGGACTGG |
| *Slc35g1* | ACAGCTAAACGCAATAACTG |
| | ACGCACTGATCTCTACAGCA |
| | TCATTTAAATTTGCAGAACG |
| | GTCCACGTGCTCCTCGCCGG |
| *Slc35g2* | GAAAATGGCGTATGTTGACA |
| | CTTGGGTCTTCTGACGACGA |
| | AGTCGTAATCTGTATCCGCT |
| | TACTATCATAGAAAGCGCCG |
| *Slc35g3* | CTTGGGCTAATCATCATCGT |
| | GAAAGCCAGTACATAGCCCA |
| | ACCACGGAACTTAAGTAACA |
| | TGTAGGCGCAACCAATGCTG |
| *Slc36a1* | AGAAGATGGACAACACACGC |
| | GTCCGGAACCACTCCCACTG |
| | CTGTACTCTACATCAGCCTG |
| | ATCGCTGTGCCAAAGAACAG |
| *Slc36a2* | TGACTAGGATGTGCATACAG |
| | GGTAGTTCTGACGCCCACCA |
| | AGCAAAGTACCTCCCCCAGT |
| | AGCCCCGATAGCAATGTACA |
| *Slc36a3* | TCCCCAGATACAGCACAGCG |
| | AGAGACCACAATGTACAGCC |
| | TGAGCGCCCCGATGGCCAAG |
| | CACCTGTTGAAAAGCAACAT |
| | TTTCGATGGGTCGTCCGACG |

| | |
|---|---|
| *Slc36a4* | TCAGGCATTGAACATCGGCA |
| | CCGCGTTCTTTATAGCCAAG |
| | TTGCCATGGAGGCTAGTCCG |
| *Slc37a1* | CTGAACAATGAGACCGACTG |
| | GTTCCCCTATTATTCCACTG |
| | CCACAGCCTGGGATTCTACG |
| | ATCATGCTGTGTTGTAGGCG |
| *Slc37a2* | TTAGCACCATTCCAGCCGAG |
| | ACGATCTCAATGATACCACC |
| | CCCACATTGAAGATGTACAA |
| | GGCTTGTGCAGACTACAGGC |
| *Slc37a3* | ACCATACTGGAGAACCGATG |
| | TCAGCGGCATCATAGGCGAT |
| | GCCTAACTATTCGATCCAGG |
| | CCTAGGTCTCCCGAGTATCG |
| *Slc37a4* | CTACGTTGACCAGACCAACC |
| | TCTTTACTCCGAAGACCACG |
| | CACAAACTTGCTGATGGCGT |
| | CAGAGCGATCTCATCCACCA |
| *Slc38a1* | ATACTTTGGTGTGCACGCGT |
| | TGCATGGTGTATGAGAAGCT |
| | TCACCATCACCACCAACACT |
| | AGATTGGCAGGACGGACGGG |
| *Slc38a10* | TTGACGCAAGAAGCTCGCAG |
| | TGCACGGCAGAATCATCATG |
| | GCTTCACAGGAGCAACGATG |
| | TCACCTGTAGCAGCACGGCA |
| *Slc38a11* | TTTCCCTCCGTGTTCATGTC |
| | TGAAAGGGTACATAAACTGT |
| | AATGTAGTCAATTCTGTTAT |
| | CCTTATTCAATGAAGCAAGC |
| *Slc38a2* | CCACCAAAGCAGCTTCCACG |
| | CTCAAGACTGCCAACGAAGG |
| | GCAGTGACAATGGAAGAATG |
| | GAGTTGAAGATGAAATAGCG |
| *Slc38a3* | AGCTGTCGCATCAGTGCTAG |
| | CATGACATACATGACAGCAA |
| | GATGTATAGGTAGCTGGACA |
| | ACCTGCTCCTCAAGTCTTCG |
| *Slc38a4* | CCACGGACACAAATAAGACG |
| | GAAGAAGCTAGCCGATTACG |
| | GATCTCGCTGCCTAATGACT |

| | |
|---|---|
| | GATGAAGAGGTAGCTTGACA |
| *Slc38a5* | AGCTACAGGCAGGAACGCGA |
| | ACCTGCCGGGAAAGTAGTCG |
| | GGAAGGTGCCAATAACAAGG |
| | ACCTCAGCAACGCTATCATG |
| *Slc38a6* | CGTGACGTCTTACGAAGATC |
| | CCTTGGTTTGGCGTATGTGA |
| | CAGATCTTCGTAAGACGTCA |
| | CCATCACATACGCCAAACCA |
| *Slc38a7* | GCTGGTCCCCAATAATGATG |
| | AGTTGAGCAGACCCGCACCG |
| | AGATAAAGAGATGCGCCCGG |
| | ATGTGGTACCTAAGAGTGAG |
| *Slc38a8* | TCACGGTGCAATACTACCTG |
| | ATTTCCCGAAGTGCTGACAG |
| | GCTGGTCCCCGATCACTCTG |
| | TCTTTGGTCTTCCTGATCAG |
| *Slc38a9* | TACTCACATAGTAACTAAGC |
| | AAAAAGGCTGGATTTACCAC |
| | AAACTCAAGAGTTACACTGA |
| | ACTTTACTGCTGCTATAGAG |
| *Slc39a1* | AGCACTGGCACGATGGACCA |
| | CCAAGATGAACTCTTGCAAG |
| | GGTGATGGAGCAGATCACGC |
| | AGACCAGGACACAAGCACGC |
| *Slc39a10* | CGTGCGTATGCTGATGACTG |
| | TGAACAATATGAGCATAACC |
| | AGTCGTTGAGATTAATCACG |
| | CGATCTGATACAGCAATGCA |
| *Slc39a11* | AGAATGGCGAGGTATACCAG |
| | CCACTGTGAACTCACCCCTG |
| | GAGATGGCGACATCCTCGGG |
| | CAGACTTCTTCATCAATGCA |
| *Slc39a12* | CAATCTTACCAGATAGGTGG |
| | ACCAATACCCTCCACCTATC |
| | GATGCACTGTTACTCACAGC |
| | TGAACATGCTCACGACCAGA |
| *Slc39a13* | ATAAAGAAAGCGAGTCCTGG |
| | TACACCTGTAACATCACCCC |
| | CTCACCTTCTGACTGTAACA |
| | CTGGGGCTATGGGTCATCGC |
| | GAGCGAGCGATCTCAGATCG |

| | |
|---|---|
| *Slc39a14* | GTAGAGGGTTCCAATCGCCA |
| | TAAAATGGTTATGCCCGTGA |
| | GTGACCGAGAAGCTACAGAA |
| *Slc39a2* | TCCATAGGGATACTCCACCT |
| | TTTATTAGGTCATCACCACA |
| | TGGATTCCACAAGCATCCAG |
| | CTGTATGGTGGCCGCCACTG |
| *Slc39a3* | CAGCGCACACCCATGGCGCG |
| | GAGTACGAGAGCCCGTTCGT |
| | GCTGCCTGTGAAGGTCATCG |
| | CATCAGCACCGACTACCCGC |
| *Slc39a4* | GGTCCTGAATACGGATAGTG |
| | CAGTTGGGGAAGATCTACAC |
| | CATGCAGCGTGATATTGGGA |
| | TGGAGAGGGTCACACCCATG |
| *Slc39a5* | CAGCAGAGCGAACTGACGAG |
| | AGGACCTAGTGAGCAATCAG |
| | GTGGAGACAATTTCACACAC |
| | TCAGATGGTCAGCCAACGAA |
| *Slc39a6* | CGTGGTCCGAGTGATGCTCG |
| | AGGAATCATTCTCTCCGTAG |
| | CAGCCACGGAACTTACGTGT |
| | GACAGCGTTGTATACCGCCG |
| *Slc39a7* | TCACGTGAGGAATTACACCA |
| | TCACAAATTTCTCCACCACG |
| | TCACATGAAGATTTCCACCA |
| | GGTGGAGGAACGCATCACCC |
| *Slc39a8* | TCAGCTGCTGTAAGATCGCG |
| | AGGGGGTTAAAATCAATCCC |
| | CGTTAGGCTCAGTGACAGCG |
| | CGGCGCCAACCGGAGCCTGT |
| *Slc39a9* | TGCACTGGCGGTCATCGTCC |
| | TTGCCACGAACACAATTAAC |
| | TGTCCAGTTAATTGTGTTCG |
| | TTTCTAGGAGCGGCTGAAGC |
| *Slc3a1* | CCATATACCAGATCTACCCG |
| | ATCCTTGGTTCCAATCGAGT |
| | AGAGGAGCCTCACCTAAAGG |
| | TGGCAAGCCATAGTACATCA |
| *Slc3a2* | GTTCACCGGCTTATCCAAGG |
| | CGCCCGAACGATGATAACCA |
| | TATCACCAAGAACTTAAGTG |

| | |
|---|---|
| | GTACTGAATCCCTAGTCACT |
| *Slc40a1* | CAGGGTACGCCTACACTCAG |
| | CCTTTGGATTGTGATCGCAG |
| | TCATCAGGATGATTCCGCAG |
| | CCCATCCATCTCGGAAAGTG |
| *Slc41a1* | TGAGTCCCGAGCTAACGCCA |
| | TCCCCAGGTACAAGCCACGG |
| | CTGGCGATACATCTATCCCC |
| | TCATTGGGTCTCGAAAGATT |
| *Slc41a2* | CCCGCGCTTCTTGGTCTTAA |
| | TAACTCTCGCCATATTAGCT |
| | TCCTTTAAGACCAAGAAGCG |
| | CTTCATCGCATCTCTACTGC |
| *Slc41a3* | GGAGTTCGATTGGTCCAAGG |
| | TTTCCCTTCAGACCGACGAG |
| | GGTGGATGCCAAATCACTGG |
| | TCTGCATAGTGATTGGTGCT |
| *Slc43a1* | GGATATCCCTGGTACCTCAG |
| | CTGACCCACAATGGTTACAT |
| | CTTAACGTTTACCTCACTCA |
| | CGGTCCATGAGAATTCCCAG |
| *Slc43a2* | GTCACTAACAGCACGGTCGG |
| | GCTTTGACCACAAGATCACA |
| | CACACTGTGCATAAACGATG |
| | GGAGGTATAGAGGGCAACTG |
| *Slc43a3* | TGACCGCTTCAAGACTACTG |
| | GATTCATCTTGCACGTGGTG |
| | TGCTCCAGAGCAATGTAACA |
| | TACCCATAGCTGTAGTTGGG |
| *Slc44a1* | CTGGAAGCAATACCGAACAG |
| | GTACATGTGGTGGTACCACG |
| | GTGGCACGGGTGTATTATGG |
| | CACCATCGCCTTGTTCCACG |
| *Slc44a2* | ATCAACAACCTTGTACACGG |
| | GAACATTACAGATCTAGTGG |
| | GGTGAAACGCATTACCTGAG |
| | TCAAGTGCAGGTACACTCGG |
| *Slc44a3* | AGGATGGCACACGTCCACAG |
| | GTTCATCATGGGTTATTCGG |
| | GAACCGGAAGGCGAACAACA |
| | CCACCACATATACCGAATGC |
| | TGGGCACGAGGAATTCAACG |

| | |
|---|---|
| *Slc44a4* | GATGTCGAAGTATAGAACGT |
| | TCTCCATCCCCAGATAGTGG |
| | AGTTGGTAGTGAAGCCCAGT |
| *Slc44a5* | TTGAAGACTATGCTACCTCG |
| | ACTGGGTAGATTAACGCACT |
| | TCAAAGTTATGGTCCCAGCG |
| | TGGCCATACTCACTCATTGG |
| *Slc45a1* | GAGTACCGGAGTCACGTACG |
| | AGGAGTACTCACCCGCCATG |
| | GGAGTGACCGATGTACCTCA |
| | CCACCCCGCACACTGTCAGG |
| *Slc45a2* | GCATGTTTACTAATGCCCGA |
| | CTGGGCCATAAGCATCACCA |
| | GGCACCCAAAATGTAGCCAA |
| | GGAGCAGGAATCCCAAGATG |
| *Slc45a3* | TGGTACAGAAGTTCGGCACA |
| | GAGGCTTAGCAGGACACCCA |
| | CAGCCACAAAGAGTCGGCGT |
| | AGCCACTCTGTTTGTGACGG |
| *Slc45a4* | AGGTCGCTCATGCTTCGGGA |
| | CGTAGCCAATGGCCCCACCA |
| | GCATGACCCATAGGCGTGTG |
| | GGAAAAGTGCAACACCAATG |
| *Slc46a1* | AGAGCTAACATCTGCCACAG |
| | GGGCAATGGATCGATGATGG |
| | TGGACCAGAAGAGTCCCACC |
| | GAACTGTGGGAACCAAAGCG |
| *Slc46a2* | GATGGGCACCTATCGAACCC |
| | CATGACCCCGGACCAATAAG |
| | AGCCCAGGACTAAATCGATG |
| | GGAGTAGTCCTGCGTCGTAG |
| *Slc46a3* | CAGCGCAGTACGTGTACCGG |
| | TGCTGGAATGAGGTTTACAT |
| | TTAGCGAGCAGCGACAACCA |
| | CAGGTTGACTATAAGAACCA |
| *Slc47a1* | AGCCAGAACTTAAAGCACGT |
| | GAAGATCATGACGTAAGTCT |
| | GGGCGCAGGAAACATTGACC |
| | GCGATGCCCGTCATAATCTG |
| *Slc47a2* | CGTCCAACTCGACTTTGCCC |
| | TACGGAAACAGCGAGTGTCA |
| | AGGCAAGAACTTGAAGCGCG |

| | |
|---|---|
| | GGTGCTCACCTGGCGACATC |
| *Slc48a1* | CGGTGGTCTACCGACAACCG |
| | TGTACATGCAGGATTACTGG |
| | GCTGGGTGATGGCCAATGCC |
| | TACCGAGCTGAAGCCGGAGT |
| *Slc4a1* | CTCCATGGCGCATAACCGAG |
| | AATAACCTGGAGTATATCGT |
| | TCTACAACAGACTTGAACGG |
| | CTTACCCACTAGCACCAGTG |
| *Slc4a10* | AGAGAACATCTGGCACGTAG |
| | CACGATGCCTGTGACGACGA |
| | TGAGATTTGCTGGCGTGAAG |
| | TTCCTGATACCAAGTCATTG |
| *Slc4a11* | GGTCCGTGCACACCGGGACC |
| | CAAAGCGGTTTAGCATAGTT |
| | GCTCTTACACACCTCTCGCA |
| | GAGTCACTGCCACTGTCCGA |
| *Slc4a2* | GGAAGTCACTTAGGTAGTGG |
| | GGGTACGGCGACACTTGGTG |
| | TAGCGGATGATGGATATGGT |
| | GAGCCCGCTGAGATGTTTCG |
| *Slc4a3* | CCGAGACCTACTACGTTCGG |
| | TGGAGCTTGACGGATTTCGG |
| | CCACCCACACTCGACCGGTG |
| | CTCACCCACAAGCACGACAG |
| *Slc4a4* | AGCCTGCTGTAGGCGAACAA |
| | GCCTCCAAAAGTGATGGCGT |
| | ACTTTGAAGTCAGAGTTGAT |
| | AGAGAATGTTCAGATGAATG |
| *Slc4a5* | AGCTATGCACGAAATCGGGG |
| | TGGGGCTGAGGCATTGACCG |
| | TTACCTTGGTGGGAAAATAG |
| | CACCTCGCCCACGAGCACGT |
| *Slc4a7* | TCAGGAACATAAGGTCCATG |
| | TCTGCAAAGGATCGAACCAG |
| | ACAGTATAGGAAAAGAATCG |
| | GCAGATCCATTAGGAAACAC |
| *Slc4a8* | TCATGAAAAAAATTCCCACG |
| | AACGACTCAATCGCACTCTG |
| | ACAGCGTGAGGGTTAAAGTG |
| | CCTTCCAATACTGTAAGGTG |
| | TAGGGGCCTTCGTAAGACTG |

| | |
|---|---|
| *Slc4a9* | ACAGACTTCGAAGCTTCTGG |
| | GTATTGACAGAAGCAACCAT |
| | GAGGTGACAGCTCTACCCCC |
| *Slc50a1* | TTATCATCGTCAATAGCGTG |
| | ACTCACCAAATCAGCCAGTG |
| | ACACTCTCACTTGACATCCG |
| | CTGAGTTACGGAGTCTTGAA |
| *Slc51a* | CACTGAAGGACACCCCGATG |
| | TGGTGAGGGCTATGTCCACT |
| | TCTACAAGTGTGAGGGCGCG |
| | TCAGAGTCCTCTTCTTGATG |
| *Slc51b* | ACCAGGATGGAATAATTCCA |
| | TCCTTGGAATTATTCCATCC |
| | TCATCAAGATGCAGGTCTTC |
| | GCATTTCTTCCAGCAGTTCC |
| *Slc52a2* | TGTTGTGGGTTCAGATGTCG |
| | ATGGGAGACACCTCGATCGG |
| | GGCCTCTCTGTGGAACCACG |
| | AAGACCGTAAAAAGGGGGGT |
| *Slc52a3* | GTGACCTCCTGGATACAGGG |
| | TGTCTCCGTGACATTGACAC |
| | GCTGGTGACTGAGTTGCCCG |
| | GGGTCGGAAGCGGTGCATCA |
| *Slc5a1* | AGGAAGAATGCTACACACCG |
| | GAGACATGTTCTTGGCCGAG |
| | CATCGCCTACCCCACGCTCG |
| | CCGGCCACCACACCATACTT |
| *Slc5a10* | ACAGGACAGATAAGTACGTG |
| | CAGGCTCGCAGACACTCGGA |
| | AGCCGGCCAATCCTACGAAG |
| | GGTTCTTACCCATATGCCCA |
| *Slc5a11* | CGATGCCAGAATATCTAAGG |
| | GCGATGTCTGAACAGCCAGA |
| | ATGTGGAAGGCATCTTCTCG |
| | CCTGCACAATGGGGATCCAG |
| *Slc5a12* | AGGTACTCAGTTTGTCGGAG |
| | CATTATCTACATCGTACAGA |
| | TCTCGGGAATTCTTAGTAGG |
| | ATCCAAGGATCAAATCATGT |
| *Slc5a2* | ATGATTTATACTGTGACAGG |
| | CTGGCACAAAAAGCCATCCG |
| | GGTCTCTTCGACAAATACCT |

| | |
|---|---|
| | ATTGGTTCTGAACATAGACT |
| *Slc5a3* | CCCCTGACCGGATGTAAATG |
| | CTGACCAAGTCATCGTACAG |
| | GAACACTGCATGCAAGTGTG |
| | CTGTGTAGATCACTGCAACA |
| *Slc5a4a* | CTGACTTACCGAATACCATG |
| | GTGGTACTGGTGCATAAACC |
| | CCCCAGCCTTGATGTAAATG |
| | TCTGAGTGTGTGAAGCATTG |
| *Slc5a4b* | TCAATGGTGAGAAGCAACCG |
| | GATGCAGGCGGCCTTCACGT |
| | AAGAACCAGCACCATAAGCA |
| | ACACTCAGAAGGTACAACGC |
| *Slc5a5* | CTGCACCTTGTACACGACCG |
| | GCTGGCCGCTAGCTTCATGT |
| | GTCCACCAGTATCAACGCTA |
| | GTACATGCCATTGCTCGTGT |
| *Slc5a6* | GGAATCCTTACCTGCATTGA |
| | CATCCTATAGGTGATATACA |
| | CTTACTTAATCCCAGAGATG |
| | TCCGCTTCAATAAAGCAGTT |
| *Slc5a7* | CACAGCTGTGAATCCGATGT |
| | CTTGGGATCTGGATACCCGT |
| | GCCCAAGCTAGACCACAACC |
| | CATGGCAAGCCTACTTCCAG |
| *Slc5a8* | CATCTATTACGCCTTCGCGG |
| | CGCCCCAAAACGGTAGACCT |
| | TCTGAGAGAGACTTAAAGCG |
| | CAGGCATCAATAACTCAACA |
| *Slc5a9* | TACCTGAAGAAACGATTTGG |
| | TGAGAGCCATAACCAGCTTG |
| | AAGAATCTTTCACATGCCAA |
| | CAGACTGTGATCATGGTTGG |
| *Slc6a1* | CACCAACATGACCAGCGCCG |
| | GCAGAAATACACGAGCACCC |
| | TACCTCTGTGGGAAAAACGG |
| | TCCATGTGTCCCGGTCAGGG |
| *Slc6a11* | CTCCCAGAACTCCATGACCG |
| | AGGCATTGGCTATGCAACAC |
| | GTTGTTGTAACTCCCCAGAG |
| | GGGTACTAAGTCGACTGGAA |
| | CTGAATCACTCATCGGCCAG |

| | |
|---|---|
| *Slc6a12* | TCTTGGGCCTCATGTAGGTG |
| | GATGGAGTTTGTGCTGTCAG |
| | GGGAATACCCATTTCTGAAG |
| *Slc6a13* | CTTGGCCAGTACACCAACCA |
| | GTAACATCACCACAGCTCGA |
| | ACCCCAACATCACACGTCTG |
| | CCAGAACTCGATGACAGGGG |
| *Slc6a14* | CTTCTCATTCTGTTAATACG |
| | CAAGAAGAACAATTTGCCCA |
| | AATGCTAGCATGATTGCATA |
| | TCAGTAAAGTGACACTTCAG |
| *Slc6a15* | TTCCCGGTACCAGTAATAGG |
| | GAAACATATAAGGACCACGT |
| | TTGTTCCAAACATGCTGCCG |
| | ACATAAGCCCTAAATTGGGT |
| *Slc6a17* | CACGAGGGTGGCCAACACCG |
| | GGCCTCTCGGTACCAAAAGT |
| | TGTAGTACAGCCCAACGAAG |
| | GAACATGACGGACCAGAACG |
| *Slc6a18* | AGAGCGTACCTACGCCACCG |
| | GTGTACCTGTGTGTCATCAG |
| | GAATGCCACTCAGACTGCGA |
| | TTGCAAGCTACAACCCACCC |
| *Slc6a19* | ATCGGTCAGAGGCTACGCAA |
| | ACAGTCATCAAAGCGCTCAG |
| | AGCTGAGCAACCCCAACACG |
| | TCCGTGGCATCGAGACCACT |
| *Slc6a2* | CTTCACCACTAGTAGATCGG |
| | AGCAGTGGGATCCATGACAT |
| | AGCCGACATAGAGGGCAATG |
| | ATAAGGGAACCGCCACACGT |
| *Slc6a20a* | TACATGTTCACACCTAAGGT |
| | AACACAGTAAGGCATCGATG |
| | GACAATGCTGGCAAATATGG |
| | CAACCACACAGGCTACGATG |
| *Slc6a20b* | CCACTGAATAGTAATCACAC |
| | CTGTGCCAGATGTATGGCGG |
| | GATGACATTGAAGTACACAG |
| | CACAAAGTAAGGCAGTGACG |
| *Slc6a3* | TAGATGATGAAGATCAACCC |
| | GCTCGTCAGGGAGTTAATGG |
| | CAGGGAGGGTGACTCCACGC |

| | |
|---|---|
| | TTACTCAAAATACTCAGCAG |
| *Slc6a4* | AGTTCCCAGTACAAGCGCTG |
| | AGGAGTCAAAACGTCTGGCA |
| | TACATATGCTACCAGAATGG |
| | CCGCATATGTGATGAAAAGG |
| *Slc6a5* | GGGATGGTCACCGATAACAC |
| | GTGTACGCATCACTGGCGAA |
| | GCTTACCGGTATGGTAGTGG |
| | GAGGACGCGAACGTGAGTGT |
| *Slc6a6* | GGCCCCCAGGCAGATTGCGT |
| | GGCCAGTACACATCAGAAGG |
| | AAACAGAATAGACCAAAAGG |
| | CATGGCAAACGGGAAAGTAG |
| *Slc6a7* | TCCAGCGAATCTCCCCCGGT |
| | TCACCTTTGAAGATAACAGG |
| | GTAGACTTCGCAGCAGACAG |
| | TACCGAGCCTACACCAATGG |
| *Slc6a8* | GGAAGCGCCACACGTTACCG |
| | GCATCAGTGTGACAGCCCGT |
| | GCACAACGAGGACCACGTAG |
| | ACTCGATGACAGGGGACCGG |
| *Slc6a9* | ATACCTCTGCTATCGCAACG |
| | GTAGTACATGATACCCGTGA |
| | ATGGTGGTGTCCACATACAT |
| | TGTGCTACCAGCGTCTACGC |
| *Slc7a1* | GCCATGGCATAGATAACTCG |
| | CACAAACGTGAAATACGGTG |
| | TGACGTGAGAACTCTCCGAT |
| | CCAGGTCCTTCAGTTCAAAG |
| *Slc7a10* | GTGTACACAAAGGTGACCAG |
| | TGATCCCTCCAAAAGTAGAG |
| | TGAGCCAATGATGTTCCCTG |
| | GGCCATGGAGAGTACTCGAG |
| *Slc7a11* | TCATTACACATACATTCTGG |
| | GAAGAGACACAAGTCTAATG |
| | GGGCTACGTACTGACAAACG |
| | ACAGGCAGACCAGAAAACCA |
| *Slc7a12* | TGATGATCCTAGATATGAGT |
| | ATTTACCTGCTATGACGATG |
| | TGGCTTCAAGCTAACCTTAA |
| | ATTCTACTCTGCAAGTCAAG |
| | GTGGGAATTCTGAATTCTCG |

| | |
|---|---|
| *Slc7a13* | GGAACAGTCCATGTGAATTG |
| | AAGTACACGACAGTTACCAG |
| | AGGAATATTTGTGTCCCCCA |
| *Slc7a14* | AAATGCCACGAACTCCCCAA |
| | GGGATGCCGTGATGGCGTAA |
| | CTTCGCAGCATAGAAGCCAT |
| | TCTGGGAGTGAAAAACTCTG |
| *Slc7a15* | AGTGATGACTGCCGTCCCCC |
| | TCGCAGCTGCATAGCACACG |
| | TGCTATGCAGCTGCGAGAGA |
| | GCAGGTAACTGATATTGACC |
| *Slc7a2* | AGGACGTCACTATTCCGATG |
| | GAACGGAACAAGCATCTACG |
| | GTATCTATACACTTACGTCA |
| | CCGAGACAACATATTTGGCG |
| *Slc7a3* | ATCTATCTCACCAATGACGT |
| | GCCATTGAATACATCACCCG |
| | CACCCAGAGCCACTAAGTCG |
| | AGTGCCGTTGGATTCACTCA |
| *Slc7a4* | AGGTATGCTGAGCCAGTACG |
| | GCACAGCCTAGACCCTGACT |
| | AATAACATCAAATCCCACGA |
| | GCCATGGCGTAGACAATGCG |
| *Slc7a5* | GCCCTCCTCGCAGTACATCG |
| | ACCCCTACTTACGCACGCAG |
| | AGCGGCCTCTTCGCCTACGG |
| | GTAGCAGAGTGCGCCCACGA |
| *Slc7a6* | AACTGAACATGCCGAATGTC |
| | TGCATTTAGGCCCCCGAAGC |
| | ACGCCTTTAAAGGTTCCTCG |
| | CAGCCACAGCGTCACTCTTA |
| *Slc7a7* | AAGAGATCAGGAACCCCGAG |
| | CAGCGCCAACACCTTAGCAT |
| | GGCCCCGGATTTCTTAATGG |
| | GAGACACACGCCATTAAGCA |
| *Slc7a8* | TGGAGCAGTTGACCCATGTG |
| | CCTGAAGAAAGAGATCGGAT |
| | TATGTGAAGGACATCTTCGG |
| | GACTCCACCAAACGTGGACA |
| *Slc7a9* | GAGCACTTACCAACCATCGT |
| | CAAAGGCCTCCATCAGATAG |
| | GGACTGCAAGAGCTCCGTTG |

| | |
|---|---|
| | GCTGGCCAACACAGAATCCG |
| *Slc8a1* | CTGATTATGAATTCACGGAA |
| | AATCGCACTCTGTGTTTACG |
| | ACTCACGTGAGCGAGAGCAT |
| | CTGCAATAATAGAAACTCCA |
| *Slc8a2* | GTGGACTACCGTACCGAGGA |
| | TTCCACTGCGAGGGTCGGTG |
| | ATCCTGGACGACGACCACGC |
| | CGGCAATGATAGACACACCC |
| *Slc8a3* | AGAACAATGAGTCCTGTTCG |
| | TACCGCACAGATAAACACCG |
| | GTACGTGGACTACAAAACAG |
| | CTGGGACCATCTACCATCGT |
| *Slc8b1* | TGACCACGTAGAACACGTAG |
| | CCACACACGGCTCGGCACTG |
| | TGGACCCCGACAAGGACGAT |
| | CCACACTCACAACTTGGCCG |
| *Slc9a1* | ACGGCCAACAGGTCGACCAG |
| | GGGGCGCATCACTACTCCTG |
| | CGCCCACAAACACCCCACCG |
| | GAAGGTCCAGTTCCACTGGT |
| *Slc9a2* | GGTCAGCACCTACCCGACAG |
| | TCGAAAGGGATTTGCACGTG |
| | AGTTCATCATTGCCTACGGA |
| | ACGCCATGATGCCTGAAAGG |
| *Slc9a3* | CTTCATAGTGTAGCGCACAG |
| | CATTGCGCTCTGGATCCTCG |
| | ACTTACCCATCAAGCCACTG |
| | CTTGGTGAAGCGAGTCACCA |
| *Slc9a4* | AGTGAAACGTGTGATAAATG |
| | TTAGTATGTTGTATAAGACC |
| | GACCGGAGGCGATTTGTGGT |
| | GAGATGCGAGCAGTATCCAG |
| *Slc9a5* | TGCCATCCTCACCTACGCCG |
| | CTCGATGATGCGGACCCGCT |
| | AATACTGACCGACGCCCTTT |
| | CCGCATCATCGAGCCGTTGC |
| *Slc9a6* | GGAAAGTGTCCTCAATGACG |
| | GAATGCCGTACCGAAGCACG |
| | TTTAACCCAACATTTGTCGT |
| | CATATGCTAAGATAGACCCA |
| | TACTTACGAGGATAGTACAA |

| | |
|---|---|
| *Slc9a7* | CATGAAGGAAACGCACCCGG |
| | CAGAAATCTTATGTATGGAG |
| | CCCTGGCAAGATCAACAACG |
| *Slc9a8* | CTTATTCACCTAGGACGAGG |
| | CGCTCGACTGCTCCTCCTGC |
| | CCACCTATTATCTTTGAGTC |
| | TATCCTGACTCAAAGATAAT |
| *Slc9a9* | GCGCCGACTGATATTGATAG |
| | ATGTTGACTAGTAGAGTTGA |
| | CACTTACCCTATGACCACAC |
| | CCTCCCATCATATTTCATGC |
| *Slc9b1* | TATACTTACCAATGAGAGGT |
| | AAGAAGGGAAACACAAACGA |
| | CCCAATAAGAACGTCCCGCA |
| | TCTAGGAGAGACTTACGCAG |
| *Slc9b2* | ATACCTTTGAATCTAGACCA |
| | CTGGCAAGAGTGATAACGAA |
| | CCAACCTTACTCATGGCCGC |
| | CAGCGGTGAAAAAAAGCCAC |
| *Slc9c1* | ATGTGCTCATTAATACCACA |
| | CCAACGACAGTACAGGAATG |
| | GACCAAGTCGAGACAAGATG |
| | CTTACCAAGGTCTCTTATGG |
| NTC | AAAAAGTCCGCGATTACGTC |
| | AAAACGGCTCGATCGGTGAT |
| | AAAACGTAATTATACCGAGC |
| | AAAATTGCACCTTCCCGGCC |
| | AAACCCCCGCGCGGAGCGTC |
| | AAACCTAGCGTAGATTCGGC |
| | AAACGAGGCTGTTCGTACAC |
| | AAACTCATACGTAGCGAATC |
| | AAACTCCCGTGTCAACCGAT |
| | AAAGACGTGCATTCAGCGAG |
| | AACATGTTAAGTCGCGTTAT |
| | AACCAGCATTTGACCGCGCT |
| | AACCCCGGCTGTCATCGCCG |
| | AACCCGCCGGAACAATCAGC |
| | AACCGGCTGCGCGTTTGCAA |
| | AACCGTACTGCGAGGAGCAT |
| | AACCTCGTCTCATGTACGAA |
| | AACGCCCCGGATTTCGTTGA |
| | AACGGCTGCGCCCGCGGCAA |

| | |
|---|---|
| | AACGGGCGCAATACCCTTTT |
