## Supplemental Table 3 for "Tissue-Specific Dependence of Th1 Cells on the Amino Acid Transporter SLC38A1 in Inflammation"

**Supplemental Table S5: Primers for CRISPR gRNA library prep**

| | | |
|---|---|---|
| | Forward | TAACTTGAAAGTATTTCGATTTCTTGGCTTTATATATCTTGTGGAAAGGACGAAACACCG |
| Array primers | Reverse | ACTTTTTCAAGTTGATAACGGACTAGCCTTATTTTAACTTGCTATTTCTAGCTCTAAAAC |
| | Forward | AATGGACTATCATATGCTTACCGTAACTTGAAAGTATTTCG |
| Adapter Primers | Reverse | ACGGCATCGCAGCTTGGATACA |
| | Forward | AATGATACGGCGACCACCGAGATCTACACTCTTTCCCTACACGACGCTCTTCCGATCTNNNNNNNNNTCTTGTGGAAAGGACGAAACACCG |
| Sequencing primers | Reverse | CAAGCAGAAGACGGCATACGAGATXXXXXXXXGTGACTGGAGTTCAGACGTGTGCTCTTCCGATCTACGGCATCGCAGCTTGGATACA |
