## Supplemental Figures for "Tissue-Specific Dependence of Th1 Cells on the Amino Acid Transporter SLC38A1 in Inflammation"

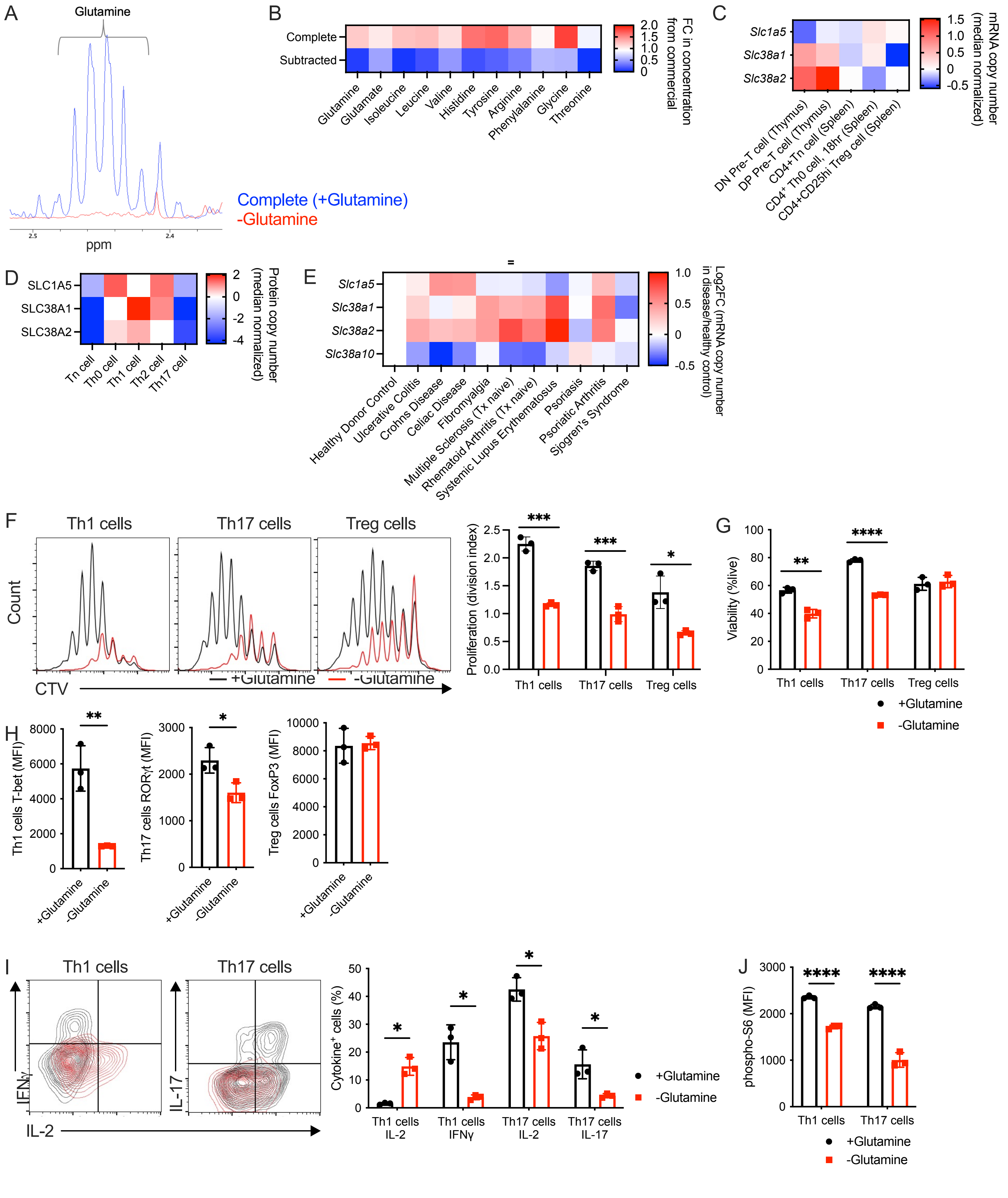


SUPPLEMENTAL FIGURE 1

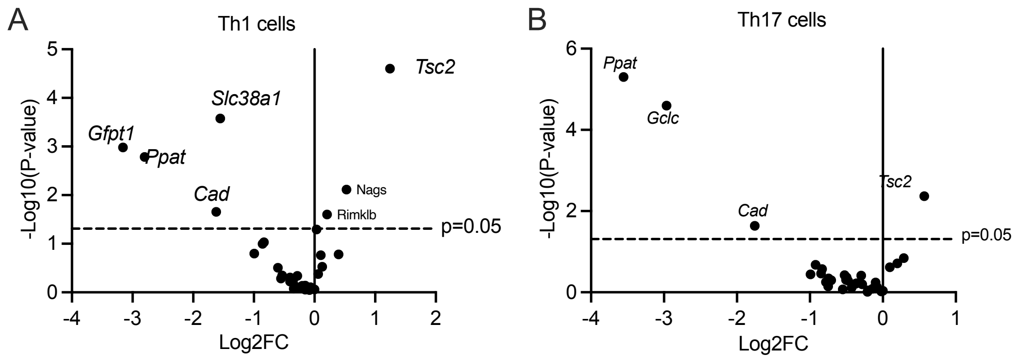


SUPPLEMENTAL FIGURE 2

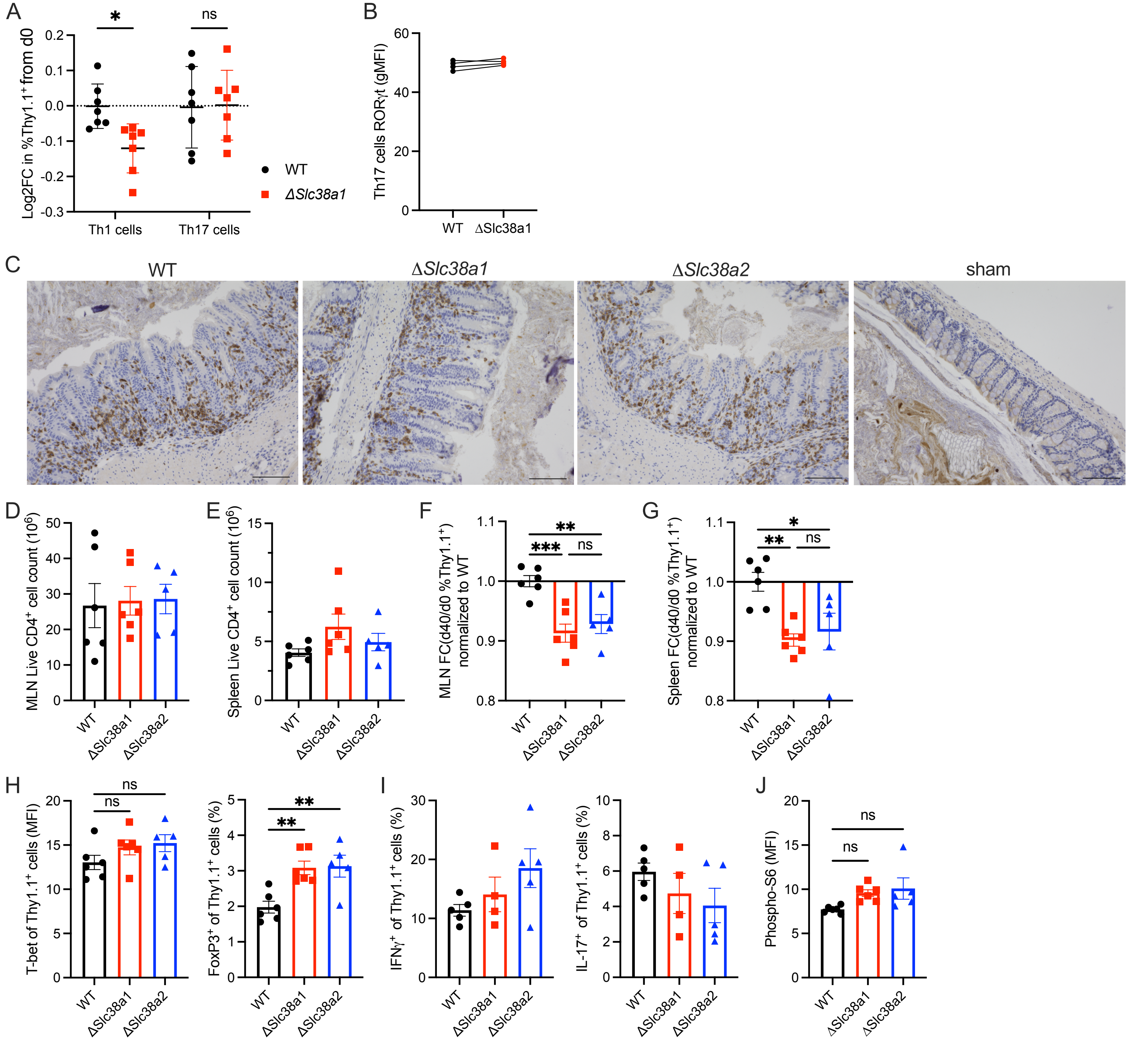


SUPPLEMENTAL FIGURE 3

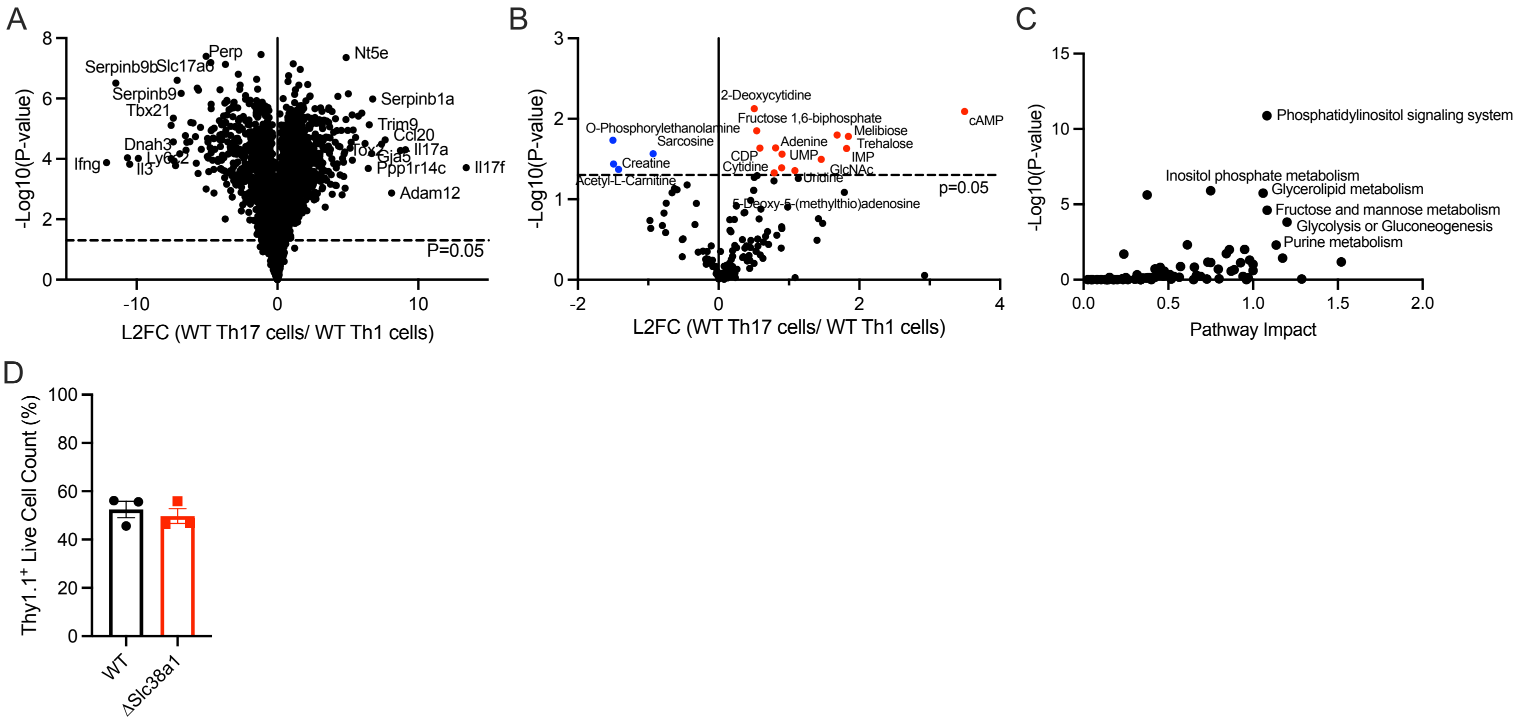


SUPPLEMENTAL FIGURE 4

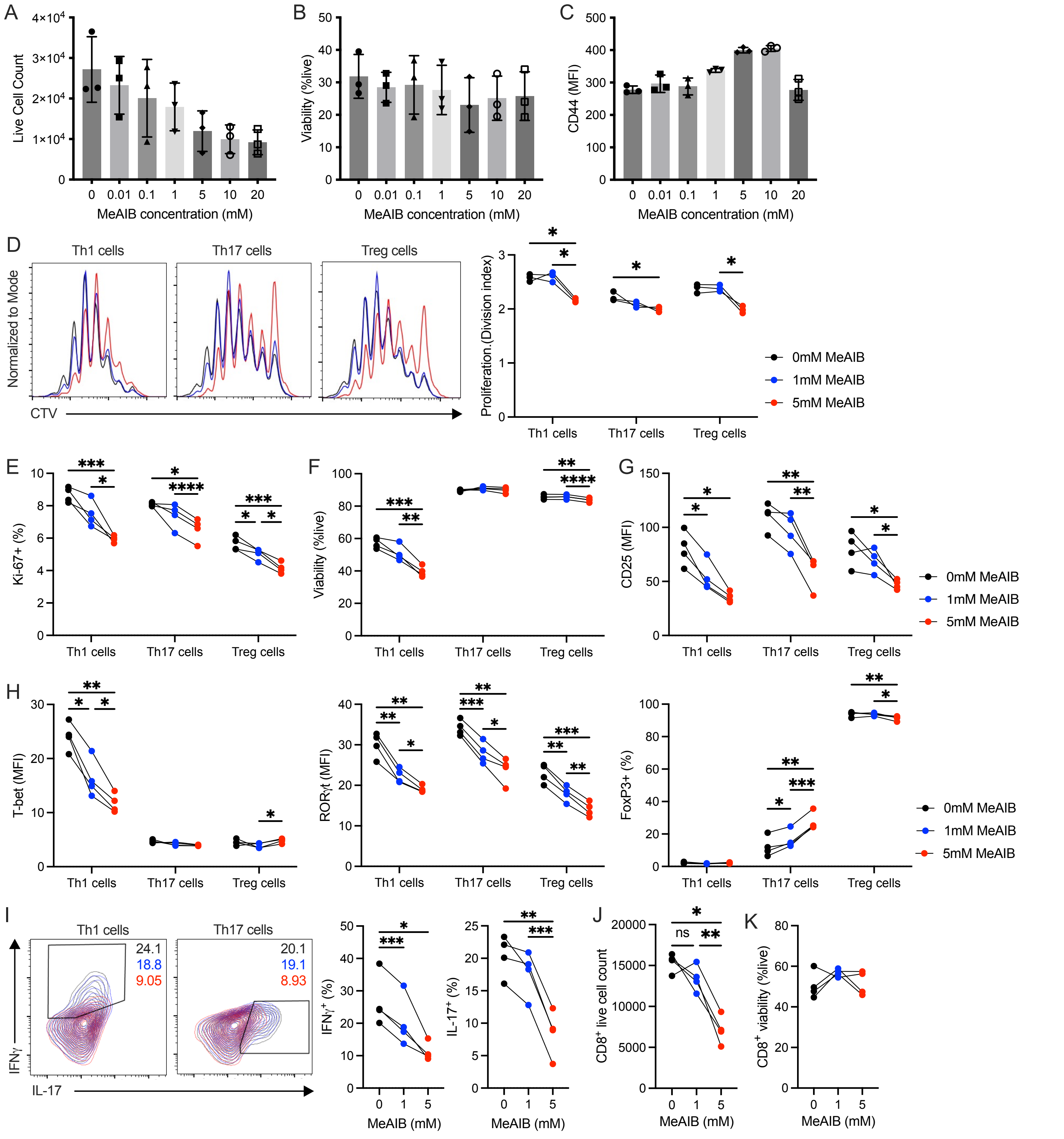


SUPPLEMENTAL FIGURE 5

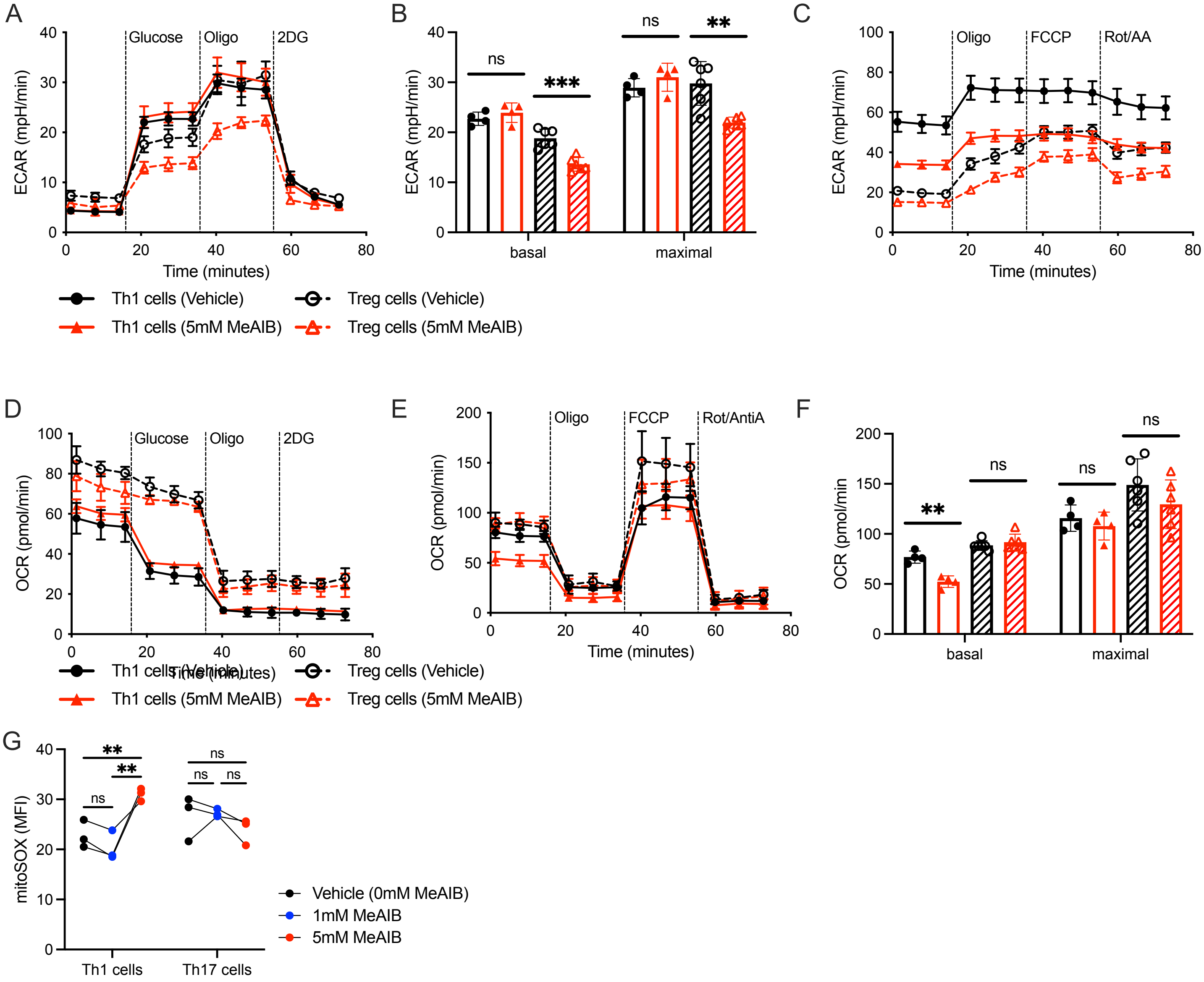


SUPPLEMENTAL FIGURE 6

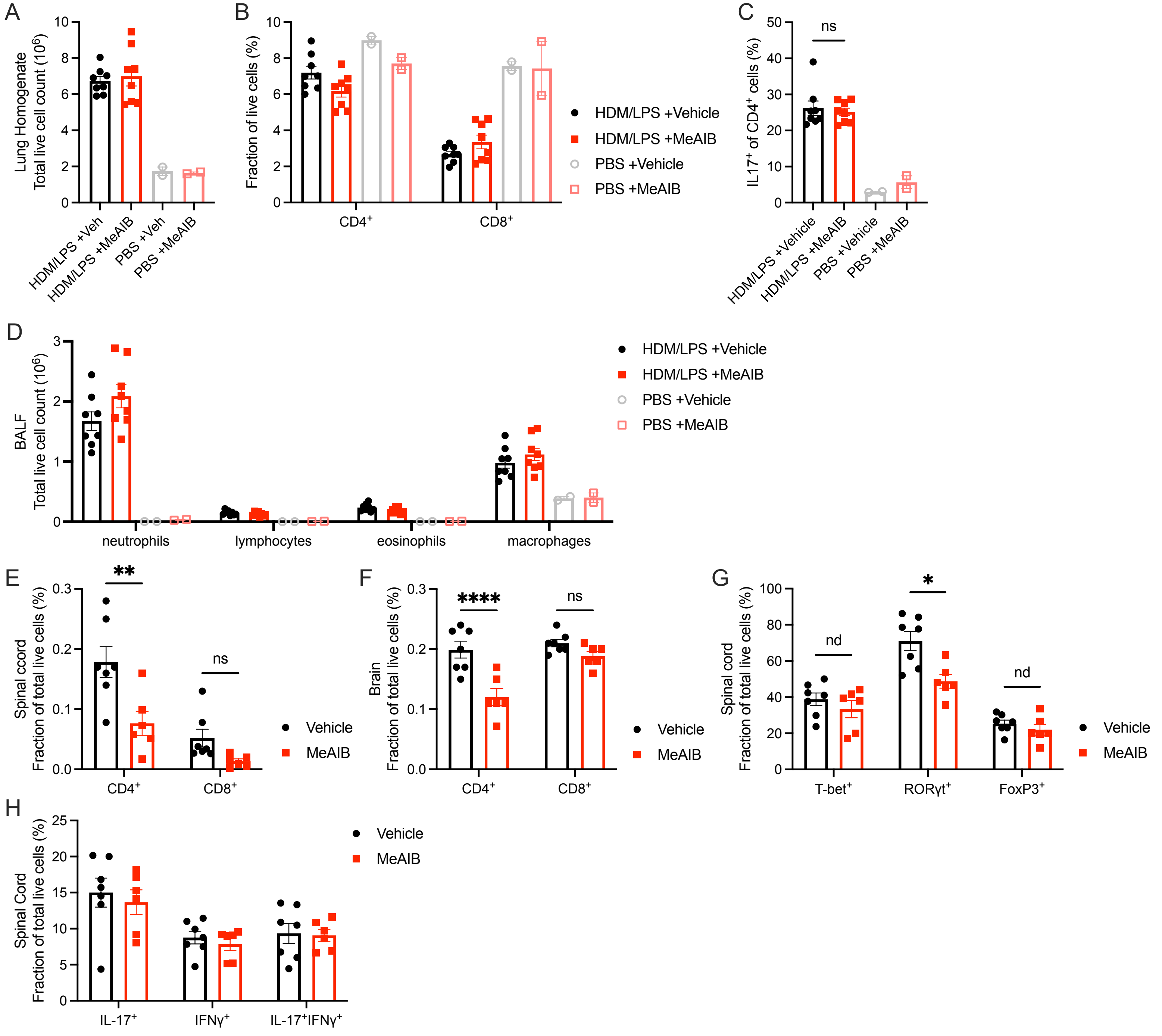


SUPPLEMENTAL FIGURE 7
