## Supplemental Figure Legends for "Tissue-Specific Dependence of Th1 Cells on the Amino Acid Transporter SLC38A1 in Inflammation"

### SUPPLEMENTAL MATERIALS

**Supplemental Figure 1: CD4$^+$ T cells are dependent on uptake of select essential and non-essential AAs.** (A) Representative NMR spectrum showing peaks assigned for glutamine in complete and -glutamine media. (B) Relative concentrations of the indicated AAs in complete media and AA subtracted media as detected by NMR. (C) Relative mRNA expression of glutamine transporters *Slc1a5*, *Slc38a1*, and *Slc38a2* in murine CD4$^+$ T cell subsets (ImmGen Deep RNAseq dataset, GSE122597). (D) Relative protein expression of SLC1A5, SLC38A1, and SLC38A2 in CD4$^+$ T cell subsets (Immunological Proteome Resource, ImmPRes) [1]. (E) mRNA expression Slc1a5, Slc38a1, Slc38a2, and Slc38a10 in whole blood of patients with indicated inflammatory and autoimmune diseases normalized to that in healthy controls. Data referenced from published RNAseq dataset [2] (GSE92472; mean, n=8 donors for healthy control and n=3-6 for patients with disease). Effect of glutamine deficiency on (F) proliferation as measured by CTV dilution and quantified by division index , (G) viability, lineage maintenance as measured by expression of (H) T-bet in Th1 cells, RORγt in Th17 cells, and FoxP3 in Treg cells, (I) effector functions as measured by cytokine expression, and (J) mTORC1 activity as measured by phosphor-S6 expression (mean±SD, two-way ANOVA for (F), (G), and (J), paired t-tests for (H), multiple paired t-tests for (I), n=3 biological replicates, flow cytometry plots show representative replicate).

**Supplemental Figure 2: *In vivo* CRISPR screening in Th1 cell- and Th17 cell-driven EAE show differential dependence on glutamine metabolism associated genes.** Volcano plot showing change in gRNA abundance from *in vivo* CRISPR screens in (A) Th1-cell driven and (B) Th17-cell driven EAE models (statistical analysis performed by MAGeCK, n=2 technical replicates). Comparison of the two datasets shown in Figure 2C.

**Supplemental Figure 3: Regulation of AA transporters and SLC38A1 dependence in CD4⁺ T cells.**

(A) Change in the frequency of gRNA transduced (Thy1.1+) cells in WT (NTC_1) and ∆Slc38a1 (SLC38A1_1) Th1 and Th17 cells on day 5 post transduction compared to day 1 (mean±SD, repeated measures two-way ANOVA with Šídák's multiple comparisons test, n=7 biological replicates).

(B) Expression of lineage-characterizing transcription factor RORγt in WT and ΔSlc38a1 Th17 cells (paired t-test, n=4 biological replicates across 2 independent experiments).

(C) Immunohistochemistry with anti-CD3 staining of colons collected from mice in WT, ΔSlc38a1, ∆Slc38a2, and sham (PBS control) groups showing T-cell infiltration of the colonic mucosa/submucosa from the IBD study (sham representative of 2 biological replicates, WT and ΔSlc38a1 representative of 6 biological replicates).

(D-E) Total CD4+ T cell count in the (D) MLNs and (E) spleens of mice from the IBD study (mean±SEM, Mann-Whitney test, n=6 biological replicates).

(F-G) Change in the fraction of gRNA-transduced Thy1.1+ cells in the transferred population on day 0 to the population recovered from the (F) MLN and (G) spleen of IBD mice at end of study on day 40 (mean±SEM, one-sample t test, n=6 biological replicates for WT, n=6 for ∆Slc38a1, n=5 for ∆Slc38a2).

(H-J) Expression of lineage-characterizing transcription factors (H) T-bet and FoxP3, (I) effector cytokines IFNγ and IL-17, and (J) mTORC1 target phosphor-S6 within the WT, ∆Slc38a1, and ∆Slc38a2 Thy1.1+ population recovered from the MLN of IBD mice at end of study (mean±SEM, one-way ANOVA with Tukey's multiple comparisons test, n=4-6 biological replicates).

**Supplemental Figure 4: Comparing SLC and metabolic pathway dependencies in WT Th1 and Th17 cells.**

(A) Volcano plot of RNAseq data showing fold change in mRNA expression between WT Th1 and Th17 cells measured by RNAseq (data analyzed with DESeq2 [3], n=3 biological replicates).
(B) Volcano plot of mass spectrometry data showing Log2 fold change in metabolite ion counts in WT Th1 and Th17 cells (median-normalized, n=3 biological replicates).
(C) Joint pathway analysis combining significantly changed genes from RNAseq and compounds from mass spectrometry performed on the same samples, identifying differential pathway dependencies between WT Th1 and Th17 cells (data analyzed with MetaboAnalyst v5.0, n=3 biological replicates).
(D) Fraction of the WT and ∆Slc38a1 Th1 cells that are Thy1.1+ after column enrichment representing successfully transduced population with consequent CRISPR/Cas9-mediated SLC38A1 loss (mean±SEM, unpaired t-test, n=4 biological replicates with 2-4 technical replicates per condition).

**Supplemental Figure 5: Treatment with SLC38A1/2 inhibitor MeAIB have differential effects on $CD4^+$ T cell subsets, and also reduces $CD8^+$ proliferation.**
(A-C) Effect of a range of MeAIB concentrations on (A) live cell count, (B) viability, and (C) activation marker CD44 expression on activated primary CD4+ T cells (mean±SD, repeated measures one-way ANOVA corrected with Tukey's multiple comparisons testing, n=3 biological replicates).
(D-E) Proliferation measured by (D) CTV dilution quantified as division index and (E) frequency of Ki-67+ cells in primary CD4+ T cells activated with anti-CD3 and anti-CD28 antibodies and cultured with Th1, Th17, or Treg cell polarizing cytokines and 0, 1, or 5mM MeAIB for 4 days (mean±SD, two-way ANOVA with Tukey's multiple comparisons test, n=3 biological replicates for CTV dilution and n=4 for Ki-67, flow cytometry plots show representative replicate).

(F-G) Cell (F) viability, (G) CD25 activation marker, and (H) lineage-characterizing transcription factors in cells prepared as above (mean±SD, two-way ANOVA with Tukey's multiple comparisons test, n=4 biological replicates).

(I) Cytokine expression in cells cultured with Th1 or Th17 cell polarizing cytokines and 0, 1, or 5mM MeAIB for 4 days post activation (mean±SD, one-way ANOVA, n=4 biological replicates, flow cytometry plots show representative replicate).

(J-K) Live cell (J) count and (K) fraction in primary CD8+ T cells activated with anti-CD3 and anti-CD28 antibodies and cultured with IL-2 and 0, 1, or 5mM MeAIB for 4 days (mean±SD, one-way ANOVA, n=4 biological replicates).

**Supplemental Figure 6: SLC38A1/2 inhibition by MeAIB differentially affects glycolytic and mitochondrial flux in Th1 and Th17 cells.**

(A-C) Glycolytic measurment of (A) change in ECAR reflecting glycolytic rate in response to sequential addition of glucose, oligomycin (Oligo), and 2-deoxyglucose (2-DG), (B) basal and maximal ECAR, and (C) change in ECAR in response to sequential addition of (Oligo), carbonylcyanide-p-trifluoromethoxyphenylhydrazone (FCCP), and rotenone/antimycin A (Rot/AA) in Th1 and Treg cells treated with either vehicle or 5mM MeAIB (mean±SEM, ordinary two-way ANOVA with Tukey's multiple comparisons test, n=4 biological replicates for Th1 cells and n=6 for Treg cells with 6-8 technical replicates per condition).

(D-F) Mitochondrial oxygen consumption measurement of (D) change in OCR in response to sequential addition of glucose, oligomycin (Oligo), and 2-deoxyglucose (2-DG), (E) OCR in response to in response to sequential addition of (Oligo), carbonylcyanide-p-trifluoromethoxyphenylhydrazone (FCCP), and rotenone/antimycin A (Rot/AA), (F) basal and maximal OCR in Th1 and Treg cells treated with either vehicle or 5mM MeAIB (mean±SEM, ordinary two-way ANOVA with Tukey's multiple comparisons test, n=4 biological replicates for Th1 cells and n=6 for Treg cells with 6-8 technical replicates per condition).

(G) Mitochondrial superoxide levels measured by mitoSOX staining in (A) Th1, Th17, and Treg cells treated with 0, 1, or 5mM MeAIB for 4 days (mean±SD, one-way ANOVA, n=3 biological replicates).

**Supplemental Figure 7: SLC38A1/2 inhibition in vivo has no effect on Th17-cell driven allergic airway disease model as predicted by in vivo CRISPR screen.**

(A) Total live cell count measured from the lung homogenate collected at the end of study from mice with allergic airway disease and treated with either vehicle or MeAIB. Allergic airway disease induced by sensitization and challenge with intranasal administration of HDM/LPS over 2 weeks, treated daily with either vehicle or MeAIB from 7 days prior to initial sensitization until end of study (mean±SEM, unpaired t-test, n=8 biological replicates for HDM/LPS, n=2 each for PBS control).

(B-C) Frequencies of (B) CD4+ and CD8+ T cell among total live cell population and (C) IL-17+ cells among CD4+ T cells in the lung homogenate from (A) (mean±SEM, multiple unpaired t tests, n=8 biological replicates for HDM/LPS, n=2 each for PBS controls).

(D) Cell count of neutrophils, lymphocytes, eosinophils and macrophages found in the BALF of mice with allergic airway disease and control mice treated with either vehicle or MeAIB (mean±SEM, multiple unpaired t tests corrected for multiple comparisons by FDR, Q=5%, n=8 biological replicates for HDM/LPS, n=2 each for PBS controls).

(E-F) CD4+ and CD8+ T cell frequency in the (E) spinal cords and (F) brains collected at the end of study from mice in Figure 7A with EAE treated daily with either vehicle or MeAIB (mean±SEM, multiple Mann-Whitney tests corrected for multiple comparisons by FDR, Q=5%, n=7 biological replicates for vehicle and n=6 for MeAIB).

(G-H) Frequency of lineage-characterizing (G) transcription factors T-bet+, RORγt+, and FoxP3+ and (H) effector cytokines IL-17+, IFNγ+, and IL-17+IFNy+ cells among total CD4+ T cell population recovered from the spinal cord of EAE mice shown in (mean±SEM, multiple unpaired t-tests corrected for multiple comparisons by FDR, Q=5%, n=7 biological replicates or vehicle and 8 for MeAIB).
