## Supplementary material for "Tissue-Specific Dependence of Th1 Cells on the Amino Acid Transporter SLC38A1 in Inflammation": Methods

### KEY RESOURCES TABLE

| REAGENT or RESOURCE | SOURCE | IDENTIFIER |
|---|---|---|
| **Antibodies** | | |
| Anti-IFNγ | Thermo Fisher Scientific | Cat# 16-7311-38, RRID:AB_2637490 |
| Anti-IL-4 | Thermo Fisher Scientific | Cat# 16-7041-95, RRID:AB_2573101 |
| Mouse Anti-CD3e | Thermo Fisher Scientific | Cat# 16-0031-86, RRID:AB_468849 |
| Mouse Anti-CD28 | Thermo Fisher Scientific | Cat# 16-0281-86, RRID:AB_468923 |
| Anti-IFNγ APC | BD Biosciences | Cat# 554413, RRID:AB_398551 |
| Anti-IL-17A PE | Thermo Fisher Scientific | Cat# 12-7177-81, RRID:AB_763582 |
| Anti-CD25 e450 | Thermo Fisher Scientific | Cat# 48-0251-82, RRID:AB_10671550 |
| Anti-CD44 PE | Thermo Fisher Scientific | Cat# 12-0441-82, RRID:AB_465664 |
| Anti-CD62L APC | Thermo Fisher Scientific | Cat# 17-0621-82, RRID:AB_469410 |
| Anti-CD4 eFluor 450 | Thermo Fisher Scientific | Cat# 48-0041-82, RRID:AB_10718983 |
| Anti-CD4 PE | BioLegend | Cat# 100512, RRID:AB_312715 |
| Anti-CD8a PE | Thermo Fisher Scientific | Cat# 12-0081-82, RRID:AB_465530 |
| Anti-FoxP3 APC | Thermo Fisher Scientific | Cat# 17-5773-82, RRID:AB_469457 |
| Anti-T-bet PE | Thermo Fisher Scientific | Cat# 12-5825-82, RRID:AB_925761 |
| Anti-RORγt PE | Thermo Fisher Scientific | Cat# 12-6988-82, RRID:AB_1834470 |
| Anti-Ki-67 eFluor 450 | Thermo Fisher Scientific | Cat# 48-5698-82, RRID:AB_11149124 |
| Anti-SLC38A1 | Cell Signaling Technology | Cat# 36057, RRID:AB_2799092 |
| Anti-ß-actin | Cell Signaling Technology | Cat# 4970 RRID:AB_2223172 |
| IRDye 800CW Secondary | LI-COR Biosciences | Cat# 926-32211, RRID:AB_621843 |
| IRDye 680LT Secondary | LI-COR Biosciences | Cat# 926-68020, RRID:AB_10706161 |
| **Bacterial strains** | | |
| ElectroMAX DH10B Cells | Thermo Fisher Scientific | Cat#: 18290015 |
| **Chemicals, peptides, and recombinant proteins** | | |
| Recombinant murine IL-12p70 | Thermo Fisher Scientific | Cat# 14-8121-62 |

| | | |
|---|---|---|
| Recombinant murine IL-6 | Miltenyi Biotec | Cat# 130-096-683 |
| Recombinant murine IL-23 | Miltenyi Biotec | Cat# 130-096-676 |
| Recombinant murine IL-1ß | Miltenyi Biotec | Cat# 130-101-681 |
| Recombinant human IL-2 | NCI | Cat# Ro 23-6019 |
| Recombinant human TGFß1 | Peprotech | Cat# 100-21 |
| MeAIB | Sigma-Aldrich | Cat# M2283 |
| GolgiPlug | BD Biosciences | Cat# 555029 |
| Retronectin | Takara Bio | Cat# T100A |
| $OVA_{323-339}$ peptide (chicken, Japanese quail) | Sigma-Aldrich | Cat# O1641 |
| $MOG_{35-55}$ peptide | GenScript | Cat# RP10245 |
| EndoFit Ovalbumin | InVivoGen | Cat# vac-pova |
| House dust mite | Greer Laboratories | Cat# XPB70D3A25 |
| LPS | Sigma-Aldrich | Cat# L5293 |
| **Critical commercial assays** | | |
| EasySep Mouse CD4 T Cell Isolation Kit | STEMCELL Technologies | Cat# 19852 |
| EasySep Mouse CD90.1 Positive Selection Kit | STEMCELL Technologies | Cat# 18958 |
| Fixation/Permeabilization Solution Kit | BD Biosciences | Cat# 554714 |
| Foxp3/Transcription Factor Staining Buffer Set | Thermo Fisher Scientific | Cat# 00-5523-00 |
| KAPA Mouse Gentoyping Kits | Roche Diagnostics | Cat# 07961804001 |
| Seahorse XFe96 FluxPaks | Agilent Technologies | Cat# 102601-100 |
| Herculase II Fusion DNA Polymerase | Agilent | Cat# 600675 |

| | | |
|---|---|---|
| Gibson Assembly Master Mix | NEB | Cat# E2611 |
| Polyplus jetPRIME DNA and siRNA transfection reagent | VWR | Cat# 89129-922 |
| EAE induction kit | Hooke Labs | Cat# EK-2110 |
| Epredia Richard-Allan Scientific Three-Step Stain Kit | Thermo Fisher Scientific | Cat# 22-050-272 |
| **Deposited data** | | |
| SLC transporter in vitro screen in Th0 cells, replicates 1&2 | This paper | Functional ImmunoGenomics resource (FIGS; https://figs.app.vumc.org/) |
| Glutamate metabolism in vitro screen in Th0 cells | This paper | FIGS |
| Glutamate metabolism in *vivo* EAE screen in Th1/17 cells | This paper | FIGS |
| Glutamate metabolism in *vivo* lung inflammation screen in Th0 cells | This paper | FIGS |
| Δ*Slc38a1* Th1/Th17 cells bulk RNAseq | This paper | GEO# GSE190131 |
| Δ*Slc38a1* Th1/Th17 cells Metabolomics | This paper | |
| **Experimental models: Cell lines** | | |
| Plat-E retroviral packaging cell line | Cell Biolabs | Cat# RV-101, RRID:CVCL_B488 |
| **Experimental models: Organisms/strains** | | |
| Mouse: C57BL/6J | Jackson Laboratory | Strain#: 000664 RRID:IMSR_JAX:000664 |

| | | |
|---|---|---|
| Mouse: Rag1-/- (B6.129S7-Rag1tm1Mom/J) | Jackson Laboratory | Strain#: 002216<br>RRID:IMSR_JAX:002216 |
| Mouse: OT-II (B6.Cg-Tg(TcraTcrb)425Cbn/J) | Jackson Laboratory | Strain#: 004194<br>RRID:IMSR_JAX:004194 |
| Mouse: 2D2 (C57BL/6-Tg(Tcra2D2,Tcrb2D2)1Kuch/J) | Jackson Laboratory | Strain#: 006912<br>RRIDIMSR_JAX:006912 |
| Mouse: Cas9 (B6J.129(Cg)-Gt(ROSA)26Sortm1.1(CAG-cas9*,-EGFP)Fezh/J) | Jackson Laboratory | Strain#: 026179<br>RRID:IMSR_JAX:026179 |
| **Oligonucleotides** | | |
| NTC gRNA#1 - AAAAAGTCCGCGATTACGTC | Mouse Brie CRISPR knockout pooled library, Control sgRNAs (Doench *et al.,* 2016) [26] | Addgene #73633;<br>RRID:Addgene_73633 |
| NTC gRNA#2 - ATTGTTCGACCGTCTACGGG | Mouse CRISPR Knockout Pooled Library, GeCKO v2, Mouse library A gRNA sequences (Sanjana *et al*., 2014) [27] | Addgene #1000000052<br>(NonTargetingControlGuideForMouse_0007, MGLibA_66412) |
| NTC gRNA#3 - ACCCATCGGGTGCGATATGG | GeCKO v2 | Addgene #1000000052<br>(NonTargetingControlGuideForMouse_0008, MGLibA_66413) |
| *Slc38a1* gRNA#1 - TGCATGGTGTATGAGAAGCT | Brie library target genes | Addgene #73633;<br>RRID:Addgene_73633 |

| | | |
|---|---|---|
| *Slc38a1* gRNA#2 - AGATTGGCAGGACGGACGG G | Brie library target genes | Addgene #73633; RRID:Addgene_73633 |
| *Slc38a2* gRNA#1 - CTCAAGACTGCCAACGAAGG | Brie library target genes | Addgene #73633; RRID:Addgene_73633 |
| SLC gRNA library | This paper | See **Table S1** |
| Glutamine metabolism gRNA library | This paper | See **Table S2** |
| Primers for CRISPR gRNA library prep | Modified from Shalem *et al.*, 2014 [52] | See **Table S3** |
| **Recombinant DNA** | | |
| pMx-U6-gRNA scaffold-PGK-GFP plasmid | Toffalini *et al.*, 2009 [49] | N/A |
| MSCV-IRES-Thy1.1 DEST plasmid | Addgene | Addgene #17442; RRID:Addgene_17442 |
| MSCV-U6-gRNA scaffold-IRES-Thy1.1 plasmid | This paper | N/A |
| **Software and algorithms** | | |
| Prism v10 | GraphPad Software | https://www.graphpad.com; RRID:SCR_002798 |
| FlowJo v10 | FlowJo | https://www.flowjo.com; RRID:SCRJD08520 |
| MAGeCK v0.5.0.3 | Li *et al.*, 2014 [46] | http://liulab.dfci.harvard.edu/Mageck |
| TopSpin v3.6 | Bruker | https://www.bruker.com/en/products-and-solutions/magnetic-resonance/nmr-software/topspin.html; RRID:SCR_014227 |

| | | |
|---|---|---|
| MetaboAnalyst v5.0 | Pang *et al*., 2021 [48] | https://www.metaboanalyst.ca; RRID:SCR_015539 |
| Agilent Wave software v2.6 | Agilent | https://www.agilent.com/en/products/cell-analysis/software-download-for-wave-desktop; RRID:SCR_014526 |
| DESeq2 v1.24.0 | Love et al., 2014 [47] | http://www.bioconductor.org/packages/release/bioc/html/DESeq2.html |
| Cutadapt v2.10 | | N/A |
| STAR v2.7.3a | | N/A |
| featureCounts v2.0.0 | | N/A |

### METHODS

#### Resource Availability

*Lead Contact:* Further information and requests for resources and reagents should be directed to and will be fulfilled upon completion of appropriate MTA by the Lead Contact, Dr. Jeffrey Rathmell.

*Materials Availability:* Plasmids and mouse lines generated in this study will be made available upon completion of appropriate MTA upon request.

*Data and code availability:* RNAseq data have been deposited at GEO and are publicly available as of the date of publication (GSE190131). Microscopy data and original western blot images reported in this paper will be shared by the lead contact upon request. Any additional information required to reanalyze the data reported in this paper is available from the lead contact upon request. CRISPR screening data and detailed library information will be available: https://figs.app.vumc.org/figs/.

**Experimental Model Details**

_**Mice:**_ All experiments were performed at Vanderbilt University animal facility in accordance with Institutional Animal Care and Utilization Committee (IACUC)-approved protocols and conformed to all relevant regulatory standards. Mice were housed in pathogen-free facilities in ventilated cages with *ad libitum* food and water and at most 5 animals per cage. Eight- to sixteen-week-old male and female mice were used for all animal experiments. All mice were obtained from Jackson Laboratory and were treatment-naive until the start of study. 2D2 mice (C57BL/6-Tg(Tcra2D2,Tcrb2D2)1Kuch/J, JAX Strain#: 006912) were crossed to Cas9 mice (B6J.129(Cg)-Gt(ROSA)26Sortm1.1(CAG-cas9*,-EGFP)Fezh/J, JAX Strain#: 006912) to generate 2D2 Cas9 double-transgenic strain, and OT-II mice (B6.Cg-Tg(TcraTcrb)425Cbn/J, JAX Strain#: 004194) were crossed to Cas9 mice to generate OT-II Cas9 double-transgenic strain. Animals were genotyped for transgenic TCR and Cas9 allele.

Cell Lines: Plat-E retroviral packaging cell line was maintained at 37°C with 5% $CO_2$ in DMEM media supplemented with 10% FBS, 100U/mL penicillin/streptomycin, 1μg/mL puromycin, and 10μg/mL blasticidin to maintain expression of viral packaging genes.

**Method Details**

*In vitro mouse CD4$^+$ T cell activation and differentiation:* Primary murine CD4$^+$ T cells were isolated from the spleens and lymph nodes of mice using a CD4 negative isolation kit according to the manufacturer's instructions. The cells were cultured at 37°C with 5% $CO_2$ in RPMI-1640 media supplemented with 10mM HEPES, 50μM 2-mercaptoethanol, 100U/mL penicillin/streptomycin, and 2mM AA, unless otherwise stated. AA subtracted media were produced by adding back all AAs except the subtracted one at concentrations specified in the RPMI-1640 media formulation to modified RPMI-1640 media lacking all AAs (200mg/L Larginine, 56.82mg/L L-asparagine•H2O, 20mg/L L-aspartic acid, 65.2mg/L L-cystine•2HCl, 20mg/L L-glutamic acid, 300mg/L L-AA, 10mg/L glycine, 15mg/L L-histidine, 20mg/L hydroxyl-L-proline, 50mg/L L-isoleucine, 50mg/L L-leucine, 40mg/L L-lysine•HCl, 15mg/L L-methionine, 15mg/L L-phenylalanine, 20mg/L L-proline, 30mg/L L-serine, 20mg/L L-threonine, 5mg/L L-tryptophan, 28.83mg/L L-tyrosine•2Na•2$H_2O$, 20mg/L L-valine). The pH of the final media was adjusted to 7.2.

Primary $CD4^+$ T cells from wildtype and Cas9-transgenic mice were activated using plate-bound anti-CD3 (3μg/mL) and anti-CD28 (5μg/mL) antibodies at 1 million cells/well in a 24-well plate. OT-II/Cas9 double-transgenic T cells were activated with splenocytes irradiated at 30 Gy and $OVA_{323\text{-}339}$ peptide (10μg/mL), and 2D2/Cas9 double-transgenic T cells with $MOG_{35\text{-}55}$ peptide (10μg/mL). T cells were cultured for 4 days with subset-specific cytokines and blocking antibodies to promote differentiation - Th1 cells: IL-12p70 (10ng/mL), IL-2 (100U/mL), anti-IL-4 (10μg/mL), anti-IFNγ (1μg/mL); Th17 cells: IL-6 (50ng/mL), TGFβ (1ng/mL), IL-23 (20ng/mL), IL-1β (10ng/mL), anti-IL-4 (10μg/mL), anti-IFNγ (10μg/mL); Treg cells: TGFβ (1.5ng/mL), IL-2 (100U/mL), anti-IL-4 (10μg/mL), anti-IFNγ (10μg/mL). MeAIB (SLC38A1/2 inhibitor) was dosed at 1mM and 5mM for *in vitro* studies.

<u>*Flow Cytometry:*</u> Cells were first stained with viability dye and antibodies for pertinent surface markers for all experiments. For intracellular and transcription factor stains, cells were then fixed and permeabilized using appropriate kits. For cytokines, cells were stimulated with 1μg/mL 12-myristate 13-acetate (PMA), 750ng/mL ionomycin, and GolgiPlug for four hours before fixation. Unstimulated cells served as negative control. All dilutions and washes were performed in PBS with 10% FBS.

<u>*In vitro single gene knockout using CRISPR/Cas9:*</u> MSCV-U6-gRNA scaffold-IRES-Thy1.1 was cloned by excising the U6 promoter and gRNA scaffold from the pMx-U6-gRNA scaffold-PGK-GFP vector [49] and inserting into the MSCV-IRES-Thy1.1 DEST vector (Addgene #17442) [50]. NTC and targeted gRNA sequences were referenced from the Mouse CRISPR Knockout

Pooled Libraries Brie (Addgene #73633) [26] and GeCKO v2 (Addgene #1000000052) [27], and cloned into the new MSCV-U6-gRNA scaffold-IRES-Thy1.1 vector following the protocol made publically available by the Zhang Lab. The gRNA sequences used are as follows: NTC gRNA#1 AAAAAGTCCGCGATTACGTC (Brie), NTC gRNA#2 ATTGTTCGACCGTCTACGGG (GeCKO v2, MGLibA_66412), NTC gRNA#3 ACCCATCGGGTGCGATATGG (GeCKO v2, MGLibA_66413), *Slc38a1* gRNA#1 TGCATGGTGTATGAGAAGCT (Brie), *Slc38a1* gRNA#2 AGATTGGCAGGACGGACGGG (Brie), and *Slc38a2* gRNA#1 (CTCAAGACTGCCAACGAAGG, Brie). For experiments with single gRNA in multiple biological replicates, NTC gRNA#1, *Slc38a1* gRNA#1, and *Slc38a2* gRNA#1 were used to generate WT, Δ*Slc38a1*, and Δ*Slc38a2* cells, respectively.

Briefly, Plat-E retroviral packaging cell line was transfected using Polyplus jetPRIME DNA and siRNA transfection reagent with the gRNA sequence-cloned expression. The viral supernatants were collected, filtered, and spun onto retronectin-coated plates at 2000xg for 2 hours at 32°C. In parallel, Cas9-transgenic $CD4^+$ T cells were activated and cultured for 48 hours, transferred to the prepared virus-bound plates, and centrifuged at 2000xg for 15 minutes at 32°C. Transduced cells were identified by Thy1.1 positivity by flow cytometry, and enriched for using CD90.1 positive selection kit for downstream applications including RNAseq and metabolomics.

<u>*In vitro CRISPR screening:*</u> Custom SLC transporter (**Table S1)** and glutamine metabolism (**Table S2)** gRNA libraries were curated by referencing the Brie and GeCKO v2 libraries. Pooled plasmid libraries were prepared following published methods [51,52] and the *in vitro* and *in vivo* CRISPR screen using the lung-infammation model were performed in primary $CD4^+$ T cells as previously described [28]. Briefly, each gRNA library was synthesized as an oligonucleotide pool he following sequence:

GGAAAGGACGAAACACCGXXXXXXXXXXXXXXXXXXXXGTTTTAGAGCTAGAAATAGCAAG TTAAAATAAGGC, where Xs denote the variable gRNA sequence. The oligo pool was bulk cloned into the pMx-U6-gRNA scaffold-PGK-GFP vector by PCR using Array primers (**Table S3)** followed by Gibson Assembly. The resultant plasmid pool was amplified using ElectroMAX DH10B Cells, with greater than 50-fold coverage of the library, and packaged in retrovirus. Cas9-transgenic $CD4^+$ T cells were activated for 48 hours and transduced at a multiplicity of infection (MOI) of 0.4. Pre- and post-selection cell samples were collected as specified for each screen.

Next, genomic DNA was isolated from cells and gRNA sequences were amplified by two rounds of PCR using Adapter primers followed by barcoded sequencing primers (**Table S3**). Amplicons were sequenced to obtain 150bp paired-end reads on the Illumina NovaSeq 6000 platform. At least 1000-fold representation of the library was maintained throughout the assay. FASTQ files were analyzed using the Model-based Analysis of Genome-wide CRISPR/Cas9 Knockout (MAGeCK v0.5.0.3) method [46].

*In vivo CRISPR screening:* The Th1- and Th17-cell driven EAE models were developed by modifying previously published methods for EAE induction by adoptive transfer of differentiated 2D2-transgenic $CD4^+$ T cells [32]. First, $CD4^+$ T cells were isolated from the spleen and lymph nodes of 2D2/Cas9 double-transgenic mice and activated with splenocytes irradiated at 30 Gy and $MOG_{35-55}$ peptide (10µg/mL). Cells were cultured for 3 days with cytokines and blocking antibodies to promote either Th1 or Th17 cell differentiation. Three days post activation, cells were split in fresh media with IL-2 for Th1 cells and IL-23 for Th17 cells and cultured for an additional three days. Six days post activation, cells were re-stimulated with plate-bound anti-CD3 (3µg/mL) and anti-CD28 (5µg/mL) antibodies and transduced with the gRNA library as above. 24 hours post transduction, a sample of the cells was collected for the early timepoint and approximately 10 million live cells were adoptively transferred into separate *Rag1*$^{-/-}$ mice for each subset by tail vein injection. On day 0 and 1 post adoptive transfer, the recipient mice were i.p. injected with 150ng PTX. Mice were monitored daily for clinical symptoms of disease progression. Once mice began exhibiting signs of bilateral hind leg paralysis and/or severe ataxia, they were sacrificed for brain and spinal cord collection. The tissues were mechanically dissociated and digested with 300U/mL Collagenase IA and 50U/mL DNase I at 37 °C for 45 minutes, and then filtered through a 70μm filter to obtain a single-cell suspension. The cells were subsequently layered on a 18.6%/62.4% Percoll gradient and centrifuged at 2400rpm for 30 minutes at room temperature to partially purify the T cells for the late timepoint collection. Cells were processed, sequenced, and analyzed in the same manner as described for *in vitro* screening.

The *in vivo* CRISPR screen in the inflammatory lung disease model was performed following published methods [28]. OT-II/Cas9 double transgenic $CD4^+$ T cells were activated with irradiated splenocytes and $OVA_{323-339}$ peptide, transduced with the gRNA library, and adoptively transferred into *Rag1*$^{-/-}$ mice. Recipient mice were intranasally administered ovalbumin protein on days 1, 3, 5, and 7 post adoptive transfer, and sacrificed on day 8 for lung collection. Lungs were mechanically dissociated using a gentleMACS Dissociator (Miltenyi Biotec), digested with 300U/mL Collagenase IA and 50U/mL DNase I at 37°C for 45 minutes, and then filtered through a 70μm filter to obtain a single-cell suspension. $CD4^+$ T cells were isolated using a positive selection kit according to the manufacturer's instructions for the late timepoint collection.

*RNAseq analysis:* Bulk RNA was isolated using the RNeasy Mini Kit. mRNA enrichment and cDNA library preparation was performed using the stranded mRNA (polyA-selected)library preparation kit (NEB). cDNA was sequenced to obtain 150bp paired-end reads on the Illumina NovaSeq 6000 platform targeting an average of 50 million reads per sample. Demultiplexed FASTQ files were analyzed as follows. Adapters were first trimmed by Cutadapt (v2.10). Reads were then mapped to the mouse genome mm10 using STAR (v2.7.3a) and quantified by featureCounts (v2.0.0). DESeq2 (v.1.24.0) [47] was used to detect differential expression between the two groups.

*Extracellular flux analysis:* Extracellular flux analyses were performed with the Seahorse XFe96 Analyzer (Agilent). Assay media were prepared by supplementing Seahorse XF RPMI Medium pH 7.4 with 10 mM glucose, 1 mM sodium pyruvate, and 2 mM glutamine, unless otherwise stated. Cells were seeded on 96-well cell culture microplates coated with Cell-Tak (1mg/mL) at 150,000 live cells per well with technical replicates for each biological replicate. The Glycolytic Stress Test was performed The Mito Stress Test was performed with 1.5μM oligomycin A, 1.5μM FCCP, and 0.5μM rotenone/antimycin A final concentrations. Data were analyzed in Agilent Wave software v2.6.

*Immunoblotting:* Cells were lysed on ice for 30 minutes with base lysis buffer containing 1% IGEPAL CA-630, 200mM NaCl, 50mM Tris pH8.0, and supplemented with the protease inhibitors aprotinin (5ug/mL), leupeptin (5ug/mL), sodium fluoride (0.9mM), dithiothreitol (DTT, 1mM), sodium vanadate (1mM), and ß-glycerophosphate (20mM). Lysates were centrifuged for 15 minutes at 4˚C to recover supernatant and quantified for protein concentration using Protein Assay Dye Reagent Concentrate. 40ug of protein was loaded per well in Mini-PROTEAN Precast Polyacrylamide Gels (Bio-Rad) for electrophoresis using. Western blotting was performed using low fluorescence PVDF membrane (Bio-Rad). Transfer was accomplished using 1X Towbin Transfer Buffer Containing 20% methanol at 300mA for 1 hour. Blots were blocked for 1 hour using Intercept (TBS) Blocking Buffer (LI-COR Biosciences) and incubated overnight at 4˚C with primary antibody overnight at 4˚C (1:1000 for anti-ß-actin, 1:1000 for anti-SLC38A1). Blots were incubated for 1 hour at room temperature with IRDye Secondary antibodies and visualized by near infrared fluorescence via Li-COR Odyssey CLx imager.

*Mass Spectrometry:* Metabolites were extracted from cells by adding cold 80% methanol and incubating at -80°C for 10 minutes, followed by centrifugation at 10,000xg for 10 minutes at 4°C. Supernatants were collected and lyophilized by speedvac. The quantity of metabolite fraction analyzed was normalized to cell number. Liquid chromatography-based targeted tandem mass spectrometry (LC-MS/MS)-based metabolomics were performed and the data analyzed as previously described [54–56]. The raw ion counts were median-normalized for each metabolite, and differences in metabolite concentrations were determined by calculating the Log2 fold change for each biological replicate ($\Delta$*Slc38a1*/WT). The volcano plots were generated by taking the average of the biological replicates, and significance testing was performed using multiple paired two-tailed t-tests.

*EAE model with in vivo MeAIB treatment:* Female C57BL/6 mice aged 8 weeks were i.p. injected with either PBS or 3mg MeAIB in PBS daily from day 0 until end of study. All mice were injected subcutaneously with 0.2mL MOG/CFA emulsion (Hooke Laboratories) on day 7, and i.p. with 100ng PTX on days 7 and 8 to induce EAE. Mice were monitored and scored daily for clinical symptoms according to the following criteria: 0- no symptoms, 0.5- partial loss of tail tonicity, 1- complete loss of tail tonicity, 2- unilateral hind limb paresis, 2.5- unilateral hind limb paralysis, 3- partial bilateral hind limb paralysis, 3.5- complete bilateral hind limb paralysis (humane endpoint), 4- forelimb paresis, and 5- moribund/death. At the end of study, surviving mice were sacrificed for spleen, spinal cord, and brain collection for further analysis by flow cytometry. Tissues were dissociated, digested, and purified as described above.

*Allergic airway disease model with in vivo MeAIB treatment:* Parallel to the EAE model, female C57BL/6 mice aged 8 weeks were treated with daily injection of i.p. PBS or 3mg MeAIB in PBS from day 0 until end of study. On days 7, 14, and 21, mice were intranasally administered 50 µl PBS or 100 µg HDM Dermatophagoides pteronyssinus extract and 0.1 µg LPS from Escherichia coli 0111:B4 in 50 µl PBS. On day 22, mice were sacrificed by fatal dose of i.p. phenobarbital. BALF was collected by instilling and withdrawing 800µL of saline through a tracheostomy tube into the lungs. Cells from the BALF were then adhered to a slide, stained using Epredia Richard-Allan Scientific Three-Step Stain Kit, and identified as eosinophils, neutrophils, lymphocytes, and macrophages using light microscopy for quantification as previously described [57]. Lungs were dissociated to obtain single-cell suspensions as above for analysis by flow cytometry

*Th1-cell driven IBD colitis model with CRISPR/Cas9-mediated Slc38a1 knockout:* WT and Δ*Slc38a1 Th1* cells were prepared following *in vitro* single-gene knockout using CRISPR/Cas9 protocol using NTC gRNA#1, *Slc38a1* gRNA#1, and *Slc38a2* gRNA#1. On day 7 post T cell activation, transduced cells were isolated by CD90.1 positive selection and 400,000 viable cells were transferred into male *Rag1*$^{-/-}$ mice aged 8 weeks by i.p. injection to induce IBD colitis. Mice were weighed twice weekly to monitor disease progression and sacrificed at 7 weeks post transfer to collect spleen and mesenteric lymph nodes for T cell phenotyping by flow cytometry. Colons were also collected and fixed in 10% formalin for histology.

**Quantification and Statistical Analysis**

Statistical analyses were performed with Prism software (v10). Significance is indicated as follows: ns denotes $p>0.05$, * $p\leq0.05$, ** $p\leq0.01$, *** $p\leq0.001$, **** $p\leq0.0001$. For multiple comparisons corrections using FDR, nd denotes no discovery, * $q\geq Q$ where Q=5% unless otherwise specified. Error bars show mean ± standard deviation (SD) unless otherwise indicated for standard error of the mean (SEM). Sample sizes were chosen based on previous studies. Flow cytometric plots shown are representative of biological replicates.

**SUPPLEMENTAL ITEM TITLES**

- **Supplemental Figure 1:** CD4$^+$ T cells are dependent on uptake of select essential and non-essential AAs. **(**related to Figure 1**)**
- **Supplemental Figure 2:** *In vivo* CRISPR screening in Th1 cell- and Th17 cell-driven EAE show differential dependence on AA metabolism associated genes. **(**related to Figure 2)
- **Supplemental Figure 3:** Regulation of AA transporters and SLC38A1 dependence in CD4$^+$ T cells. **(**related to Figure 3)
- **Supplemental Figure 4:** Comparing SLC and metabolic pathway dependencies in WT Th1 and Th17 cells. **(**related to Figure 4)
- **Supplemental Figure 5:** Treatment with SLC38A1/2 inhibitor MeAIB have differential effects on CD4+ T cell subsets, and also reduces CD8+ proliferation. (related to Figure 7)
- **Supplemental Figure 6:** SLC38A1/2 inhibition by MeAIB differentially affects glycolytic and mitochondrial flux in Th1 and Th17 cells. **(**related to Figure 7)
- **Supplemental Figure 7:** SLC38A1/2 inhibition in vivo has no effect on Th17-cell driven allergic airway disease model as predicted by in vivo CRISPR screen. (related to Figure 7)
- **Supplemental Table 1**: SLC gRNA library **(**related to Figure 1)
- **Supplemental Table 2:** Glutamine metabolism gRNA library **(**related to Figure 2)
- **Supplemental Table 3:** Primers for CRISPR gRNA library prep
